# Ultra-rare genetic variation in the epilepsies: a whole-exome sequencing study of 17,606 individuals

**DOI:** 10.1101/525683

**Authors:** Epi25 Collaborative, Yen-Chen Anne Feng, Daniel P. Howrigan, Liam E. Abbott, Katherine Tashman, Felecia Cerrato, Tarjinder Singh, Henrike Heyne, Andrea Byrnes, Claire Churchhouse, Dennis Lal, Erin L. Heinzen, Gianpiero L. Cavalleri, Hakon Hakonarson, Ingo Helbig, Roland Krause, Patrick May, Sarah Weckhuysen, Slavé Petrovski, Sitharthan Kamalakaran, Sanjay M. Sisodiya, Patrick Cossette, Chris Cotsapas, Peter De Jonghe, Tracy Dixon-Salazar, Renzo Guerrini, Patrick Kwan, Anthony G. Marson, Randy Stewart, Chantal Depondt, Dennis J. Dlugos, Ingrid E. Scheffer, Pasquale Striano, Catharine Freyer, Kevin McKenna, Brigid M. Regan, Susannah T. Bellows, Costin Leu, Caitlin A. Bennett, Esther M.C. Johns, Alexandra Macdonald, Hannah Shilling, Rosemary Burgess, Dorien Weckhuysen, Melanie Bahlo, Terence J. O’Brien, Marian Todaro, Hannah Stamberger, Danielle M. Andrade, Tara R. Sadoway, Kelly Mo, Heinz Krestel, Sabina Gallati, Savvas S. Papacostas, Ioanna Kousiappa, George A. Tanteles, Katalin Štěrbová, Markéta Vlčková, Lucie Sedláčková, Petra Laššuthová, Karl Martin Klein, Felix Rosenow, Philipp S. Reif, Susanne Knake, Wolfram S. Kunz, Gábor Zsurka, Christian E. Elger, Jürgen Bauer, Michael Rademacher, Manuela Pendziwiat, Hiltrud Muhle, Annika Rademacher, Andreas van Baalen, Sarah von Spiczak, Ulrich Stephani, Zaid Afawi, Amos D. Korczyn, Moien Kanaan, Christina Canavati, Gerhard Kurlemann, Karen Müller-Schlüter, Gerhard Kluger, Martin Häusler, Ilan Blatt, Johannes R. Lemke, Ilona Krey, Yvonne G. Weber, Stefan Wolking, Felicitas Becker, Christian Hengsbach, Sarah Rau, Ana F. Maisch, Bernhard J. Steinhoff, Andreas Schulze-Bonhage, Susanne Schubert-Bast, Herbert Schreiber, Ingo Borggräfe, Christoph J. Schankin, Thomas Mayer, Rudolf Korinthenberg, Knut Brockmann, Gerhard Kurlemann, Dieter Dennig, Rene Madeleyn, Reetta Kälviäinen, Pia Auvinen, Anni Saarela, Tarja Linnankivi, Anna-Elina Lehesjoki, Mark I. Rees, Seo-Kyung Chung, William O. Pickrell, Robert Powell, Natascha Schneider, Simona Balestrini, Sara Zagaglia, Vera Braatz, Michael R. Johnson, Pauls Auce, Graeme J. Sills, Larry W. Baum, Pak C. Sham, Stacey S. Cherny, Colin H.T. Lui, Nina Barišić, Norman Delanty, Colin P. Doherty, Arif Shukralla, Mark McCormack, Hany El-Naggar, Laura Canafoglia, Silvana Franceschetti, Barbara Castellotti, Tiziana Granata, Federico Zara, Michele Iacomino, Francesca Madia, Maria Stella Vari, Maria Margherita Mancardi, Vincenzo Salpietro, Francesca Bisulli, Paolo Tinuper, Laura Licchetta, Tommaso Pippucci, Carlotta Stipa, Raffaella Minardi, Antonio Gambardella, Angelo Labate, Grazia Annesi, Lorella Manna, Monica Gagliardi, Elena Parrini, Davide Mei, Annalisa Vetro, Claudia Bianchini, Martino Montomoli, Viola Doccini, Carla Marini, Toshimitsu Suzuki, Yushi Inoue, Kazuhiro Yamakawa, Birute Tumiene, Lynette G. Sadleir, Chontelle King, Emily Mountier, S. Hande Caglayan, Mutluay Arslan, Zuhal Yapıcı, Uluc Yis, Pınar Topaloglu, Bulent Kara, Dilsad Turkdogan, Aslı Gundogdu-Eken, Nerses Bebek, Sibel Uğur-İşeri, Betül Baykan, Barış Salman, Garen Haryanyan, Emrah Yücesan, Yeşim Kesim, Çiğdem Özkara, Annapurna Poduri, Russell J. Buono, Thomas N. Ferraro, Michael R. Sperling, Warren Lo, Michael Privitera, Jacqueline A. French, Steven Schachter, Ruben I. Kuzniecky, Orrin Devinsky, Manu Hegde, Pouya Khankhanian, Katherine L. Helbig, Colin A. Ellis, Gianfranco Spalletta, Fabrizio Piras, Federica Piras, Tommaso Gili, Valentina Ciullo, Andreas Reif, Andrew McQuillin, Nick Bass, Andrew McIntosh, Douglas Blackwood, Mandy Johnstone, Aarno Palotie, Michele T. Pato, Carlos N. Pato, Evelyn J. Bromet, Celia Barreto Carvalho, Eric D. Achtyes, Maria Helena Azevedo, Roman Kotov, Douglas S. Lehrer, Dolores Malaspina, Stephen R. Marder, Helena Medeiros, Christopher P. Morley, Diana O. Perkins, Janet L. Sobell, Peter F. Buckley, Fabio Macciardi, Mark H. Rapaport, James A. Knowles, Cohort (GPC) Consortium Genomic Psychiatry, Ayman H. Fanous, Steven A. McCarroll, Namrata Gupta, Stacey B. Gabriel, Mark J. Daly, Eric S. Lander, Daniel H. Lowenstein, David B. Goldstein, Holger Lerche, Samuel F. Berkovic, Benjamin M. Neale

## Abstract

Sequencing-based studies have identified novel risk genes for rare, severe epilepsies and revealed a role of rare deleterious variation in common epilepsies. To identify the shared and distinct ultra-rare genetic risk factors for rare and common epilepsies, we performed a whole-exome sequencing (WES) analysis of 9,170 epilepsy-affected individuals and 8,364 controls of European ancestry. We focused on three phenotypic groups; the rare but severe developmental and epileptic encephalopathies (DEE), and the commoner phenotypes of genetic generalized epilepsy (GGE) and non-acquired focal epilepsy (NAFE). We observed that compared to controls, individuals with any type of epilepsy carried an excess of ultra-rare, deleterious variants in constrained genes and in genes previously associated with epilepsy, with the strongest enrichment seen in DEE and the least in NAFE. Moreover, we found that inhibitory GABA_A_ receptor genes were enriched for missense variants across all three classes of epilepsy, while no enrichment was seen in excitatory receptor genes. The larger gene groups for the GABAergic pathway or cation channels also showed a significant mutational burden in DEE and GGE. Although no single gene surpassed exome-wide significance among individuals with GGE or NAFE, highly constrained genes and genes encoding ion channels were among the top associations, including *CACNA1G, EEF1A2*, and *GABRG2* for GGE and *LGI1, TRIM3*, and *GABRG2* for NAFE. Our study confirms a convergence in the genetics of common and rare epilepsies associated with ultra-rare coding variation and highlights a ubiquitous role for GABAergic inhibition in epilepsy etiology in the largest epilepsy WES study to date.

## Introduction

Epilepsy is a group of disorders characterized by repeated seizures due to excessive electrical activity in the brain, one of the most common and burdensome neurological conditions worldwide^1; 2^. A core challenge for epilepsy genetics is identifying and disentangling the genetic architecture and biological mechanisms underlying the variety of epilepsy types (e.g., focal vs. generalized) and electroclinical syndromes. While the occurrence of epilepsy for many affected individuals carries an underlying genetic component^3-5^, the highly heterogeneous nature of epileptic seizures, epilepsy types, severity, and comorbidity makes it difficult to determine the specific genetic risks for each patient. For individuals with common, complex types of epilepsy, where inheritance may be due to strongly acting mutations, oligogenic or polygenic, the discovery of genetic risk factors is particularly challenging.

Considerable progress in our understanding of the genetic risk factors for epilepsy has been made in recent years thanks to the rapid growth and advancement in sequencing technology. Dozens of epilepsy-causing genes have been identified in individuals diagnosed with severe epilepsy syndromes^6-10^, known as the developmental and epileptic encephalopathies (DEE). DEE are rare in the population and typically begin early in life. Incidence of the entire group is not well established, but a recent epidemiological study of severe epilepsies limited to onset under 18 months found an incidence of 1 in 2000 births^11^. The incidence of Dravet syndrome, one of the important specific forms of DEE, has been shown in several studies as 1 in 22,000^10^. Individuals with DEE usually have developmental impairment, ranging from profound to mild. With such severity, sequencing-based studies continue to discover *de novo* pathogenic variants for DEE and have implicated genes encoding neuronal ion channels and receptors and genes involved in cellular signaling^6-10; 12^. The common subgroups of epilepsy, broadly comprising genetic generalized epilepsy (GGE) and non-acquired focal epilepsy (NAFE), account for a major proportion of all incident epilepsies^2; 13^ and have been shown robust heritability in twin, family, and genome-wide association studies (GWAS)^4;14-16^. Disappointingly, only a limited number of genes had been discovered to date for the common epilepsies, mostly from rare monogenic families with focal epilepsies, and attempts to identify clear risk genes for GGE have been least successful^12;17;18^. In most cases, especially for non-familial onset, the specific pathogenic variants are not yet known, and gene findings from small-scale studies have often not been reproducible^19-21^.

Current evidence of the genetic etiology of epilepsies has revealed both extensive phenotypic and genetic heterogeneity. Many of the identified genes are associated with a spectrum of mild to severe epilepsies, showing phenotypic pleiotropy or variable expressivity, and most of the electroclinical syndromes have diverse genetic causes^10; 22^. Two recent studies using whole-exome sequencing (WES) of hundreds of individuals with common familial epilepsies found an enrichment in ultra-rare genetic variation in genes associated with rare epilepsy syndromes^17^ and in missense variants in a group of genes encoding all GABA_A_ receptors, the most important neurotransmitter receptors for neuronal inhibition in the mammalian brain^18^. Given the complex genetic architecture of the epilepsies, it is therefore critical to pinpoint the distinct and overlapping genetic risk factors underlying different groups of epilepsy on a scale much larger than previous sequencing studies and beyond familial cases.

Here, we evaluate a WES case-control study of epilepsy from the Epi25 collaborative— an ongoing global effort that collected an unprecedented number of patient cohorts for primarily the three major classes of non-lesional epilepsies: DEE, GGE, and NAFE^22^. We aimed to characterize the genetic risk of ultra-rare coding variants across these common and rare epilepsy subgroups by evaluating the burden at the individual gene level and in candidate gene sets to understand the role of rare genetic variation in epilepsy and identify specific epilepsy risk genes.

## Subjects and Methods

### Study design and participants

We collected DNA and detailed phenotyping data on individuals with epilepsy from 37 sites in Europe, North America, Australasia and Asia (**Supplemental Subjects and Methods; Table S1**). Here we analyzed subjects with genetic generalized epilepsy (GGE, also known as idiopathic generalized epilepsy; N=4,453), non-acquired focal epilepsy (NAFE; N=5,331) and developmental and epileptic encephalopathies (DEE; N=1,476); and a small number of other epilepsies were also included in the initiative (**Table S1**).

Control samples were aggregated from local collections at the Broad Institute (Cambridge, MA, USA) or obtained from dbGaP, consisting of 17,669 individuals of primarily European ancestry who were not ascertained for neurological or neuropsychiatric conditions (**Table S2; Supplemental Subjects and Methods**).

### Phenotyping procedures

Epilepsies were diagnosed on clinical grounds based on criteria given in the next paragraph (see below for GGE, NAFE and DEE, respectively) by experienced epileptologists and consistent with International League Against Epilepsy (ILAE) classification at the time of diagnosis and recruitment. De-identified (non-PHI [protected health information]) phenotyping data were entered into the Epi25 Data repository hosted at the Luxembourg Centre for Systems Biomedicine via detailed on-line case record forms based on the RedCAP platform. Where subjects were part of previous coordinated efforts with phenotyping on databases (e.g., the Epilepsy Phenome/Genome Project^23^ and the EpiPGX project (www.epipgx.eu)), deidentified data were accessed and transferred to the new platform. Phenotyping data underwent review for uniformity among sites and quality control by automated data checking, followed by manual review if required. Where doubt remained about eligibility, cases were reviewed by the phenotyping committee and sometimes further data was requested from the source site before a decision was made.

### Case Definitions

GGE required a convincing history of generalized seizure types (generalized tonic-clonic seizures, absence, or myoclonus) and generalized epileptiform discharges on EEG. We excluded cases with evidence of focal seizures, or with moderate to severe intellectual disability and those with an epileptogenic lesion on neuroimaging (although neuroimaging was not obligatory). If a diagnostic source EEG was not available, then only cases with an archetypal clinical history as judged by the phenotyping committee (e.g., morning myoclonus and generalized tonic-clonic seizures for a diagnosis of Juvenile Myoclonic Epilepsy) were accepted.

Diagnosis of NAFE required a convincing history of focal seizures, an EEG with focal epileptiform or normal findings (since routine EEGs are often normal in focal epilepsy), and neuroimaging showing no epileptogenic lesion except hippocampal sclerosis (MRI was preferred but CT was accepted). Exclusion criteria were a history of generalized onset seizures or moderate to severe intellectual disability.

The DEE group comprised subjects with severe refractory epilepsy of unknown etiology with developmental plateau or regression, no epileptogenic lesion on MRI, and with epileptiform features on EEG. As this is the group with the largest number of gene discoveries to date, we encouraged inclusion of those with non-explanatory epilepsy gene panel results, but we did not exclude those without prior testing (**Table S7**).

### Informed Consent

Patients or their legal guardians provided signed informed consent according to local national ethical requirements. Samples had been collected over a 20-year period in some centers, so the consent forms reflected standards at the time of collection. Samples were only accepted if the consent did not exclude data sharing. For samples collected after January 25, 2015, consent forms required specific language according to the NIH Genomic Data Sharing policy (http://gds.nih.gov/03policy2.html).

### Whole exome sequencing data generation

All samples were sequenced at the Broad Institute of Harvard and MIT on the Illumina HiSeq X platform, with the use of 151 bp paired-end reads. Exome capture was performed with Illumina Nextera® Rapid Capture Exomes or TruSeq Rapid Exome enrichment kit (target size 38 Mb), except for three control cohorts (MIGen ATVB, MIGen Ottawa, and Swedish SCZ controls) for which the Agilent SureSelect Human All Exon Kit was used (target size 28.6 Mb – 33 Mb). Sequence data in the form of BAM files were generated using the Picard data-processing pipeline and contained well-calibrated reads aligned to the GRCh37 human genome reference. Samples across projects were then jointly called via the Genome Analysis Toolkit (GATK) best practice pipeline^24^ for data harmonization and variant discovery. This pipeline detected single nucleotide (SNV) and small insertion/deletion (indel) variants from exome sequence data.

### Quality control

Variants were pre-filtered to keep only those passing the GATK VQSR (Variant Quality Score Recalibration) metric and those lying outside of low complexity regions^25^. Genotypes with GQ < 20 and heterozygous genotype calls with allele balance > 0.8 or < 0.2 were set to missing. To control for capture platform difference, we retained variants that resided in GENCODE coding regions where 80% of Agilent and Illumina-sequenced samples show at least 10x coverage. This resulted in the removal of ∼50% of the called sites (23% of the total coding variants and 97% of the total non-coding variants) but effectively reduced the call rate difference between cases and controls (**Figure S1**). To further identify potential false positive sites due to technical variation, we performed single variant association tests (for variants with a minor allele frequency MAF > 0.001) among the controls, treating one platform as the pseudo-case group with adjustment for sex and the first ten principal components (PCs), and removed variants significantly associated with capture labels (*p*-value < 0.05). We also excluded variants with a call rate < 0.98, case-control call rate difference > 0.005, or Hardy-Weinberg Equilibrium (HWE) test *p*-value < 1×10^-6^ based on the combined case and control cohort.

Samples were excluded if they had a low average call rate (< 0.98), low mean sequence depth (< 30; **Figure S2**), low mean genotype quality (< 85), high freemix contamination estimate (> 0.04), or high percent chimeric reads (> 1.4%). We performed a series of principal component analyses (PCAs) to identify ancestral backgrounds and control for population stratification, keeping only individuals of European (EUR) ancestry classified by Random Forest with 1000 Genomes data (**Figure S3**). Within the EUR population, we removed controls not well-matched with cases based on the top two PCs, and individuals with an excessive or a low count of synonymous singletons—a number that increases with the North-to-South axis (**Figure S4**). We also removed one sample from each pair of related individuals (proportion identity-by-descent > 0.2) and those whose genetically imputed sex was ambiguous or did not match with self-reported sex. Outliers (>4SD from the mean) of transition/transversion ratio, heterozygous/homozygous ratio, or insertion/deletion ratio within each cohort were further discarded (**Figures S5-7**). At the phenotype level, we removed individuals with epilepsy phenotype to-be-determined or marked as “excluded” from further review.

The number of variant and sample dropouts at each step are detailed in **Tables S3** and **S4.**

### Variant annotation

Annotation of variants was performed with Ensembl’s Variant Effect Predictor (VEP)^26^ for human genome assemble GRCh37. Based on the most severe consequence, we defined four mutually exclusive functional classes of variants using relevant terms and SnpEff^27^ impact (**Table S5**): protein-truncating variant (PTV), damaging missense (predicted by PolyPhen-2 and SIFT), other/benign missense (predicted by PolyPhen-2 and SIFT), and synonymous. To further discriminate likely deleterious missense variants from benign missense variants, we applied an *in silico* missense deleteriousness predictor (“Missense badness, PolyPhen-2, and regional Constraint”, or MPC score)^28^ that leverages regional constraint information to annotate a subset of missense variants that are highly deleterious (MPC ≥ 2). The MPC ≥ 2 group accounts for a small proportion of the total damaging and benign missense variants annotated by PolyPhen-2 and SIFT. Because many of our control samples were obtained from external datasets used in the Exome Aggregation Consortium (ExAC)^29^ (**Table S2)**, we used the DiscovEHR cohort—an external population allele frequency reference cohort that contains 50,726 whole-exome sequences from a largely European and non-diseased adult population^30^—to annotate if a variant is absent in the general population (**Figure S8**).

### Gene-set burden analysis

To estimate the excess of rare, deleterious protein-coding variants in individuals with epilepsy, we conducted burden tests across the entire exome, for biologically relevant gene sets and at the individual gene level. We focused on two definitions of “ultra-rare” genetic variation (URV) for the primary analyses—variants not seen in the DiscovEHR database and observed only once among the combined case and control test cohort (allele count AC=1) or absent in DiscovEHR and observed no more than three times in the test cohort (allele count AC≤3)—where the strongest burden of deleterious pathogenic variants have been observed previously^17; 31^ and in our study compared to less stringent allele frequency thresholds (**Figure S9 & S10**). We performed these case-control comparisons separately for each of the three primary epilepsy disorders (DEE, GGE, NAFE) and again for all epilepsy-affected individuals combined.

Gene-set burden tests were implemented using logistic regression to examine the enrichment of URVs in individuals with epilepsy versus controls. We performed the test by regressing case-control status on certain classes of URVs aggregated across a target gene set in an individual, adjusting for sex, the top ten PCs, and exome-wide variant count. This analysis tested the burden of URVs separately for five functional coding annotations: synonymous, benign missense predicted by PolyPhen-2 and SIFT, damaging missense predicted by PolyPhen-2 and SIFT, protein-truncating variants, and missense with MPC≥2 (**Table S5**). To help determine whether our burden model was well calibrated, we used synonymous substitutions as a negative control, where significant burden effects would more likely indicate insufficient control of population stratification or exome capture differences. The inclusion of overall variant count as a covariate—which tracks with ancestry—made our test conservative but allows for better control of residual population stratification not captured by PCs, and effectively reduces inflation of signals in synonymous variants (**Figure S11**). We collected and tested eleven different gene sets, including constrained genes, brain-enriched genes, and genes reported to be associated with epilepsy or epilepsy-related mechanisms^9; 17;18; 32; 33^ (**Table S6**). Unlike the gene-based burden tests, because most of the gene-set tests were not independent, we used a false discovery rate (FDR) correction for multiple testing that accounted for the number of functional categories (5), gene sets (11) and epilepsy phenotypes (4), totaling 220 tests, and defined a significant enrichment at FDR < 0.05.

### Gene-based collapsing analysis

For gene-based tests, we restricted to deleterious URVs annotated as either PTV, missense with MPC≥2, or in-frame insertion/deletion. For each gene, individuals who had at least one copy of these deleterious variants were counted as a carrier, and we used a two-tailed Fisher’s Exact test (FET) to assess if the proportion of carriers among epilepsy subgroup cases was significantly higher than controls. Instead of assuming a uniform distribution for p-values under the null, we generated empirical p-values by permuting case-control labels 500 times, ordering the FET p-values of all genes for each permutation, and taking the average across all permutations to form a rank-ordered estimate of the expected p-value distribution. This was done by modifying functions in the “QQperm” R package^34^. To avoid potential false discoveries, we defined a stringent exome-wide significance as p-value < 6.8e-07, using Bonferroni correction to account for 18,509 consensus coding sequence genes tested and the four individual case-control comparisons.

Considering that recessive pathogenic variants were implicated in a number of epilepsy-associated genes, mostly identified from individuals with a DEE phenotype^7^, we conducted a secondary gene-based Fisher’s exact test using a recessive model, comparing the proportion of carriers that are homozygous for the minor allele between cases and controls. The recessive model was assessed for PTVs, missense (MPC≥2) variants, and in-frame indels separately. For this analysis, we did not restrict to non-DiscovEHR variants and relaxed the allele frequency up to MAF < 0.01 to account for the sparse occurrences.

Additionally, to evaluate the contribution of low frequency deleterious variants to epilepsy risk, we explored the gene burden of all protein-truncating and damaging missense variants for those with a MAF < 0.01 using SKAT^35^, including sex and the top ten PCs as covariates in the analysis. We performed the tests with the default weighting scheme (dbeta(1,25)).

### Single variant association

Associations of common and low-frequency variants (MAF > 0.001) with epilepsy were estimated using logistic regression by Firth’s method, correcting for sex and the first ten PCs.

Quality control, annotation, and analysis were largely performed using Hail^36^, an open-source software for scalable genomic data analysis, in conjunction with R (version 3.4.2).

## Results

### Whole exome sequencing, quality control, and sample overview

We performed WES on an initial dataset of over 30,000 epilepsy affected and control individuals. After stringent quality control (QC), we identified a total of 9,170 individuals with epilepsy and 8,436 controls without reported neurological or neuropsychiatric-related conditions, all of whom were unrelated individuals of European descent. Among the individuals with epilepsy, 1,021 were diagnosed with DEE, 3,108 with GGE, 3,597 with NAFE, and 1,444 with other epilepsy syndromes (lesional focal epilepsy, febrile seizures, and others). Cases and controls were carefully matched on genetic ancestry to eliminate the possibility of false positive findings induced by population stratification. Due to the lack of cosmopolitan controls from non-European populations, cases identified from PCA with a non-European ancestry were removed. Furthermore, to ensure the distribution of rare variants was balanced between cases and controls^37^, we removed a subset of case and control-only cohorts (from Sweden, Finland, Cyprus, and Turkey) where the mean synonymous singleton count that significantly deviated from the overall average being the consequence of incomplete ancestry matching (**Figure S4**). We called a total of 1,844,644 sites in 18,509 genes in the final dataset, comprising 1,811,325 SNVs and 33,319 indels, 48.5% of which were absent in the DiscovEHR database^30^. Among the non-DiscovEHR sites, 85% were singletons (defined as only one instance of that variant), and 99% had a minor allele count (AC) not more than three (equivalent to MAF ≤0.01%; **Figure S8**); the missense with MPC≥2 annotation accounted for 2.0% of the total missense variants (5.5% of the damaging and 1.0% of the benign missense variants predicted by PolyPhen-2 and SIFT). In our primary burden analyses, we focused on the “ultra-rare” non-DiscovEHR variants (URVs) that are unique to the 17,606 individuals under study and are seen either only once (AC=1) or no more than three times (AC≤3) in our dataset. These URVs were shown to confer the largest risk of epilepsy compared to singletons observed in DiscovEHR, doubletons, or beyond (**Figure S9 & S10**). As previously described, epilepsy enrichment signals diminished with an increase in allele frequency^17^.

### Enrichment of ultra-rare deleterious variants in constrained genes in DEE and GGE

We first tested the burden of singleton URVs for each epilepsy subgroup, as well as for all epilepsy-affected individuals combined, versus controls among gene sets collected based on current understanding and hypothesis of epilepsy causation (**Table S6**). To evaluate the burden in constrained genes, we defined “loss-of-function (LoF) intolerant” genes with either a pLI score^29^ > 0.9 (3,488 genes) or separately a pLI score > 0.995 (1,583 genes) and those as “missense-constrained” for genes with a missense Z-score > 3.09 (1,730 genes)^33^. We used a version of the scores derived from the non-neuropsychiatric subset of the Exome Aggregation Consortium (ExAC) samples. Because some of our control cohorts are also in ExAC (**Table S2**), we restricted our constrained gene burden tests to controls outside of the ExAC cohort (N=4,042).

We found that, consistent with a recent study that evaluated *de novo* burden in autism^38^, burden signals of PTVs were mostly contained in genes with a pLI > 0.995 compared to pLI > 0.9 (**Figure S12** & **S13**). Focusing on pLI > 0.995 in the all-epilepsy case-control analysis, both protein-truncating and damaging missense (MPC^28^≥2) URVs in LoF-intolerant genes showed a mutational burden with an odds ratio of 1.3 (*adjP* = 1.6×10^-4^) and 1.1 (*adjP* = 0.039), respectively. Breaking this down by epilepsy types, there was a significant excess of these deleterious URVs among individuals with DEE (*OR*_*PTV*_ = 1.4, *adjP*_*PTV*_ = 0.013; *OR*_*MPC*_ = 1.2, *adjP*_*MPC*_ = 0.019), as expected. This enrichment was also seen in individuals with GGE with a magnitude comparable to that in DEE (*OR*_*PTV*_ = 1.4, *adjP*_*PTV*_ = 9.1×10^-5^; *OR*_*MPC*_ = 1.2, *adjP*_*MPC*_ = 5.5×10^-3^), but was not significant in individuals with NAFE (*OR*_*PTV*_ = 1.2, *adjP*_*PTV*_ = 0.062; *OR*_*MPC*_ =1.0, *adjP*_*MPC*_ = 0.37; **Figure 1**). There was no evidence of excess burden in synonymous URVs, suggesting that enrichment of deleterious pathogenic variants was unlikely to be the result of un-modeled population stratification or technical artifact. Among *in-silico* missense predictors, MPC≥2 annotations consistently showed a higher burden than those predicted by PolyPhen-2 and SIFT. The burden among missense-constrained genes exhibited a similar pattern, with PTVs showing a higher burden in DEE than in the common epilepsy types (**Figure S14**). In addition, both large gene sets were more enriched for PTVs than for damaging missense variants.

**Figure 1.**
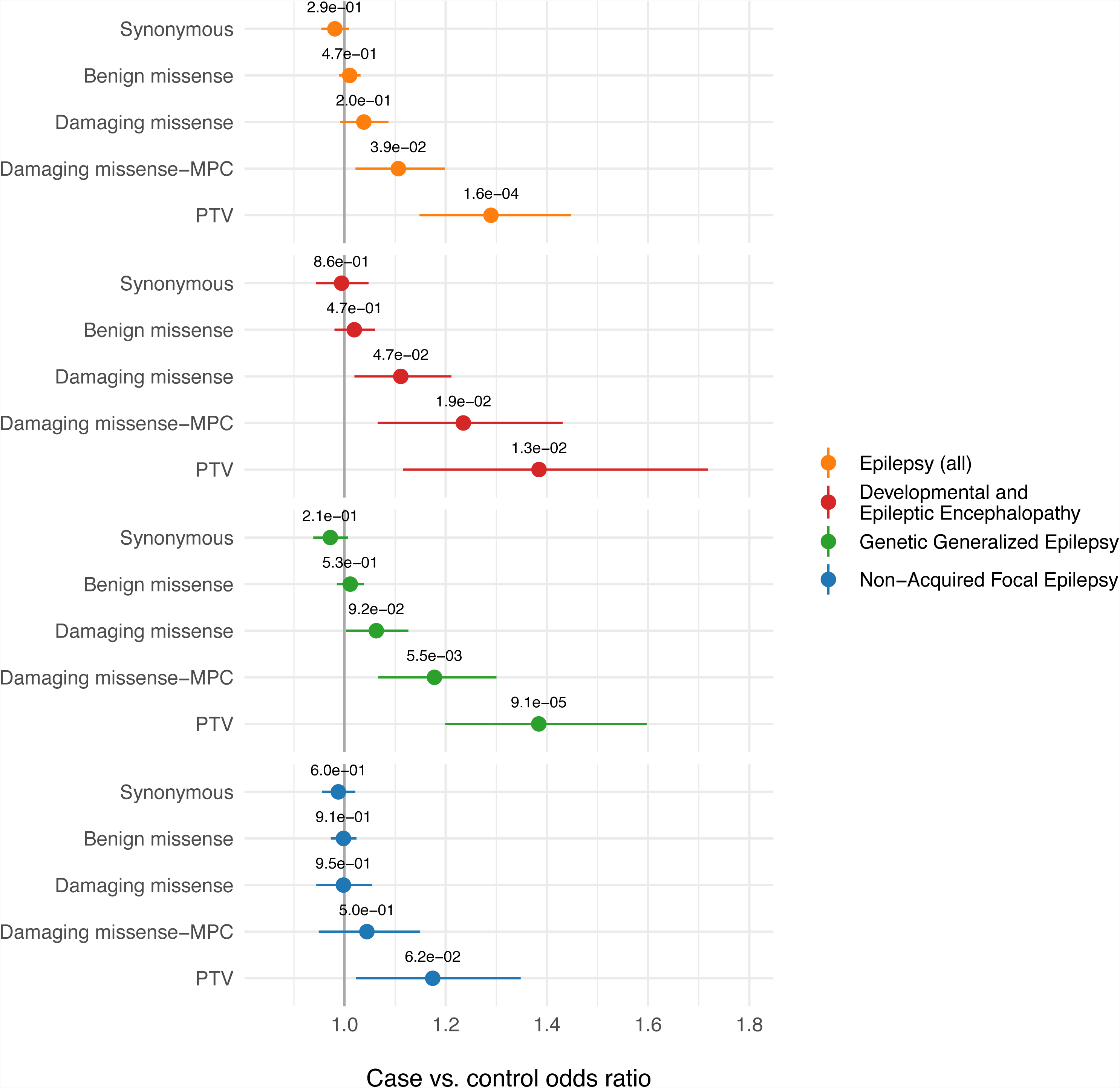
Burden of ultra-rare singletons in LoF-intolerant genes (pLI > 0.995) This analysis was restricted to 4,042 non-ExAC controls for comparison with epilepsy cases. We focused on “ultra-rare” variants not observed in the DiscovEHR database. Significance of association was displayed in FDR-adjusted p-values; odds ratios and 95% CIs were not multiplicity adjusted. The five functional coding annotations were defined as described in **Table S5**. PTV denotes protein-truncating variants; the “damaging missense” and “benign missense” categories were predicted by PolyPhen-2 and SIFT, while “damaging missense-MPC” was a group of missense variants with a missense badness score (MPC) ≥ 2. From top to bottom are the results based on all-epilepsy, DEE, GGE, and NAFE. Epilepsy cases, except for individuals with NAFE, carried a significant excess of ultra-rare PTV and damaging missense (MPC≥2) variants compared to controls (FDR < 0.05). PTV burden was higher than missense (MPC≥2) burden across epilepsy types.

### Burden in candidate genetic etiologies associated with epilepsy

Among URVs in previously reported epilepsy genes, we found an expected and pronounced difference in the number of singleton protein-truncating URVs in individuals with DEE relative to controls. PTVs were associated with an increased DEE risk in 43 known dominant epilepsy genes^17^ (*OR* = 6.3, *adjP* = 2.1×10^-08^), 50 known dominant DEE genes^9^ (*OR* = 9.1, *adjP* = 7.8×10^-11^), and 33 genes with *de novo* burden in neurodevelopmental disorders with epilepsy^9^ (*OR* = 14.8, *adjP* =1.7×10^-12^). Evidence for an excess of ultra-rare PTVs was also observed in individuals with GGE, with an odds ratio ranging from 2 to 4. No enrichment of PTVs was observed among people with NAFE (**Figure 2A; Table S8**). In contrast, the burden of singleton missense (MPC≥2) URVs was more pervasive across epilepsy types. Compared to controls, there was a 3.6-fold higher rate of these missense URVs in established epilepsy genes in individuals with DEE (*adjP* = 1.6×10^-10^), a 2.3-fold elevation in individuals with GGE (*adjP* = 6.4×10^-07^), and a 1.9-fold elevation in individuals with NAFE (*adjP* = 2.8×10^-4^).

**Figure 2.**
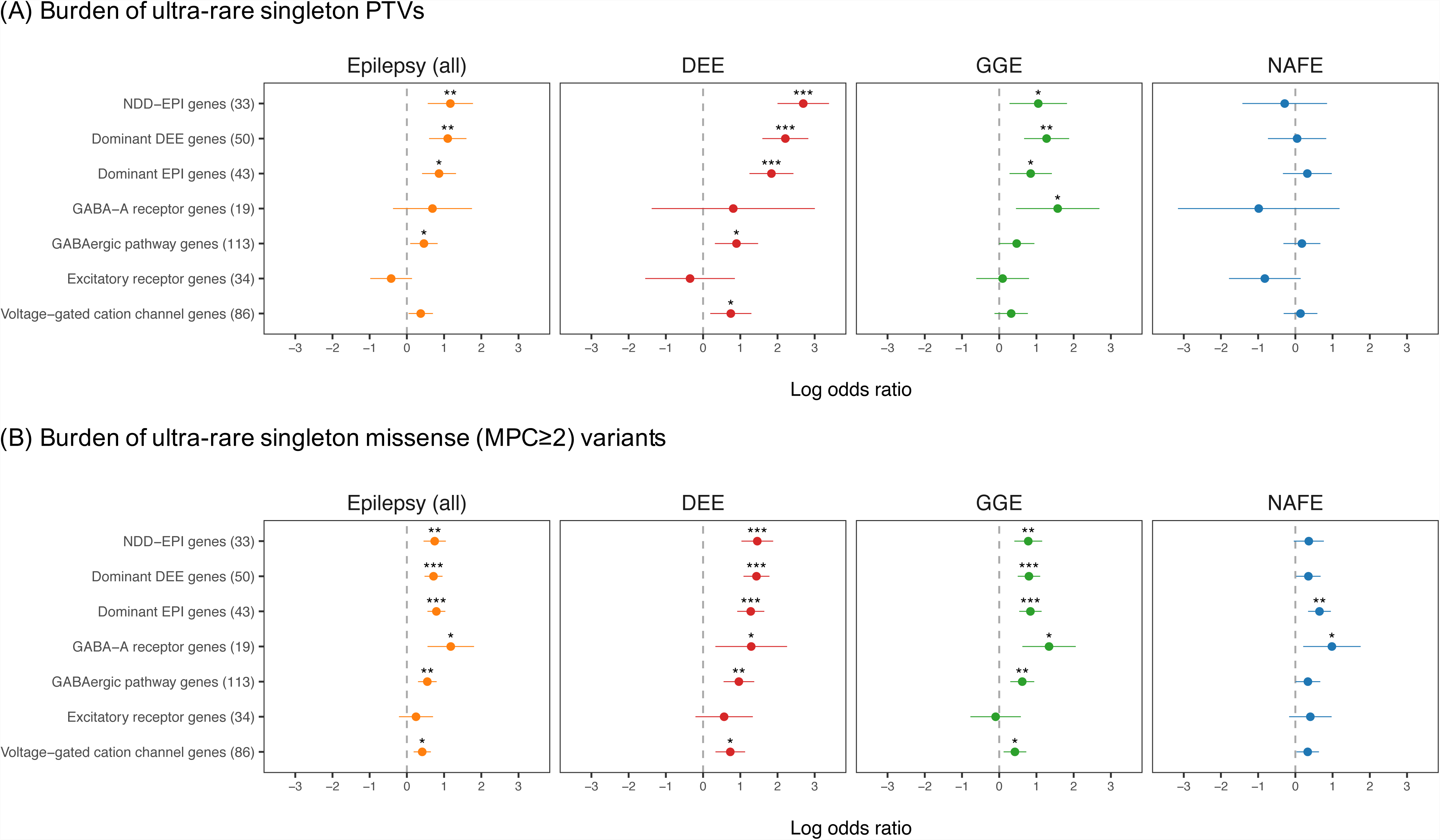
Burden of ultra-rare singletons annotated as (A) protein-truncating variants or (B) damaging missense (MPC≥2) variants. “Ultra-rare” variants (URVs) were defined as not observed in the DiscovEHR database. Gene sets were defined in **Table S6**, with the number of genes specified in the parenthesis. DEE stands for individuals with developmental and epileptic encephalopathies, GGE for genetic generalized epilepsy, NAFE for non-acquired focal epilepsy, and EPI for all epilepsy; NDD-EPI genes are genes with *de novo* burden in neurodevelopmental disorders with epilepsy. Star signs indicate significance after FDR control (“*”: FDR-adjusted p-value < 0.05; “**”: adjusted p-value < 1×10^-3^; “***”: adjusted p-value < 1×10^-5^). PTVs were enriched in candidate epilepsy genes for individuals with DEE relative to other epilepsy subgroups, but did not show a strong signal in inhibitory, excitatory receptors or voltage-gated cation channel genes. The burden of damaging missense (MPC≥2) variants, on the other hand, was stronger across these gene sets compared to PTVs, especially for GABA_A_ receptor genes and genes involved in GABAergic pathways. Relative to other epilepsy types, individuals with NAFE consistently showed the least burden of deleterious URVs. No enrichment was observed from excitatory receptors.

### Burden in genes encoding for cation channels and neurotransmitter receptors

Among brain-enriched genes—those defined as genes with at least a 2-fold increase in expression in brain tissues relative to their average expression across tissues based on GTEx data^32^—both protein-truncating and damaging missense (MPC≥2) URVs were significantly enriched in epilepsy cases versus controls, and the missense burden was much higher than the PTV burden (**Figure S15**). We then investigated the burden in four smaller gene sets previously implicated as mechanisms driving the etiology of epilepsy; these included 19 genes encoding GABA_A_ receptor subunits, 113 genes involved in GABAergic pathways, 34 genes encoding excitatory receptors (ionotropic glutamate receptor subunits and nicotinic acetylcholine receptor subunits), and 86 voltage-gated cation channel genes (e.g., sodium, potassium, calcium—full list in **Table S6**)^18^. We discovered that, relative to damaging missense variants, the distribution of PTVs in most of these gene sets did not differ significantly between epilepsy cases and controls (**Figure 2A**; **Table 1**). The PTV signals that remained significant after FDR correction included, for individuals with DEE, an increased burden in GABAergic pathway genes and voltage-gated cation channels, and noticeably, for individuals with GGE, an increased burden in the inhibitory GABA_A_ receptors (*OR* = 4.8, *adjP* = 0.021). No PTV burden was detected for individuals with NAFE. In contrast, the enrichment of missense (MPC≥2) URVs was more extensive in these gene sets across all epilepsy-control comparisons (**Figure 2A**; **Table 1**). The burden of these damaging missense pathogenic variants was seen in GABA_A_ receptor genes (*OR*_*DEE*_ = 3.7, *adjP*_*DEE*_ = 0.028; *OR*_*GGE*_ = 3.8, *adjP*_*GGE*_ = 1.4×10^-3^; *OR*_*NAFE*_ = 2.7, *adjP*_*NAFE*_ = 0.039), GABAergic pathway genes (*OR*_*DEE*_ = 2.6, *adjP*_*DEE*_ = 4.7×10^-5^; *OR*_*GGE*_ = 1.9, *adjP*_*GGE*_ = 9.9×10^-04^; *OR*_*NAFE*_ = 1.4, adjP_NAFE_= 0.11), and voltage-gated cation channel genes (*OR*_*DEE*_ = 2.1, *adjP*_*DEE*_ = 1.7×10^-03^; *OR*_*GGE*_ = 1.5, *adjP*_*GGE*_ = 0.023; *OR*_*NAFE*_ = 1.4, *adjP*_*NAFE*_ = 0.081). However, no enrichment was detected in genes encoding excitatory receptors. For individuals with NAFE, the burden signals were consistently the weakest across gene sets compared to the other epilepsy phenotypes. None of the gene sets was enriched for putatively neutral variation, except for a slightly elevated synonymous burden in GABA_A_ receptor genes (**Table S8**). These results support a recent finding where rare missense variation in GABA_A_ receptor genes conferred a significant risk to GGE^18^, and together implicate the relative importance and involvement of damaging missense variants in abnormal inhibitory neurotransmission in both rare and common epilepsy types.

**Table 1.**
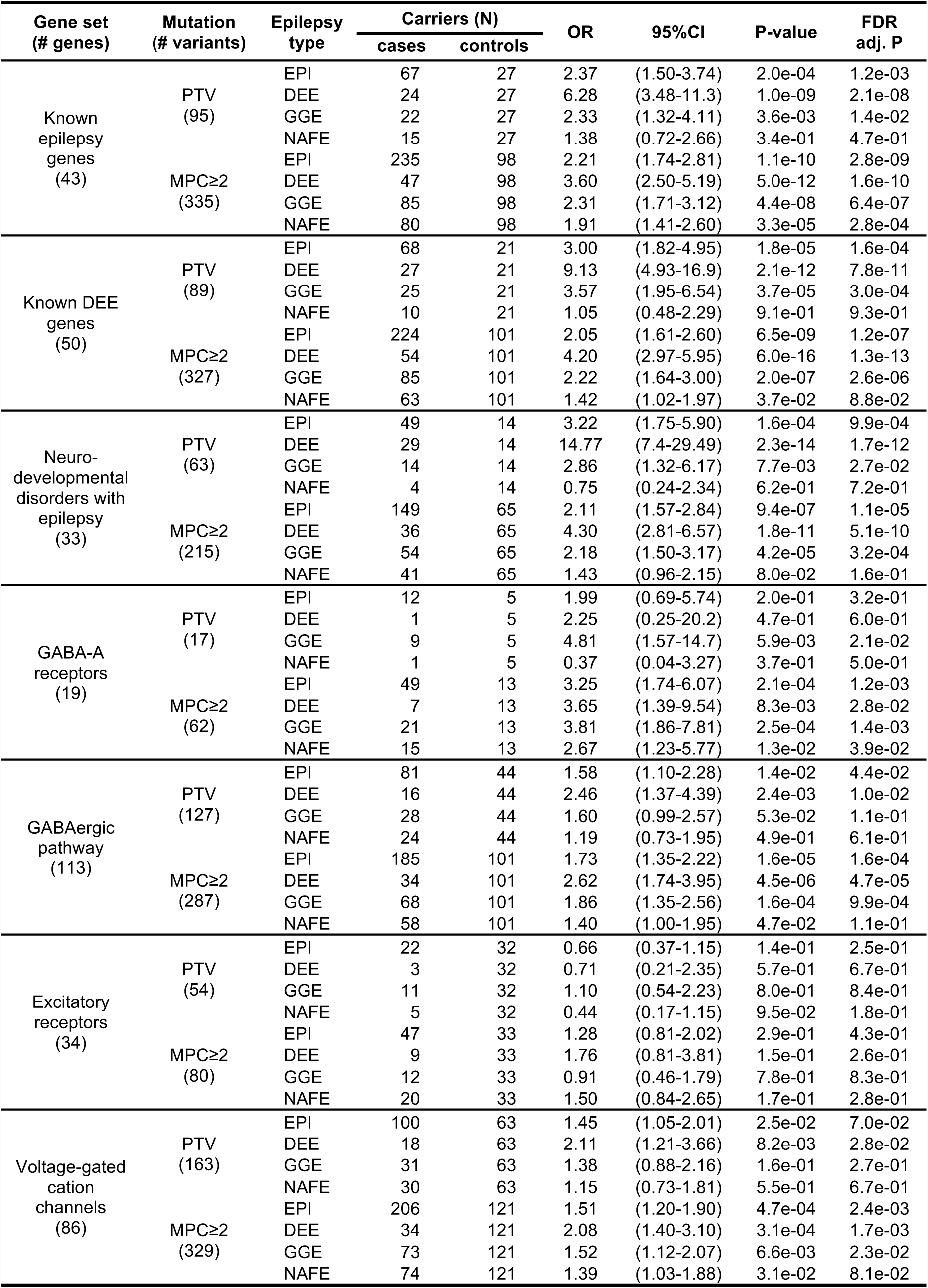
Enrichment of ultra-rare protein-truncating or damaging missense (MPC≥2) singletons in epilepsy This analysis compared the burden of deleterious pathogenic variants between cases and controls using logistic regression, adjusting for sex, the first ten principal components, and overall variant count. FDR correction was based on a full list of burden tests in **Table S8**. Tested epilepsy types included all epilepsies (**EPI**; N=9,170), developmental and epileptic encephalopathies (**DEE**; N=1,021), genetic generalized epilepsy (**GGE**; N=3,108), and non-acquired focal epilepsy (**NAFE**; N=3,597). All were compared against 8,436 control samples. **Figure 2** shows the enrichment pattern of PTVs and MPC≥2 variants across the seven gene sets listed here.

For gene sets other than the three lists of previously associated genes (**Table S6**; 74 non-overlapping genes in total), we evaluated the residual burden of URVs after correcting for events in the 74 known genes. For the gene sets of cation channel and neurotransmitter receptor genes, the adjusted burden signals of singleton deleterious URVs was largely reduced, with some weak associations remaining in GABA_A_ receptor-encoding or GABAergic genes among individuals with DEE or GGE. For the larger gene groups of constrained genes and brain-enriched genes, burden signals were attenuated but many remained significant, especially the strong enrichment of missense MPC≥2 variants in brain-enriched genes across all three classes of epilepsy (**Figure S16**). These findings suggest that although most gene burden is driven by previously identified genes, more associations could be uncovered with larger sample sizes.

### Gene-based collapsing analysis recapture known genes for DEE

For gene discovery, because both protein-truncating and damaging missense (MPC≥2) URVs showed an elevated burden in epilepsy cases, we aggregated both together as deleterious pathogenic variants along with in-frame insertions and deletions in our gene collapsing analysis. This amassed to a total of 46,917 singleton URVs and 52,416 URVs with AC≤3. Surprisingly, for individuals diagnosed with DEE, we re-identified several of the established candidate DEE genes as top associations (**Figure 3A**). Although screening was not performed systematically, many DEE patients were screened-negative for these genes using clinical gene panels prior to enrollment. Based on the results of singleton URVs, *SCN1A* was the only gene that reached exome-wide significance (*OR* = 18.4, *P* = 5.8×10^-8^); other top-ranking known genes included *NEXMIF* (previously known as *KIAA2022*; *OR* > 99, *P* = 1.6×10^-6^*), KCNB1* (*OR* = 20.8, *P* = 2.5×10^-4^), *SCN8A* (*OR* = 13.8, *P* = 6.1×10^-4^), and *SLC6A1* (*OR* = 11.1, *P* = 3.6×10^-3^) (**Table S10**). Some carriers of deleterious URVs in lead genes were affected individuals with a normal result for gene panel testing; for example, 2 out of the 3 carriers of qualified URVs for *PURA* and 2 out of 5 for *KCNB1* had undergone previous genetic screening. (**Table S7**). This could be because different sample-contributing sites adopted different gene panels and not all of them included the lead genes found to carry variants qualifying from this study, or that during screening patients were found to carry a variant of uncertain significance that did not satisfy the ACMG guidelines^39^. The gene burden results held up when considering URVs with AC≤3, often showing even stronger associations; two other well-studied genes, *STXBP1* (*OR* = 13.3, *P* = 1.4×10^-5^) and *WDR45* (*OR* > 49, *P* = 1.2 × 10^-3^), emerged on top, both of which have been implicated in DEE and developmental disorders (**Table S11**).

**Figure 3.**
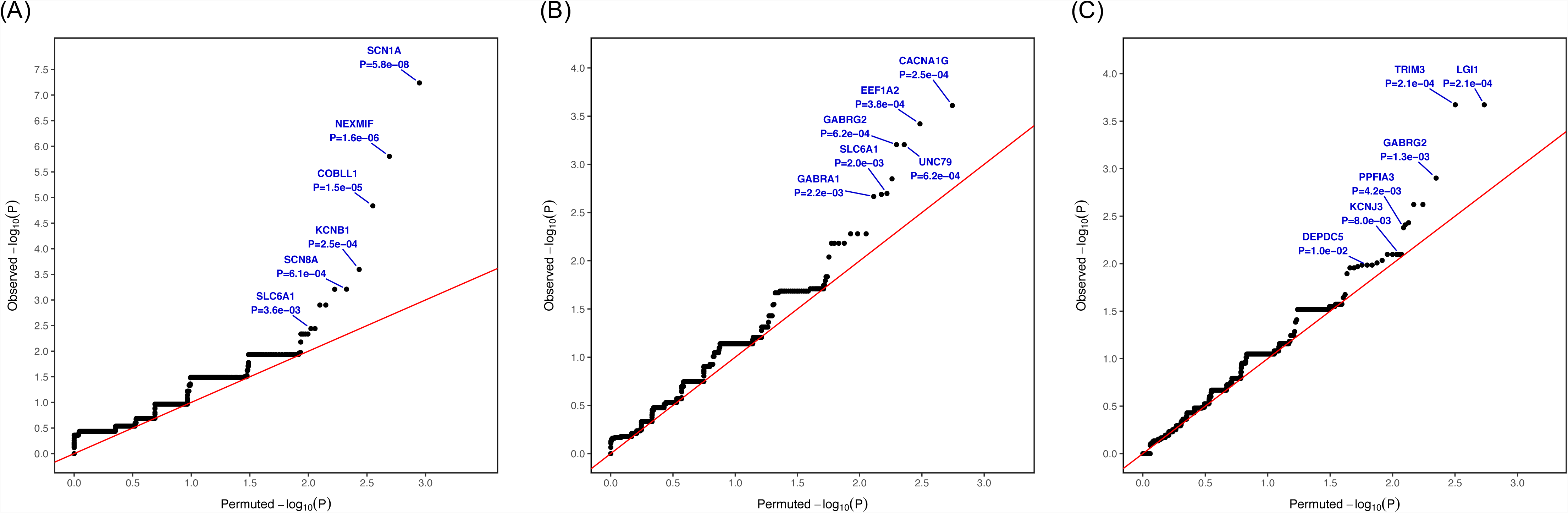
Gene burden for individuals diagnosed with (A) developmental and epileptic encephalopathies, (B) genetic generalized epilepsy, or (C) non-acquired focal epilepsy. This analysis focused on ultra-rare (non-DiscovEHR) singleton variants annotated as PTV, damaging missense (MPC≥2), or in-frame insertion/deletion and used Fisher’s exact test to identify genes with a differential carrier rate of these ultra-rare deleterious variants in individuals with epilepsy compared to controls. Exome-wide significance was defined as p-value < 6.8e-07 after Bonferroni correction (Methods). Only *SCN1A* achieved exome-wide significance for individuals with DEE.

### Channel and transporter genes implicated in common epilepsies

When evaluating gene burden in the GGE and NAFE epilepsy subgroups, we did not identify any exome-wide significant genes. However, several candidate epilepsy genes made up the lead associations, including ion channel and transporter genes known to cause rare forms of epilepsy. For the GGE case-control analysis in singleton deleterious URVs, the lead associations included four previously-associated genes (*EEF1A2, OR* = 32, *P* = 3.8×10^-4^; *GABRG2*, *OR* = 19.0, *P* = 6.2×10^-4^; *SLC6A1*, *OR* = 7.3, *P* = 2.0×10^-3^; and *GABRA1, OR* = 9.5, *P* = 2.2×10^-3^), and two genes (*CACNA1G, OR* = 9.1, *P* = 2.5×10^-4^; *UNC79, OR* = 19.0, *P* = 6.2×10^-4^) that were not previously linked to epilepsy but are both highly expressed in the brain and under evolutionary constraint (**Figures 3B**; **Table S12**). Although evidence has been mixed, *CACNA1G* was previously implicated as a potential susceptibility gene for GGE in mutational analysis^40^ and reported to modify mutated sodium channel (*SCN2A*) activity in epilepsy^41^. *UNC79* is an essential part of the UNC79-UNC80-NALCN channel complex that influences neuronal excitability by interacting with extracellular calcium ions^42^, and this channel complex has been previously associated with infantile encephalopathy^43^. Notably, all these lead genes were more enriched for damaging missense (MPC≥2) than for protein-truncating URVs despite the lower rate of MPC≥2 variants relative to PTVs (**Table S12**).

For individuals with NAFE, the analysis of singleton deleterious URVs identified *LGI1* and *TRIM3* as the top two genes carrying a disproportionate number of deleterious URVs, however neither reached exome-wide significance (*OR* > 32, *P* = 2.1×10^-4^). *GABRG2*, a lead association in individuals with GGE, was among the top ten most enriched genes, along with two brain-enriched, constrained genes (*PPFIA3, OR* = 8.2, *P* = 4.2×10^-3^; and *KCNJ3, OR* = 16.4, *P* = 1.2×10^-3^). *GABRG2* has previously been reported to show an enrichment of variants compared to controls in a cohort of individuals with Rolandic epilepsy (childhood epilepsy with centrotemporal spikes) or related phenotypes, the most common group of focal epilepsies of childhood^44^. Two other genes previously associated with epilepsy, *DEPDC5* and *SCN8A* (both *OR* = 5.5, *P* = 0.01), were among the top twenty associations (**Figures 3C**; **Table S14**). *LGI1* and *DEPDC5* are established genes for focal epilepsy, and *DEPDC5* was the only exome-wide significant hit in the Epi4K WES study for familial NAFE cases^17^. *TRIM3* has not been previously implicated in epilepsy, but evidence from a mouse model study implicates it in regulation of GABA_A_ receptor signaling and thus modulation of seizure susceptibility^45^. Single gene burden for both GGE and NAFE remained similar when considering URVs with an allele count up to AC≤3 (**Tables S13 & S15**). Gene burden tests collapsing all epilepsy phenotypes recapitulated the lead genes in each of the subgroup-specific analyses, but none of the genes achieved exome-wide significance (**Tables S16 & S17**). It is worth noting that some of the genes were enriched for deleterious URVs among the “controls”, which is clearly driven by non-neuropsychiatric disease ascertainment for many of the available controls (e.g., *LDLR* in **Table S16**; most control carriers were individuals with cardiovascular diseases from the MIGen cohorts in **Table S2**). Thus, these should not be interpreted as potential protective signals for epilepsy.

### Recessive model, SKAT gene test, and single variant association

The secondary gene-based test of a recessive model did not identify genes that differed significantly in the carrier rate of homozygous deleterious variants between epilepsy-affected individuals and controls (**Table S18**). Even if we considered variants up to MAF < 0.01, for most of the lead genes, only one case carrier was identified. For the DEE cohort, these genes included recessive genes previously implicated, such as *ARV1, BRAT1, CHRDL*^46^ with a homozygous PTV and *OPHN1*^46^ with a recessive missense (MPC≥2) variant (**Table S18A**). For the two common forms of epilepsy, a few studied recessive epilepsy genes were also observed in the lead gene associations, such as *SLC6A8*^46^ (a homozygous PTV) for GGE (**Table S18B**), and *SLC6A8* (a homozygous missense-MPC) and *SYN1*^46^ (a homozygous PTV) for NAFE (**Table S18C**). One GGE-affected individual was found homozygous for an in-frame deletion on *CHD2*, a dominant DEE gene^46^ (**Table S18B**). These findings suggest an even larger cohort will be needed to identify with clarity risk genes that act in a recessive manner for different groups of epilepsy.

Beyond URVs, we studied the contribution of low frequency deleterious variants to epilepsy risk using SKAT (MAF < 0.01). Top associations for individuals with DEE included known genes, such as missense-enriched *STXBP1* (*P* = 9.3×10^-9^), *KCNA2* (*P* = 1.0×10^-5^; **Figure S17**), and PTV-enriched *NEXMIF* (*P* = 7.1×10^-8^), and *SCN1A* (*P* = 3.9×10^-4^; **Figure S18**). However, no significant gene enrichment was observed for the two common types of epilepsy or when combining all epilepsy cases. The tests for PTVs and missense variants with MPC≥2 were mostly underpowered due to sparse observations (**Figure S17 & S18**). No individual low-frequency variant (MAF > 0.001) was significantly associated with overall epilepsy or with any of the studied epilepsy phenotypes (**Figure S19**). The primary gene-based test results and single variant associations are available on our Epi25 WES browser (**Web Resources**).

## Discussion

In the largest exome study of epilepsies to date, we show that ultra-rare deleterious coding variation—variation absent in a large population-based exome database—is enriched in both rare and common epilepsy syndromes when compared to ancestrally matched controls. When all genes were considered in the tested gene sets, PTVs showed a more significant signal than missense variants with an MPC≥2, and enrichment in deleterious URVs was more pronounced in individuals diagnosed with DEE and GGE relative to NAFE. While no single gene surpassed exome-wide significance in the non-hypothesis-driven analysis for GGE or NAFE, specific gene groups previously associated with epilepsy or encoding biologically interesting entities showed a clear enrichment of deleterious URVs. Specifically, we observed a significant excess of deleterious URVs in constrained genes, established epilepsy genes, GABA_A_ receptor subunit genes, a larger group of genes delineating the GABAergic pathway, and all cation channel-encoding genes. Our results thus support the concept that defects in GABAergic inhibition underlie various forms of epilepsy. These findings, based on a more than 5-fold increase in sample size over previous exome-sequencing studies^17-19;47^, clearly reveal observations that have been hypothesized for common epilepsies from studies of rare, large monogenic families, and confirm that the same genes are relevant in both settings. Thus, a further increase in sample size will continue to unravel the complex genetic architecture of common epilepsies. Interestingly, no enrichment was seen in genes encoding the excitatory glutamate and acetylcholine receptors. For GGE, this difference between variants in inhibitory versus excitatory receptor genes may be real, as excitatory receptor variants have not been shown so far in single subjects or families. In NAFE, however, we suspect that it is probably due to a lack of power and/or genetic heterogeneity, since genetic variants in specific subunits of nicotinic acetylcholine and NMDA receptors have been described extensively in different types of non-acquired familial focal epilepsies^48^.

Notably, our overall finding of a mild to moderate burden of deleterious coding URVs in NAFE (**Figure 1** & **2**) contrasts with results reported in the Epi4K WES study, where the familial NAFE cohort showed a strong enrichment signal of ultra-rare functional variation in known epilepsy genes and ion channel genes^17^. In addition, our findings for GGE showed a genetic risk comparable or even stronger than the Epi4K familial GGE cohort. The strong signal in our GGE cohort likely reflects the larger sample size, whereas the weaker signal in our NAFE cohort is most likely due to differences in patient ascertainment. In Epi4K the cohort was deliberately enriched with familial cases, most of whom had an affected first-degree relative and were ascertained in sibling or parent-child pairs or multiplex families, and familial NAFE is relatively uncommon. In the Epi25 collaboration, a positive family history of epilepsy was not a requirement and only 9% of DEE, 12% of GGE, and 5% of NAFE patients had a known affected first-degree relative. Indeed, our results were consistent with the Epi4K sporadic NAFE cohort, where no signals of enrichment were observed^17; 49^. This difference may reflect the substantial etiological and genetic heterogeneity of epilepsy even within subgroups especially in NAFE. In particular, the dramatically weaker genetic signals, per sample, observed in individuals with NAFE studied here compared with those in the previous Epi4K study illustrate a pronounced difference in the genetic signals associated with familial and non-familial NAFE. The reasons for this striking difference remain to be elucidated. Comparing the two common classes of epilepsy, our findings showed a larger genetic burden from URVs for GGE relative to NAFE, which could be due to heterogeneity in electroclinical syndromes within each class and should not be viewed as conclusive. On the other hand, in the latest GWAS of common epilepsies of 15,212 cases and 29,677 controls from the ILAE Consortium^15^, fewer GWAS hits were discovered and less heritability was explained by common genetic variation for the focal epilepsy cohort (9.2%) compared to the GGE cohort (32.1%), suggesting that current evidence from both common and rare variant studies are converging on a larger genetic component underlying the etiology of non-familial cases of GGE relative to NAFE, as originally postulated.

We found that ultra-rare missense variants with an MPC score^28^ ≥ 2 (2.0% of missense variants) were enriched in individuals with epilepsy at an effect size approaching PTVs in the investigated gene groups. For common epilepsy types, the burden of these missense variants (MPC≥2) was even more prominent than PTVs in known epilepsy genes and GABAergic genes (**Figure 2**). At the gene level, some of the top channel genes (e.g. *GABRG2, CACNA1G*) carried a higher number of missense variants (MPC≥2) than PTVs in people with epilepsy. For instance, in the gene-based collapsing analysis considering all epilepsies, 15 *GABRG2* pathogenic variants were found in epilepsy-affected individuals (including 7 GGE and 7 NAFE; **Tables S12, 14 & 16**) versus only 1 pathogenic variant in controls; among the case-specific pathogenic variants, one was a splice site mutation, while the other 14 were all missense variants (MPC≥2) (**Figure S20**), linking to an impaired channel function. This is in line with findings from a recent exome-wide study of 6,753 individuals with neurodevelopmental disorder with and without epilepsy^9^ that detected an association of missense *de novo* variants with the presence of epilepsy, particularly when considering only ion channel genes. A disease-association of missense variants rather than PTVs points to a pathophysiological mechanism of protein-alteration (e.g., gain-of-function or dominant-negative effects) rather than haploinsufficiency, but ultimately only functional tests can elucidate these mechanisms. A recent study on the molecular basis of 6 *de novo* missense variants in *GABRG2* identified in DEE reported an overall reduced inhibitory function of *GABRG2* due to decreased cell surface expression or GABA-evoked current amplitudes, suggesting GABAergic disinhibition as the underlying mechanism^50^. Surprisingly, 2 of those recurrent *de novo* missense variants were seen in two GGE-affected individuals in our study (A106T and R323Q), and another recently reported variant in *GABRB2* (V316I) also occurred both *de novo* in DEE^51^ and as an inherited variant in a GGE family showing a loss of receptor function^18^. This suggests that changes in protein function from the same missense pathogenic variant may cause not only severe epilepsy syndromes, but also contribute to common epilepsies with milder presentations, similar to what is known about variable expressivity in large families carrying *GABRG2* variants^48;^ 52-54. Reduced receptor function due to *GABRG2* variants has been also shown for childhood epilepsy with centrotemporal spikes previously^44; 54^, which belong to the NAFE group in this study. Moving forward, discovering how variant-specific perturbations of the neurotransmission and signaling system in a gene can link to a spectrum of epilepsy syndromes will require in-depth functional investigation.

Although we have increased the sample size from the Epi4K and EuroEPINOMICS WES studies for both GGE and NAFE subgroups by more than 5-fold, the phenotypic and genetic heterogeneity of common epilepsies—on par with other complex neurological and neuropsychiatric conditions—will require many more samples to achieve statistical power for identifying exome-wide significant genes. Furthermore, while we implemented stringent QC to effectively control for the exome capture differences between cases and controls, this concomitantly resulted in a loss of a substantial amount of the called sites and reduced our detection power to identify associated variants. As sample sizes grow, the technical variation across projects and sample collections will remain a challenge in large-scale sequencing studies relying on a global collaborative effort.

With this largest epilepsy WES study to date, we demonstrated a strong replicability of existing gene findings in an independent cohort. GABA_A_ receptor genes affected by predicted-pathogenic missense variants were enriched across the three subgroups of epilepsy. An ongoing debate in epilepsy genetics is the degree to which generalized and focal epilepsies segregate separately, and whether their genetic determinants are largely distinct or sometimes shared^4;55^. Whilst clinical evidence for general separation of pathophysiological mechanisms in these two forms is strong, and most monogenic epilepsy families segregate either generalized or focal syndromes, the distinction is not absolute. Here, the finding of rare variants in GABA_A_ receptor genes in both forms adds weight to the case for shared genetic determinants.

Our results suggest that clinical presentations of common epilepsy types with complex inheritance patterns have a combination of both common and rare genetic risk variants. The latest ILAE epilepsy GWAS of over 15,000 patients and 25,000 controls identified 16 genome-wide significant loci for common epilepsies^15^, mapped these loci to ion channel genes, transcriptional factors, and pyridoxine metabolism, and implicated a role in epigenetic regulation of gene expression in the brain. A combination of rare and common genetic association studies with large sample sizes, along with the growing evidence from studies of copy number variation and tandem repeat expansions in epilepsy^12; 56; 57^, will further decipher the genetic landscape of common epilepsy subgroups. The ongoing effort of the Epi25 collaborative is expected to double the patient cohorts in upcoming years with the goal of elucidating shared and distinct gene discoveries for common and rare forms of epilepsy, ultimately facilitating precision medicine strategies in the treatment of epilepsy.

## Supplemental Data

Supplemental data includes affiliations of the contributing authors, descriptions of patient recruitment and phenotyping from individual participating cohorts, supplemental acknowledgment, 20 figures and 18 tables.

## Consortia

### Author contributions

S.F.B., H.L., D.B.G., and D.H.L conceived the consortium and contributed to the study design. S.F.B., C.D., D.J.D., D.H.L., A.G.M., I.E.S., and P.S. designed patient recruitment and phenotyping criteria. D.H.L., D.B.G., H.L., S.F.B, C.D., D.J.D., A.G.M., I.E.S., P.S, C.F., R.K., K.M., B.M.R., S.T.B., G.L.C., P.C., C.C., P.D.G., T.D-S., R.G., H.H., E.L.H., I.H., P.K., S.P., S.M.S., R.S., and S.W. advised and approved the consortium activities. Authors from individual patient and control cohorts (Supplemental Data) contributed to site-specific recruitment and phenotyping. B.M.N., M.J.D., D.P.H., and F.C. directed and managed sample aggregation and sequencing effort at the Broad Institute. Sequence data generation was directed by S.B.G and managed by F.C. and N.G., with E.S.L, M.J.D., and B.M.N. overseeing the work. B.M.N, D.B.G., C.C., D.L, L.E.A., M.J.D., E.L.H., G.L.C., H.H., H.L., I.H., R.K., S.W., S.K., and S.P. supervised and reviewed genetic data analyses. Lead analyst Y.A.F. conducted the analyses with D.P.H, L.E.A., and K.T. Y.A.F. wrote the manuscript and interpreted the results with B.M.N., D.P.H., E.S.L., S.F.B., H.L., D.B.G., E.L.H., and D.H.L.

## Acknowledgments

We gratefully thank the Epi25 principal investigators, local staff from individual cohorts, and all of the patients with epilepsy who participated in the study for making possible this global collaboration and resource to advance epilepsy genetics research. This work is part of the Centers for Common Disease Genomics (CCDG) program, funded by the National Human Genome Research Institute (NHGRI) and the National Heart, Lung, and Blood Institute (NHLBI). CCDG research activities at the Broad Institute was supported by NHGRI grant UM1 HG008895. The Genome Sequencing Program efforts were also supported by NHGRI grant 5U01HG009088-02.

The content is solely the responsibility of the authors and does not necessarily represent the official views of the National Institutes of Health. Supplemental grant for Epi25 phenotyping was supported by “Epi25 Clinical Phenotyping R03”, National Institutes of Health (1R03NS108145-01), with D.H.L and S.F.B as the principal investigators. Additional funding sources and acknowledgment of individual patient and control cohorts were listed in **Supplemental Data**. We thank the Stanley Center for Psychiatric Research at the Broad Institute for supporting sequencing effort and control sample aggregation. The authors would like to thank the Discov-EHR collaboration of Geisinger Health System and Regeneron for providing exome variant data for comparison. We also thank Sali Farhan, Kyle Satterstrom, and Chai-Yen Chen for helpful discussions, Nick Watts and Matthew Solomonson for browser development, and the Hail team for analysis support.

## Web Resources

The URLs for the consortium, data, and results presented herein are as follows: Epi25 Collaborative, http://epi-25.org/

Exome Aggregation Consortium (ExAC), http://exac.broadinstitute.org

The DiscovEHR cohort, http://www.discovehrshare.com

Epi25 Year1 whole-exome sequence data on dbGaP, http://www.ncbi.nlm.nih.gov/gap through accession number phs001489 (the current study includes Year1-2 samples, and the Year2 data will later be made available)

Epi25 WES results browser, http://epi25.broadinstitute.org/

