## Supplemental Data 1 for "Ultra-rare genetic variation in the epilepsies: a whole-exome sequencing study of 17,606 individuals"

**Table of contents**

**Supplemental Subjects and Methods**
A full list of authors and affiliations
Details of individual participating Epi25 cohorts
Details of individual control cohorts (not including dbGaP samples)
Supplemental acknowledgments

**Supplemental Figures**
**Figure S1.** Variant pre-filtering
**Figure S2.** Average coverage depth
**Figure S3.** PCA: all samples
**Figure S4.** PCA: European samples
**Figure S5.** Sex check
**Figure S6.** IBD filtering
**Figure S7.** Ti/Tv, Het/Hom, and Indel ratios
**Figure S8.** Post-QC allele count distribution, by functional class
**Figure S9.** Exome-wide burden of URVs in all-epilepsy vs. controls
**Figure S10.** Exome-wide burden of rare and low frequency variants in all-epilepsy vs. controls
**Figure S11.** Correcting background variation in gene-set burden test
**Figure S12.** Burden of URVs in LoF-intolerant genes using non-ExAC controls: 0.9 < pLI < 0.995
**Figure S13.** Burden of URVs in LoF-intolerant genes using non-ExAC controls: pLI >0.9 vs. pLI >0.995
**Figure S14.** Burden of URVs in missense constrained genes
**Figure S15.** Burden of URVs in brain-specific genes
**Figure S16.** Burden of ultra-rare singleton PTVs and missese-MPC≥2 variants before and after correcting for events in genes previously associated with epilepsy
**Figure S17.** Burden of deleterious missense variants (MAF < 0.01) assessed using SKAT
**Figure S18.** Burden of protein-truncating variants (MAF < 0.01) assessed using SKAT
**Figure S19.** Association with common and low frequency variants
**Figure S20.** Mutational diagram of selected top genes in gene burden analysis

**Supplemental Tables**
**Table S1.** Patient cohort collection
**Table S2.** Control sample collection
**Table S3.** Sample QC and number of sample dropout at each QC step
**Table S4.** Variant QC and number of variant dropout at each QC step
**Table S5.** Grouping of functional consequences of the called sites
**Table S6.** Gene sets collected for burden analysis
**Table S7.** Prior gene panel screening of the 1,021 DEE-affected individuals
*In a separate Excel file (large tables):*
**Table S8.** Gene set burden of URVs for each epilepsy type versus controls (AC=1)
**Table S9.** Gene set burden of URVs for each epilepsy type versus controls (AC≤3)
**Table S10.** Top 200 genes with burden of deleterious URVs in developmental and epileptic encephalopathy (AC=1)
**Table S11.** Top 200 genes with burden of deleterious URVs in developmental and epileptic encephalopathy (AC≤3)
**Table S12.** Top 200 genes with burden of deleterious URVs in genetic generalized epilepsy (AC=1)
**Table S13.** Top 200 genes with burden of deleterious URVs in genetic generalized epilepsy (AC≤3)
**Table S14.** Top 200 genes with burden of deleterious URVs in non-acquired focal epilepsy (AC=1)
**Table S15.** Top 200 genes with burden of deleterious URVs in non-acquired focal epilepsy (AC≤3)
**Table S16.** Top 200 genes with burden of deleterious URVs in all epilepsy cases (AC=1)
**Table S17.** Top 200 genes with burden of deleterious URVs in all epilepsy cases (AC≤3)
**Table S18.** Top genes with burden of deleterious URVs for the recessive model by epilepsy subgroup

**Supplemental References**

**Supplemental Subjects and Methods**

***A full list of authors and affiliations***

**Epi25 sequencing, analysis, and project management at the Broad Institute**
Yen-Chen Anne Feng[1-4], Daniel P. Howrigan[1,3,4], Liam E. Abbott[1,3,4], Katherine Tashman[1,3,4], Felecia Cerrato[3], Tarjinder Singh[1,3,4], Henrike Heyne[1,3,4], Andrea E. Byrnes[1,3,4], Dennis Lal[4,5], Claire Churchhouse[1,3,4], Namrata Gupta[3], Stacey B. Gabriel[146], Mark J. Daly[1,3,4], Eric S. Lander[146,147,148], Benjamin M. Neale[1,3,4]

**Epi25 executive committee**
Samuel F. Berkovic [9], Holger Lerche[8], David B. Goldstein [6], Daniel H. Lowenstein[7]

**Epi25 strategy, phenotyping, analysis, informatics, and project management committees**
Samuel F. Berkovic[9], Holger Lerche[8], David B. Goldstein[6], Daniel H. Lowenstein[7], Gianpiero L. Cavalleri[67,70], Patrick Cossette[106], Chris Cotsapas[111], Peter De Jonghe[12-14], Tracy Dixon-Salazar[112], Renzo Guerrini[82], Hakon Hakonarson[101], Erin L. Heinzen[6], Ingo Helbig[28,29,102], Patrick Kwan[10,11], Anthony G. Marson[58], Slavé Petrovski[11,116], Sitharthan Kamalakaran[6], Sanjay M. Sisodiya[57], Randy Stewart[113], Sarah Weckhuysen[12-14], Chantal Depondt[15], Dennis J. Dlugos[101], Ingrid E. Scheffer[9], Pasquale Striano[74], Catharine Freyer[7], Roland Krause[114], Patrick May[114], Kevin McKenna[7], Brigid M. Regan[9], Susannah T. Bellows[9], Costin Leu[4,5,57]

**Authors from individual Epi25 cohorts:**

**Australia: Melbourne (AUSAUS)**
Samuel F. Berkovic[9], Ingrid E. Scheffer[9], Brigid M. Regan[9], Caitlin A. Bennett[9], Susannah T. Bellows[9], Esther M.C. Johns[9], Alexandra Macdonald[9], Hannah Shilling[9], Rosemary Burgess[9], Dorien Weckhuysen[9], Melanie Bahlo[119,120]

**Australia: Royal Melbourne (AUSRMB)**
Terence J. O'Brien[10,11], Patrick Kwan[10,11], Slavé Petrovski[11,116], Marian Todaro[10,11]

**Belgium: Antwerp (BELATW)**
Sarah Weckhuysen[12-14], Hannah Stamberger[12-14], Peter De Jonghe[12-14]

**Belgium: Brussels (BELULB)**
Chantal Depondt[15]

**Canada: Andrade (CANUTN)**
Danielle M. Andrade[16,17], Tara R. Sadoway[17], Kelly Mo[17]

**Switzerland: Bern (CHEUBB)**
Heinz Krestel[18], Sabina Gallati[19]

**Cyprus (CYPCYP)**
Savvas S. Papacostas[20], Ioanna Kousiappa[20], George A. Tanteles[21]

**Czech Republic: Prague (CZEMTH)**
Katalin Štěrbová[22], Markéta Vlčková[23], Lucie Sedláčková[22], Petra Laššuthová[22]

**Germany: Frankfurt/Marburg (DEUPUM)**
Karl Martin Klein[24,25], Felix Rosenow[24,25], Philipp S. Reif[24,25], Susanne Knake[25]

**Germany: Bonn (DEUUKB)**
Wolfram S. Kunz[26,27], Gábor Zsurka[26,27], Christian E. Elger[27], Jürgen Bauer[27], Michael Rademacher[27]

**Germany: Kiel (DEUUKL)**
Ingo Helbig[28,29,102], Karl Martin Klein[24,25], Manuela Pendziwiat[29], Hiltrud Muhle[29], Annika Rademacher[29], Andreas van Baalen[29], Sarah von Spiczak[29], Ulrich Stephani[29], Zaid Afawi[30], Amos D. Korczyn[31], Moien Kanaan[32], Christina Canavati[32], Gerhard Kurlemann[33], Karen Müller-Schlüter[34], Gerhard Kluger[35,36], Martin Häusler[37], Ilan Blatt[31,115]

**Germany: Leipzig (DEUULG)**
Johannes R. Lemke[38], Ilona Krey[38]

**Germany: Tuebingen (DEUUTB)**
Holger Lerche[8], Yvonne G. Weber[8,151], Stefan Wolking[8], Felicitas Becker[8,39], Christian Hengsbach[8], Sarah Rau[8], Ana F. Maisch[8], Bernhard J. Steinhoff[40], Andreas Schulze-Bonhage[41], Susanne Schubert-Bast[42], Herbert Schreiber[43], Ingo Borggräfe[44], Christoph J. Schankin[45], Thomas Mayer[46], Rudolf Korinthenberg[47], Knut Brockmann[48], Gerhard Kurlemann[33], Dieter Dennig[49], Rene Madeleyn[50]

**Finland: Kuopio (FINKPH)**
Reetta Kälviäinen[51], Pia Auvinen[51], Anni Saarela[51]

**Finland: Helsinki (FINUVH)**
Tarja Linnankivi[52], Anna-Elina Lehesjoki[53]

**Wales: Swansea (GBRSWU)**
Mark I. Rees[54,55], Seo-Kyung Chung[54,55], William O. Pickrell[54], Robert Powell[54,56]

**UK: UCL (GBRUCL)**
Sanjay M. Sisodiya[57], Natascha Schneider[57], Simona Balestrini[57], Sara Zagaglia[57], Vera Braatz[57]

**UK: Imperial/Liverpool (GBRUNL)**
Anthony G. Marson[58], Michael R. Johnson[59], Pauls Auce[60], Graeme J. Sills[61]

**Hong Kong (HKGHKK)**
Patrick Kwan[10,11,62], Larry W. Baum[117,118,63], Pak C. Sham[117,118,63], Stacey S. Cherny[64], Colin H.T. Lui[65]

**Croatia (HRVUZG)**
Nina Barišić[66]

**Ireland: Dublin (IRLRCI)**
Gianpiero L. Cavalleri[67,70], Norman Delanty[67,70], Colin P. Doherty[68,70], Arif Shukralla[69], Mark McCormack[67], Hany El-Naggar[69,70]

**Italy: Milan (ITAICB)**
Laura Canafoglia[71], Silvana Franceschetti[71], Barbara Castellotti[72], Tiziana Granata[73]

**Italy: Genova (ITAIGI)**
Pasquale Striano[74], Federico Zara[75], Michele Iacomino[75], Francesca Madia[75], Maria Stella Vari[74], Maria Margherita Mancardi[75], Vincenzo Salpietro[74]

**Italy: Bologna (ITAUBG)**
Francesca Bisulli[76,77], Paolo Tinuper[76,77], Laura Licchetta[76,77], Tommaso Pippucci[78], Carlotta Stipa[79], Raffaella Minardi[76]

**Italy: Catanzaro (ITAUMC)**
Antonio Gambardella[80], Angelo Labate[80], Grazia Annesi[81], Lorella Manna[81], Monica Gagliardi[81]

**Italy: Florence (ITAUMR)**
Renzo Guerrini[82], Elena Parrini[82], Davide Mei[82], Annalisa Vetro[82], Claudia Bianchini[82], Martino Montomoli[82], Viola Doccini[82], Carla Marini[82]

**Japan: RIKEN Institute (JPNRKI)**
Toshimitsu Suzuki[83], Yushi Inoue[84], Kazuhiro Yamakawa[83]

**Lithuania (LTUUHK)**
Birute Tumiene[85,86]

**New Zealand: Otago (NZLUTO)**
Lynette G. Sadleir[87], Chontelle King[87], Emily Mountier[87]

**Turkey: Bogazici (TURBZU)**
S. Hande Caglayan[88], Mutluay Arslan[89], Zuhal Yapıcı[90], Uluc Yis[91], Pınar Topaloglu[90], Bulent Kara[92], Dilsad Turkdogan[93], Aslı Gundogdu-Eken[88]

**Turkey: Istanbul (TURIBU)**
Nerses Bebek[94,95], Sibel Uğur-İşeri[95], Betül Baykan[94], Barış Salman[95], Garen Haryanyan[94], Emrah Yücesan[149], Yeşim Kesim[94], Çiğdem Özkara[96]

**USA: BCH (USABCH)**
Annapurna Poduri[97,98]

**USA: Philadelphia/CHOP (USACHP) and Philadelphia/Rowan (USACRW)**
Russell J. Buono[99,100,101], Thomas N. Ferraro[99,102], Michael R. Sperling[100], Dennis J. Dlugos[101,102], Warren Lo[103], Michael Privitera[104], Jacqueline A. French[105], Patrick Cossette[106], Steven Schachter[107], Hakon Hakonarson[101]

**USA: EPGP (USAEGP)**
Daniel H. Lowenstein[7], Ruben I. Kuzniecky[108], Dennis J. Dlugos[101,102], Orrin Devinsky[105]

**USA: NYU HEP (USAHEP)**
Daniel H. Lowenstein[7], Ruben I. Kuzniecky[108], Jacqueline A. French[105], Manu Hegde[7]

**USA: Penn/CHOP (USAUPN)**
Ingo Helbig[28,102], Pouya Khankhanian[109,110], Katherine L. Helbig[28], Colin A. Ellis[110]

**Authors from individual control cohorts (not including samples obtained from dbGaP):**

**Italian controls**
Gianfranco Spalletta[121,122], Fabrizio Piras[121], Federica Piras[121], Tommaso Gili[123,121], Valentina Ciullo[121,124]

**German controls**
Andreas Reif[125,126]

**UK/IRL controls 1**
Andrew McQuillin[127], Nick Bass[127]

**UK/IRL controls 2**
Andrew McIntosh[128], Douglas Blackwood[128], Mandy Johnstone[128]

**FINRISK controls**
Aarno Palotie[1,2,4,129,130]

**Genomic Psychiatry Cohort (GPC) controls**
Michele T. Pato[131], Carlos N. Pato[131], Evelyn J. Bromet[132], Celia Barreto Carvalho[133], Eric D. Achtyes[134], Maria Helena Azevedo[135], Roman Kotov[132], Douglas S. Lehrer[136], Dolores Malaspina[137], Stephen R. Marder[138], Helena Medeiros[131], Christopher P. Morley[139], Diana O. Perkins[140], Janet L Sobell[141], Peter F. Buckley[142], Fabio Macciardi[143], Mark H. Rapaport[144], James A. Knowles[131], Genomic Psychiatry Cohort (GPC) Consortium, Ayman H. Fanous[131,145], Steven A. McCarroll[3,4,150]

Affiliations:

1 Analytic and Translational Genetics Unit, Department of Medicine, Massachusetts General Hospital and Harvard Medical School, Boston, MA 02114, USA
2 Psychiatric & Neurodevelopmental Genetics Unit, Department of Psychiatry, Massachusetts General Hospital and Harvard Medical School, Boston, MA 02114, USA
3 Program in Medical and Population Genetics, Broad Institute of Harvard and MIT, 7 Cambridge Center, Cambridge, MA 02142, USA
4 Stanley Center for Psychiatric Research, Broad Institute of Harvard and MIT, Cambridge, MA 02142, USA
5 Genomic Medicine Institute, Cleveland Clinic, Cleveland, OH 44195, USA
6 Institute for Genomic Medicine, Columbia University, New York, NY 10032, USA
7 Department of Neurology, University of California, San Francisco, CA 94110, USA
8 Department of Neurology and Epileptology, Hertie Institute for Clinical Brain Research, University of Tübingen, 72076 Tübingen, Germany
9 Epilepsy Research Centre, Department of Medicine, University of Melbourne, Victoria, Australia
10 Department of Neuroscience, Central Clinical School, Monash University, Alfred Hospital, Melbourne, Australia
11 Departments of Medicine and Neurology, University of Melbourne, Royal Melbourne Hospital, Parkville, Australia
12 Neurogenetics Group, Center for Molecular Neurology, VIB, Antwerp, Belgium
13 Laboratory of Neurogenetics, Institute Born-Bunge, University of Antwerp, Belgium
14 Division of Neurology, Antwerp University Hospital, Antwerp, Belgium
15 Department of Neurology, Université Libre de Bruxelles, Brussels, Belgium
16 Department of Neurology, Toronto Western Hospital, Toronto, ON M5T 2S8, Canada
17 University Health Network, University of Toronto, Toronto, ON, Canada

[18] Departments of Neurology and BioMedical Research, Bern University Hospital and University of Bern, Bern, Switzerland
[19] Institute of Human Genetics, Bern University Hospital, Bern, Switzerland
[20] Neurology Clinic B, The Cyprus Institute of Neurology and Genetics, 2370 Nicosia, Cyprus
[21] Department of Clinical Genetics, The Cyprus Institute of Neurology and Genetics, 2370 Nicosia, Cyprus
[22] Department of Paediatric Neurology, 2nd Faculty of Medicine, Charles University and Motol Hospital, Prague, Czech Republic
[23] Department of Biology and Medical Genetics, 2nd Faculty of Medicine, Charles University and Motol Hospital, Prague, Czech Republic
[24] Epilepsy Center Frankfurt Rhine-Main, Center of Neurology and Neurosurgery, Goethe University Frankfurt, Frankfurt, Germany
[25] Epilepsy Center Hessen-Marburg, Department of Neurology, Philipps University Marburg, Marburg, Germany
[26] Institute of Experimental Epileptology and Cognition Research, University Bonn, 53127 Bonn, Germany
[27] Department of Epileptology, University Bonn, 53127 Bonn, Germany
[28] Division of Neurology, Children's Hospital of Philadelphia, Philadelphia, PA 19104, USA
[29] Department of Neuropediatrics, Christian-Albrechts-University of Kiel, 24105 Kiel, Germany
[30] Sackler School of Medicine, Tel-Aviv University, Ramat Aviv, Israel
[31] Tel-Aviv University Sackler Faculty of Medicine, Ramat Aviv 69978, Israel
[32] Hereditary Research Lab, Bethlehem University, Bethlehem, Palestine
[33] Department of Neuropediatrics, Westfälische Wilhelms-University, Münster, Germany
[34] Epilepsy Center for Children, University Hospital Neuruppin, Brandenburg Medical School, Neuruppin, Germany
[35] Neuropediatric Clinic and Clinic for Neurorehabilitation, Epilepsy Center for Children and Adolescents, Vogtareuth, Germany
[36] Research Institute Rehabilitation / Transition / Palliation, PMU Salzburg, Austria
[37] Division of Neuropediatrics and Social Pediatrics, Department of Pediatrics, University Hospital, RWTH Aachen, Aachen, Germany
[38] Institute of Human Genetics, Leipzig, Germany
[39] RKU-University Neurology Clinic of Ulm, Ulm, Germany
[40] Kork Epilepsy Center, Kehl-Kork, Germany
[41] Epilepsy Center, University of Freiburg, Freiburg im Breisgau, Germany
[42] Section Neuropediatrics and Inborn Errors of Metabolism, University Children's Hospital, Heidelberg, Germany
[43] Neurological Practice Center & Neuropoint Patient Academy, Ulm, Germany.
[44] Department of Pediatric Neurology and Developmental Medicine, LMU Munich, Munich, Germany
[45] Department of Neurology, University of Munich Hospital-Großhadern, Munich, Germany
[46] Saxonian Epilepsy Center Radeberg, Radeberg, Germany
[47] Division of Neuropediatrics and Muscular Disorders, University Hospital Freiburg, Freiburg, Germany
[48] University Children's Hospital, Göttingen, Germany
[49] Private Neurological Practice, Stuttgart, Germany
[50] Department of Pediatrics, Filderklinik, Filderstadt, Germany
[51] Neurocenter, Kuopio University Hospital, Kuopio Finland and Institute of Clinical Medicine, University of Eastern Finland, Finland
[52] Child Neurology, University of Helsinki and Helsinki University Hospital, Helsinki, Finland
[53] Medicum, University of Helsinki, Helsinki, Finland and Folkhälsan Research Center, Helsinki, Finland
[54] Neurology Research Group, Swansea University Medical School, Swansea University SA2 8PP, UK
[55] Faculty of Medicine and Health, University of Sydney, Sydney, Australia
[56] Department of Neurology, Morriston Hospital, Abertawe BroMorgannwg HealthBoard, Swansea, UK
[57] Department of Clinical and Experimental Epilepsy, UCL Queen Square Institute of Neurology, London, UK and Chalfont Centre for Epilepsy, Chalfont St Peter, UK
[58] Department of Molecular and Clinical Pharmacology, University of Liverpool, Liverpool, UK
[59] Division of Brain Sciences, Imperial College London, London, UK
[60] Department of Neurology, Walton Centre NHS Foundation Trust, Liverpool, UK
[61] School of Life Sciences, University of Glasgow, Glasgow, UK

[62] Department of Medicine and Therapeutics, Chinese University of Hong Kong, Hong Kong, China
[63] Department of Psychiatry, University of Hong Kong, Hong Kong, China
[64] Department of Epidemiology and Preventive Medicine and Department of Anatomy and Anthropology, Sackler Faculty of Medicine, Tel Aviv University, Israel
[65] Department of Medicine, Tseung Kwan O Hospital, Hong Kong, China
[66] Department of Pediatric University Hospital centre Zagreb, Croatia
[67] The Department of Molecular and Cellular Therapeutics, The Royal College of Surgeons in Ireland, Dublin, Ireland
[68] Neurology Department, St. James Hospital, Dublin, Ireland
[69] The Department of Neurology, Beaumont Hospital, Dublin, Ireland
[70] The FutureNeuro Research Centre, Ireland
[71] Neurophysiopathology, Fondazione IRCCS Istituto Neurologico Carlo Besta, Milan, Italy
[72] Unit of Genetics of Neurodegenerative and Metabolic Diseases, Fondazione IRCCS Istituto Neurologico Carlo Besta, Milan, Italy
[73] Department of Pediatric Neuroscience, Fondazione IRCCS Istituto Neurologico Carlo Besta, Milan, Italy
[74] Pediatric Neurology and Muscular Diseases Unit, Department of Neurosciences, Rehabilitation, Ophthalmology, Genetics, Maternal and Child Health, University of Genoa, "G. Gaslini" Institute, Genova, Italy
[75] Laboratory of Neurogenetics, "G. Gaslini" Institute, Genova, Italy
[76] IRCCS, Institute of Neurological Sciences of Bologna, Bologna, Italy
[77] Department of Biomedical and Neuromotor Sciences, University of Bologna, Bologna, Italy
[78] Medical Genetics Unit, Polyclinic Sant'Orsola-Malpighi University Hospital, Bologna, Italy
[79] Department of Biomedical and Neuromotor Sciences, University of Bologna, Bologna, Italy
[80] Institute of Neurology, Department of Medical and Surgical Sciences, University “Magna Graecia”, Catanzaro, Italy
[81] Institute of Molecular Bioimaging and Physiology, CNR, Section of Germaneto, Catanzaro, Italy
[82] Pediatric Neurology, Neurogenetics and Neurobiology Unit and Laboratories, Children's Hospital A. Meyer, University of Florence, Italy
[83] Laboratory for Neurogenetics, RIKEN Center for Brain Science, Saitama, Japan
[84] National Epilepsy Center, Shizuoka Institute of Epilepsy and Neurological Disorder, Shizuoka, Japan
[85] Institute of Biomedical Sciences, Faculty of Medicine, Vilnius University, Vilnius, Lithuania
[86] Centre for Medical Genetics, Vilnius University Hospital Santaros Klinikos, Vilnius, Lithuania
[87] Department of Paediatrics and Child Health, University of Otago, Wellington
[88] Department of Molecular Biology and Genetics, Bogaziçi University, Istanbul, Turkey
[89] Department of Child Neurology, Gulhane Education and Research Hospital, Health Sciences University, Ankara, Turkey
[90] Department of Child Neurology, Istanbul Faculty of Medicine, Istanbul University, Istanbul, Turkey
[91] Department of Child Neurology, Medical School, Dokuz Eylul University, Izmir, Turkey
[92] Department of Child Neurology, Medical School, Kocaeli University, Kocaeli, Turkey
[93] Department of Child Neurology, Medical School, Marmara University, Istanbul, Turkey
[94] Department of Neurology, Istanbul Faculty of Medicine, Istanbul University, Istanbul, Turkey
[95] Department of Genetics, Aziz Sancar Institute of Experimental Medicine, Istanbul University, Istanbul, Turkey
[96] Department of Neurology, Faculty of Medicine, Cerrahpaşa University Istanbul, Istanbul, Turkey
[97] Epilepsy Genetics Program, Department of Neurology, Boston Children's Hospital, Boston, MA 02115, USA
[98] Department of Neurology, Harvard Medical School, Boston, MA 02115, USA
[99] Cooper Medical School of Rowan University, Camden, NJ 08103, USA
[100] Thomas Jefferson University, Philadelphia, PA 19107, USA
[101] The Children's Hospital of Philadelphia, Philadelphia, PA 19104, USA
[102] Perelman School of Medicine, University of Pennsylvania, PA 19104, USA
[103] Nationwide Children's Hospital, Columbus, OH 43205, USA
[104] University of Cincinnati, Cincinnati, OH 45220, USA
[105] Department of Neurology, New York University/Langone Health, New York, NY 10016, USA
[106] University of Montreal, Montreal, QC H3T 1J4, Canada
[107] Beth Israel Deaconess/Harvard, Boston, MA 02115, USA

[108] Department of Neurology, Hofstra-Northwell Medical School, New York, NY 11549, USA
[109] Center for Neuro-engineering and Therapeutics, University of Pennsylvania, Philadelphia, PA 19104 , USA
[110] Department of Neurology, Hospital of University of Pennsylvania, Philadelphia, PA 19104, USA
[111] School of Medicine, Yale University, New Haven, CT 06510, USA
[112] LGS Foundation, NY 11716, USA
[113] National Institute of Neurological Disorders and Stroke, MD 20852, USA
[114] Luxembourg Centre for Systems Biomedicine, University Luxembourg, Esch-sur-Alzette, Luxembourg
[115] Department of Neurology, Sheba Medical Center, Ramat Gan, Israel
[116] Centre for Genomics Research, Precision Medicine and Genomics, IMED Biotech Unit, AstraZeneca, Cambridge, UK
[117] The State Key Laboratory of Brain and Cognitive Sciences, University of Hong Kong, Hong Kong, China
[118] Centre for Genomic Sciences, University of Hong Kong, Hong Kong, China
[119] Population Health and Immunity Division, the Walter and Eliza Hall Institute of Medical Research, Parkville 3052, VIC, Australia
[120] Department of Medical Biology, The University of Melbourne, Melbourne 3010, VIC, Australia
[121] Neuropsychiatry Laboratory, IRCCS Santa Lucia Foundation, Rome, Italy
[122] Division of Neuropsychiatry, Menninger Department of Psychiatry and Behavioral Sciences, Baylor College of Medicine, Houston, TX, USA
[123] IMT School for Advanced Studies Lucca, Lucca, Italy
[124] Department of Neurosciences, Psychology, Drug Research and Child Health, University of Florence, Florence, Italy
[125] Department of Psychiatry, Psychosomatic Medicine and Psychotherapy, University Hospital Frankfurt
[126] Department of Psychiatry, Psychotherapy and Psychosomatics, University Hospital Würzburg
[127] Division of Psychiatry, University College London, London, UK
[128] Division of Psychiatry, Centre for Clinical Brain Sciences, University of Edinburgh, Edinburgh, UK
[129] Institute for Molecular Medicine Finland, University of Helsinki, 00014, Finland
[130] Department of Neurology, Massachusetts General Hospital, Boston, MA, USA
[131] Department of Psychiatry and Behavioral Sciences, SUNY Downstate Medical Center, Brooklyn, NY, USA
[132] Department of Psychiatry, Stony Brook University, Stony Brook, NY, USA
[133] Faculty of Social and Human Sciences, University of Azores, PT
[134] Cherry Health and Michigan State University College of Human Medicine, Grand Rapids, MI, USA
[135] Institute of Medical Psychology, Faculty of Medicine, University of Coimbra, Coimbra, PT
[136] Department of Psychiatry, Wright State University, Dayton, OH, USA
[137] Departments of Psychiatry, Genetics & Genomics, Icahn School of Medicine at Mount Sinai, NY, USA
[138] Semel Institute for Neuroscience at UCLA, Los Angeles, CA, USA
[139] Departments of Public Health and Preventive Medicine, Family Medicine, and Psychiatry and Behavioral Sciences, State University of New York, Upstate Medical University, Syracuse, NY, USA
[140] Department of Psychiatry, University of North Carolina, Chapel Hill, NC, USA
[141] Department of Psychiatry & Behavioral Sciences, University of Southern California, Los Angeles, CA, USA
[142] Department of Psychiatry, Virginia Commonwealth University School of Medicine, Richmond, VA, USA
[143] Department of Psychiatry and Human Behavior, University of California, Irvine, CA, USA
[144] Department of Psychiatry and Behavioral Sciences, Emory University, Atlanta, GA, USA
[145] Department of Psychiatry, Veterans Administration New York Harbor Healthcare System, Brooklyn, NY, USA
[146] Broad Institute of MIT and Harvard, Cambridge, MA, USA
[147] Department of Biology, Massachusetts Institute of Technology, Cambridge, MA, USA
[148] Department of Systems Biology, Harvard Medical School, Boston, MA, USA
[149] Bezmialem Vakif University, Institute of Life Sciences and Biotechnology, Istanbul, Turkey
[150] Department of Genetics, Harvard Medical School, Boston, MA, USA
[151] Department of Neurosurgery, University of Tübingen, Tübingen, Germany

***Details of individual participating Epi25 cohorts***

**Australia: Austin Hospital, Melbourne (AUSAUS)**

The Epilepsy Research Centre at the Austin Hospital in Melbourne, Australia, has been investigating the genetic basis of the epilepsies for over 20 years. The cohort in the Epi25 Collaborative were recruited to the epilepsy genetics research program over this period from the Austin Hospital, epilepsy clinics around Melbourne, and referrals from neurologists Australia-wide. Informed consent was obtained from patients or their parent/guardian as appropriate. DNA was extracted from blood or saliva samples. A skilled team of researchers and clinicians conducted detailed clinical phenotyping which involved a systematic review of medical records, including EEG and MRI reports, and a validated epilepsy questionnaire. Information on family history of seizures and other neurological disorders has also been collected via interviews with the patients and their families. Patients with GGE, non-acquired focal epilepsy, or a DEE were included in Epi25. A very heterogenous collection of epilepsy syndromes are represented in the cohort, including patients with EOAE, CAE, JME and late-onset GGE in the GGE cohort and patients with TLE, FLE, and benign childhood focal epilepsies in the non-acquired focal cohort. The DEE cohort is particularly heterogenous and includes patients with a range of DEE syndromes such as Ohtahara syndrome, Lennox-Gastaut syndrome, epilepsy with myoclonic-atonic seizures, and non-syndromic DEE. An additional subset of patients with lesional focal epilepsy, such as malformations of cortical development or acquired epilepsy, were also included, as well as a selection of patients with familial febrile seizures or FS+.

Most patients in the cohort are of European descent (‘Anglo-Australian’) although there is a diverse range of ethnic backgrounds including Asian, Middle Eastern, Indigenous Australian and mixed ethnicities. There is a known family history of seizures in 46% of the cohort (58% in the subset with GGE). The majority of the patients have had some previous genetic testing, including CNV testing and single gene testing. The DEE cohort have been extensively investigated with multiple iterations of a research panel of known, novel and putative genes for epilepsy. In addition, many patients with focal epilepsy have had a panel of known genes.

**Australia: Royal Melbourne Hospital, Melbourne (AUSRMB)**

The cohort includes patients that have been referred to a seizure clinic. These clinics include the First Seizure Clinic, Video EEG Monitoring Unit and the Epilepsy Clinic based at the Royal Melbourne Hospital. Patients were also referred from the private rooms of neurologists around Victoria.

**Belgium: Antwerp (BELATW)**

Patients were recruited by the Neurogenetics Group of the University of Antwerp through epilepsy clinics at the different university hospitals in Belgium. All patients were diagnosed with a (so far) unexplained presumed genetic epilepsy, and should have had at least 1 MRI of the brain excluding acquired causal lesions.The study was approved by the ethics committee of the University of Antwerp, and parents or the legal guardian of each proband signed an informed consent form for participation in the study. Genomic DNA of individuals was extracted from peripheral blood according to standard procedures. Clinical information was extracted from clinical files, as reported by their treating (paediatric) neurologists, and a subset was reviewed independently by two research team clinicians to ensure data quality and consistency.

**Belgium: Brussels (BELULB)**

Adult patients with epilepsy were recruited consecutively through outpatient clinics and hospitalizations at Hôpital Erasme, Brussels, Belgium (between October 2004 and June 2017) and UZ Gasthuisberg, Leuven, Belgium (between October 2004 and June 2009). The study was approved by the Institutions' Review Boards. All patients provided written informed consent for data collection; patients with learning disability were included after consent from a parent or guardian. DNA was extracted from peripheral blood lymphocytes[1].

**Canada: Andrade (CANUTN)**
There are 96 patients (49 DEE, 3 NAFE, 44IGE) in the Andrade cohort, typically from the Greater Toronto region of Ontario, Canada. There are 44 males and 52 females, mostly of Caucasian ancestry, but also African, South Asian, East Asian, Latino, Middle Eastern, Jewish and Indigenous. Patients were recruited to each group through an REB protocol allowing for the collection of blood or saliva and data collaboration. Patients that were previously consented were reconsented to allow for whole exome sequencing and data sharing with the EPI25 group. After collection, the sample was de-identified, and then extracted and stored at the Hospital for Sick Children, Toronto, Canada.

**Switzerland: Bern University Hospital and University of Bern, Bern (CHEUBB)**
In the recruitment of our cohort, the Departments of Neurology and BioMedical Research, Bern University Hospital and University of Bern, Bern, Switzerland, and the Institute of Human Genetics, Bern University Hospital, Bern, Switzerland, were involved. The Swiss study population encompasses 28 patients with epilepsy between 2 and 63 years of age. All patients have been codified for the Epi25 Study. The patient ascertainment protocol was according to Epi25 phenotyping requirements. Phenotyping information was taken from medical records, stored in the hospital's database, and entered in codified form into a RedCap database provided by Ep25. DNA source was patients' venous blood. DNA was extracted with standard kits at the Institutes of Human Genetics or Clinical Chemistry of Bern University Hospital and stored there at -80 degrees C. Informed consent declarations are available from all patients and have been approved by the American Institutional Review Board involved in the Epi25 Study. The Cantonal Ethics Committee Bern, Switzerland, granted permission for participation of Bern University Hospital and University of Bern in the Epi25 Study including all steps described above. The patient cohort has not been described yet elsewhere.

**Cyprus: The Cyprus Institute of Neurology and Genetics (CYPCYP)**
Epilepsy-affected subjects of the Cyprus cohort were largely recruited and enrolled in the Epi25 Consortium by physicians during routine clinical visits in the Cyprus Institute of Neurology and Genetics. Phenotypic data were collected at the time of enrollment and submitted into the Epi25 RedCap database in a de-identified manner. There are 123 unrelated individuals of Southern European ancestry in the Cyprus cohort, 59 GGE subjects, 53 NAFE and 11 DEE. All subjects selected for this study had clinical, neuroimaging and EEG or video-EEG characteristics meeting the International League against Epilepsy (ILAE) 2017 Seizure Classification. The controls cohort consisted of a group of 32 individuals. of Southern European ancestry and were not diagnosed with epilepsy or other neuropsychiatric phenotypes. Genomic DNA samples were extracted from whole blood with the Gentra Puregene Blood Kit (Qiagen, Hilden, Germany) according to the manufacturer's guidelines. This study was carried out in compliance with the Cyprus National Bioethics Committee (ΕΕΒΚ/ΕΠ/2015/22). Written informed consent was obtained from all study participants or their legal guardians at the Cyprus Institute of Neurology and Genetics.

**Czech Republic: University Hospital Motol, Prague (CZEMTH)**
Patients in our cohort have been diagnosed with West syndrome, myoclonic-astatic epilepsy or developmental and epileptic encephalopathy of unknown aetiology. Brain magnetic resonance imaging and metabolic screening excluded any underlying pathology. Patients were collected at the Department of Child Neurology of the 2nd Medical Faculty and University Hospital Motol.

Legal guardians of patients signed an informed consent. The study was approved by the local ethics committee.

**Germany: Epilepsy Center Frankfurt Rhine-Main, Goethe University, Frankfurt, and Epilepsy Center Hessen, Philipps University, Marburg (DEUPUM)**
The principal investigators (Karl Martin Klein, Felix Rosenow, Philipp S. Reif, and Susanne Knake) contributed 259 samples. The patients were recruited from the outpatient clinics and the video EEG monitoring units at the Epilepsy Centers Frankfurt Rhine-Main and Hessen-Marburg and contained 64 samples with genetic generalized epilepsy, 130 samples with non-acquired focal epilepsy, 60 samples with lesional focal epilepsy and 4 samples with DEE. All patients were phenotyped in detail by epilepsy specialists (KMK, FR, SK, PSR) within the EpimiRNA project (European Union's 'Seventh Framework' Programme (FP7) under Grant Agreement no. 602130). EEGs and MRIs were performed as part of the clinical workup. Phenotypic classification and data entry for the biobank for paroxysmal neurological disorders was performed by KMK. Patient selection and data export from the biobank for the study was performed by KMK. DNA was extracted from peripheral blood or saliva. All patients provided written informed consent.

**Germany: University of Bonn, Bonn (DEUUKB)**
The sample recruitment site is the Department of Epileptology at the University of Bonn. The collection of blood DNA samples from patients with epilepsy was conducted from 2007 till 2015 within the studies Epicure (Functional Genomics in Neurobiology of Epilepsy: A Basis for New Therapeutic Strategies) and NGEN-Plus (Genetic basis of Levetiracetam pharmacoresistance and side effects in human epilepsy) and has been approved by the Ethics committee of University Bonn Medical Center (040/07). Genomic DNA was isolated from 10 ml aliquots of EDTA-anticoagulated blood by a salting-out technique[2]. From selected samples of this cohort GWAS data have been published in several studies[3-6].

**Germany: University Hospital Schleswig-Holstein, Kiel (DEUUKL)**
Patients were recruited by the Neuropedriatics Group of the University Hospital of Schleswig-Holstein and through the Israeli-Palestinian Family Consortium. The recruitment and analysis of these samples is covered by the Kiel IRB. Patients, their parents or the legal guardian of each proband signed an informed consent form for participation in the study. Clinical data was collected from clinical files and a subset of patients from Israel or Palestine was interviewed by a research team of clinicians to provide their clinical data. Genomic DNA of patients was extracted from peripheral blood according to standard procedures.

**Germany: TLE Leipzig (DEUULG)**
Patients were recruited by the Swiss Epilepsy Center in Zurich, Switzerland and samples were transferred for research and storage to the Institute of Human Genetics at the University of Leipzig, Germany. All patients were diagnosed with temporal lobe epilepsy due to an indicative EEG. Most patients had at least 1 MRI of the brain with focus on focal abnormalities, especially of the temporal lobe / hippocampal structures. The study was approved by the "Kantonale Ethikkommission Zürich". Parents or the legal guardian of each proband signed an informed consent form for participation in research studies including whole genome analyses. Genomic DNA of individuals was extracted from peripheral blood according to standard procedures. Clinical information was extracted from clinical files, as reported by their treating neurologists.

**Germany: University of Tübingen, Tübingen (DEUUTB)**
Our study cohort consists of over 1000 samples with mainly Caucasian origin. These samples were recruited at Tubingen and 38 other cooperating departments of neurology of university clinics and outpatient clinics in Germany. The Ethics / informed consent was approved by the

ethics committee of the Medical Faculty of the University and the University Clinic of Tübingen. Patients with the diagnosis of genetic generalized epilepsies, epileptic encephalopathies and non-acquired focal epilepsies were systematically recruited by a letter of invitation. After approving the informed consent by the participants, the data were collected retrospectively from medical reports and in exceptional cases, by personal interviews of patients or their relatives. The DNA source was blood.

**Finland: Kuopio University Hospital, Kuopio (FINKPH)**
Patients diagnosed with epilepsy and visiting Epilepsy Center, Kuopio University Hospital, Finland have given their written informed consent to record their clinical data to a research registry and collect a blood sample for DNA analysis. The ethics committee of Kuopio University Hospital has approved the study.

**Finland: University of Helsinki, Helsinki (FINUVH)**
Patients were recruited at the University of Helsinki through pediatric neurology clinics in the Helsinki and Tampere University Hospitals in Finland. All patients were diagnosed with a presumed genetic epilepsy, the etiology remaining unknown. All patients had an MRI done to exclude acquired causal lesions. The study was approved by an ethics committee of The Hospital District of Helsinki and Uusimaa, Finland. The parents or the legal guardian of each proband has signed an informed consent form. Genomic DNA of the patients was extracted from peripheral blood or saliva according to standard procedures. Clinical information was extracted from clinical files, as reported by the treating pediatric neurologists.

**Wales: Swansea (GBRSWU)**
The samples from Wales: Swansea are recruited from regional NHS HealthBoard Clinics and consented into the IRAS-ethical permissions framework of the Swansea Neurology Biobank (17/WA/0290). Informed consent is given for samples to be used for research purposes and for third-party consortia with appropriate MTA agreements. Patents attend epilepsy or general neurology clinics and are prioritised if they reach the clinical evidence for submission to studies. Blood samples are sent to the UK Porton Down EcACC facility and returned to SNB in batches where they are checked and validated through gender testing. The samples submitted to Epi25 must pass the DNA QC standards for NGS pipelines and have the level of certainty for clinical diagnosis.

**UK: University College London, London (GBRUCL)**
Participant recruitment took place at the National Hospital for Neurology and Neurosurgery (United Kingdom). Written informed consent or assent was obtained between 10/01/2000 and 01/25/2015 from all participants according to local and national requirements and blood samples were collected for DNA extraction. 709 epilepsy cases were submitted for analysis. Allocation to the following groups was based on the clinical diagnosis and the specific inclusion and exclusion criteria of the Epi25 consortium: generalized genetic epilepsy (n=393, 145 male), non-acquired focal epilepsy (n=313, 146 male), developmental and epileptic encephalopathy (n=3, 2 male). Additionally, relatives were included, where samples were available (n=3, 2 male). Phenotypic information was obtained from local medical records by clinical or trained non-clinical researchers.

**UK: University of Liverpool, Liverpool and Imperial College London, London (GBRUNL)**
GBRUNL samples are derived from four separate, UK-wide, ethically approved studies coordinated by the University of Liverpool (UK) and Imperial College London (UK). The SANAD and MESS linked DNA Bank and Relational Database study recruited individuals with newly-diagnosed focal, generalised or unclassified epilepsy from out-patient neurology clinics between 2003-2006[7; 8]. The Pharmacogenetics of GABAergic Mechanisms of Benefit and Harm in Epilepsy

study recruited patients with refractory focal epilepsy, previously or prospectively exposed to adjunctive treatment with clobazam or vigabatrin, from out-patient neurology clinics between 2005-2009. The Refractory Juvenile Myoclonic Epilepsy Cohort (ReJuMEC) study recruited individuals with valproic acid resistant juvenile myoclonic epilepsy from out-patient neurology clinics between 2009 and 2010. The ongoing Standard and New Antiepileptic Drugs (SANAD-II) study, which is recruiting individuals with newly-diagnosed focal, generalised or unclassified epilepsy from out-patient neurology clinics between 2013-2019. In all cases, study participants provided written informed consent to the collection (via blood or saliva sampling) and analysis of their DNA for use in genetic and pharmacogenetic research related to epilepsy and its treatment. All studies were approved by research ethics committees in operation at the relevant time (SANAD DNA bank, North West MREC ref 02/8/45; GABAergic mechanisms, UCLH REC ref 04/Q0505/95; ReJuMEC, Cheshire REC ref 09/H1017/55; SANAD-II, North West REC ref 12/NW/0361). Assembly of the GBRUNL cohort was supported by generous funding from The Wellcome Trust, the Imperial College NIHR Biomedical Research Centre, the Department of Health (UK), the Medical Research Council (UK), and the National Institute of Health Research (UK).

**Hong Kong: Chinese University of Hong Kong (HKGHKK)**
Epilepsy patients of Han Chinese ethnicity aged between 2 and 91 years were recruited from neurology clinics of five regional hospitals in Hong Kong covering a combined catchment population of approximately 3 million. Syndromic classification was adapted from the revised international organization of phenotypes in epilepsy. DNA was extracted from venous blood. The study was approved by ethics committees of the participating hospitals, and all patients or their legal guardians gave written informed consent. The sample collection methodology has been described previously[9].

**Croatia: University Clinical Centre Zagreb, Zagreb (HRVUZG)**
Pediatric patients were recruited from University Medical Centre Zagreb and 2 patients from 2 other epilepsy clinics in Croatia. All patients were diagnosed as possible genetic epilepsy not yet explained. All patients underwent MR brain imaging at least once, the acquired epilepsy causes were excluded. DNA was extracted from peripheral blood according to the accepted protocol. The study was approved by Hospital ethical committee and all parents or legal guardian of probands signed informed consent for participation in the study. Clinical information was extracted from clinical files. The cohort was also reviewed by reviewed by collaborative research team clinicians from University of Antwerp to ensure data quality and consistency.

**Ireland: Dublin (IRLRCI)**
Patients were recruited from a specialized epilepsy clinic at Beaumont Hospital and St. James' Hospital, Dublin, Ireland. Patients were mostly of Irish ethnicity. DNA was extracted from a combination of lymphocytes and saliva. All participants provided written informed consent.

**Italy: Fondazione IRCCS Istituto Neurologico Carlo Besta, Milan (ITAICB)**
Fondazione IRCCS Istituto Neurologico Carlo Besta, Milan, Italy (ITAICB): Our cohort included the DNA samples of 207 patients and 62 controls (healthy subjects not related with the patients, and without epilepsy history). The patient population included 106 patients with generalized epilepsy (GGE), 51 patients with developmental and epileptic encephalopathy (DEE), 33 patients with focal epilepsy due to cerebral malformations (nodular heterotopia), and 15 patients with non-acquired focal epilepsy (NAFE). All of the patients were diagnosed and followed at our Institute. Diagnosis of epilepsy was based on clinical, EEG and neurophysiological data, neuroimaging (MRI). Metabolic screening, karyotype, CGH array, analyses of single genes and customized panels were performed in some cases, when appropriate. The patients did not undergo to exome

sequencing analysis (before the Epi25 collection). The DNA of the patients was extracted from peripheral blood, according with standard procedures, after signature of an Informed Consent form. The genetic study was approved by The Ethic Committee of our Institute. No publications have described genetic findings pertaining to the collected patients until now. Clinical information was extracted from clinical files, as reported by their treating (paediatric and adult) neurologists.

**Italy: Gaslini Institute, Genova (ITAIGI)**
Patients with generalized and focal epilepsy or developmental and epileptic encephalopathy followed- up or referred for genetic analysis at 'Gaslini Institute'. The study was approved by the IRB and written informed consent was signed by the patients/parents. Clinical information, including data on EEG and antiepileptic therapy, were recorded on data collection forms. Genomic DNA isolation and genetic analysis was carried out with the Nimblegen-SeqCapEZ-V244M enrichment kit on the Illumina HiSeq2000 system.

**Italy: IRCCS Institute of Neurological Science of Bologna, Bologna (ITAUBG)**
Patients were recruited by the Adult and Pediatric Neurologists of the IRCCS Institute of Neurological Sciences, Bellaria Hospital, Bologna. Patients with generalized epilepsy, focal epilepsy (with or without brain lesions) and developmental and epileptic encephalopathy were referred by epilepsy clinics. All patients were diagnosed with a (so far) epilepsy of uncertain aetiology and underwent neuro-radiological imaging (CT or MRI) and EEG. The study was approved by the local ethics committee (cod. CE:16057). Patients themselves or parents or the legal guardian of each proband signed an informed consent form for the study participation. Genomic DNA of individuals was extracted from peripheral blood according to standard procedures. Clinical information was collected from medical records, as reported by their treating neurologists[10; 11].

**Italy: University Magna Graecia, Catanzaro (ITAUMC)**
Patients were recruited by the Epilepsy Group of the University Magna Graecia of Catanzaro (Italy) that includes a Pediatric and Adult Neurologic Unit with a specific focus on genetic epilepsy. In each patient, the diagnosis of epilepsy syndrome is based on comprehensive clinical, neuropsychological, electroencephalographic, and MR evaluations. Clinical data are stored into a database. The study was approved by the ethics committee of the University of Catanzaro Italy, and parents or the legal guardian of each proband signed an informed consent form for participation in the study. Genomic DNA of individuals was extracted from peripheral blood according to standard procedures

**Italy: Meyer Hospital, Florence (ITAUMR)**
Patients were recruited by the Neurology and Neurogenetics Group of the Meyer Hospital of Florence. All patients were diagnosed with a unexplained presumed genetic epilepsy. The study was approved by the Pediatric Ethics Committee of the Regione Toscana, and parents or the legal guardian of each proband signed an informed consent form for participation in the study. Genomic DNA of individuals was extracted from peripheral blood according to standard procedures. Clinical information was extracted from clinical files, as reported by their treating (paediatric) neurologists.

**Japan: RIKEN Institute, Tokyo (JPNRKI)**
Japanese patients with epilepsies were recruited by National Epilepsy Center, Shizuoka Institute of Epilepsy and Neurological Disorder. Epileptic seizures and epilepsy syndrome diagnoses were performed according to the International League Against Epilepsy classification of epileptic syndromes. Genomic DNA was extracted from peripheral venous blood samples using QIAamp DNA Blood Midi Kit according to manufacturer's protocol (Qiagen). The experimental protocols

were approved by the Ethical Committee of RIKEN Institution and National Epilepsy Center, Shizuoka Institute of Epilepsy and Neurological Disorder. Written informed consent was obtained from all individuals and/or their families in compliance with the relevant Japanese regulations.

**Lithuania: Vilnius University Hospital Santaros Klinikos, Vilnius (LTUUHK)**
Patients were recruited in Vilnius University Hospital Santaros Klinikos by a clinical geneticist through a referral of a neurologist or a child neurologist and according to inclusion/ exclusion criteria. All patients were diagnosed with one of the three forms of epilepsy - genetic generalized, focal non-lesional or developmental and epileptic encephalopathy - after exclusion of acquired causes and should have had at least 1,5T brain MRI and a diagnostic EEG. The study was approved by Vilnius Regional Biomedical Research Ethics Committee, and each proband or parents/ legal guardians of a proband signed an informed consent form for participation in the study. Samples of genomic DNA were obtained during the routine procedure for blood sampling for genetic testing done in a clinical testing and the majority of patients had chromosomal microarray, metabolic testing and/or gene/gene panel testing prior to the inclusion into the study. Clinical information was extracted from clinical files and obtained during the clinical genetic consultation.

**New Zealand: University of Otago, Wellington (NZLUTO)**
Cases were recruited as part of a larger study from neurology, paediatric and genetic outpatient services throughout New Zealand. Participants were between 1 month and 63 years of age from the following ethnic groups: New Zealand European, Māori, Pasifika, Asian, Hispanic, Ethiopian. Using a structured interview and review of medical records diagnosis was based on the International League of Epilepsy (ILAE) classification and made by a paediatric neurologist. The study protocol was approved by the New Zealand Health and Disability Ethics Committee. Participants gave written informed consent for clinical and genetic analysis. DNA was extracted from blood or saliva.

**Turkey: Bogazici University, Istanbul (TURBZU)**
Patients were recruited by the Child Neurology and Neurology clinics at the different university hospitals in Turkey. The study was approved by the Institutional Review Board for Research with Human Subjects (INAREK) of Boğaziçi University, and parents or the legal guardian of each proband signed an informed consent form for participation in the study. Genomic DNA of individuals was extracted from peripheral blood according to standard procedures. Clinical information was reported by their treating (pediatric) neurologists. All developmental and epileptic encephalopathy patients had severe epilepsy, with developmental delay and regression, normal neuroimaging and epileptiform activity on EEG. Turkish population control group included individuals with no symptoms of any neurological disorder.

**Turkey: Istanbul University, Istanbul (TURIBU)**
Epilepsy patients were recruited from Epilepsy Clinic, Department of Neurology, Istanbul Faculty of Medicine, Istanbul University. The study population consisted of patients with idiopathic/genetic generalized epilepsies, lesional or non-lesional focal epilepsies and epileptic encephalopathies, including sporadic and familial cases. All patients were long-term follow-up. Seizure types, age of onset, neurological examinations, past and family history, prognosis and response to treatment, features of electroencephalography and neuroimaging were evaluated. Ethics committee approval was obtained. Peripheral blood samples were collected from all patients following written informed consent. DNA isolation was performed in Department of Genetics, Aziz Sancar Institute of Experimental Medicine, Istanbul University.

**USA: Boston Children's Hospital, Boston (USABCH)**

Cases from Boston Children's Hospital (BCH) were ascertained from 3 local repositories. All repository protocols are approved by the BCH Institutional Review Board and participants were consented under one (or more) of the following protocols. The Genetics of Epilepsy and Related Disorders protocol, led by Dr. Annapurna Poduri, enrolls patients with a clinical epilepsy diagnosis for genotype/phenotype correlation. Samples are obtained from BCH and non-BCH patients and biological samples collected for genetic sequencing. Patient medical records (BCH and outside records) are reviewed for phenotyping purposes[12]. The Phenotyping and Banking Repository of Neurological Disorders is a local repository led by Dr. Mustafa Sahin. BCH patients with any neurological phenotype, including epilepsy, are enrolled and biological samples collected. Boston Children's Biobank for Health Discovery is a local repository led by Dr. Kenneth Mandl that enrolls any patient of BCH, regardless of diagnosis or phenotype. Samples from these two broader repositories are available to BCH researchers through an application process, including a supporting IRB-approved protocol. Patients with a clinical diagnosis of epilepsy were reviewed for Epi25 eligibility using their BCH medical records.

**USA: Philadelphia/CHOP (USACHP) and Philadelphia/Rowan (USACRW)**
The Philadelphia Cohort began in 1997 and collected blood, saliva and brain tissues from patients with common forms of idiopathic human epilepsy, mostly genetic generalized epilepsy (GGE) and non-acquired focal epilepsy (NAFE). The collection began at Thomas Jefferson University Hospital in Philadelphia and expanded to include six other sites: The Children's Hospital of Philadelphia, The University of Pennsylvania, The University of Cincinnati, Nationwide Children's Hospital, Beth Israel Deaconess and The University of Montreal. The cohort consists of 2615 samples from epilepsy patients collected and supported during two periods of NIH funding (R01NS493060, 2001-2007 RJ Buono PI and R01NS06415401, 2009-2012 RJ Buono and H Hakonarson Co- PI). Over 1000 additional samples from first degree relatives of the patients were also collected. Many of these samples are available to the research community via the NINDS sample repository at the Coriell Institute in Camden NJ. All studies were approved by Institutional Review Boards at each participating site. All patients were identified and recruited by trained epileptologists at tertiary care centers using inclusion and exclusion criteria previously published. Diagnostic methods applied included EEG, MRI, and collection of deep phenotypic information on family history, medications, risk factors, age of onset, and other information. Inclusion and exclusion criteria for the cohort were published previously[6; 13]. For the Epi25K project, blood and saliva were used as the source of DNA.

**USA: Epilepsy Phenome/Genome Project (USAEGP)**
Infantile spasms (IS), Lennox–Gastaut syndrome (LGS), genetic generalized epilepsy (GGE), and non-acquired focal epilepsy (NAFE) patients were collected through the Epilepsy Phenome/Genome Project[14] (EPGP, http://www.epgp.org). More than 4,000 participants in EPGP were enrolled across 27 clinical sites from around the world. The subset of samples included in Epi25 were enrolled from 20 sites across the USA and in Australia. IS patients were required to have hypsarrhythmia or a hypsarrhythmia variant on EEG. LGS patients were required to have EEG background slowing or disorganization for age and generalized spike and wave activity of any frequency or generalized paroxysmal fast activity (GPFA). IS and LGS cases were enrolled as trios with both biological parents. Participants with NAFE and GGE were required to have a first degree relative who also had NAFE or GGE (did not have to be concordant). All patients had no confirmed genetic or metabolic diagnosis, and no history of congenital TORCH infection, premature birth (before 32 weeks gestation), neonatal hypoxic-ischaemic encephalopathy or neonatal seizures, meningitis/encephalitis, stroke, intracranial haemorrhage, significant head trauma, or evidence of acquired epilepsy. Enrollment required detailed confirmation of detailed phenotypic data including medical record review and abstraction, patient

interviews, EEG and MRI, and comprehensive review by expert scientific cores for EEG, MRI, and clinical final diagnosis.

**USA: NYU Human Epilepsy Project (USAHEP)**
Participants were recruited for the Human Epilepsy Project at 33 different medical centers located in US, Canada, Australia, Austria, Finland, and Ireland. All participants were between 12 and 60 at the age of enrollment and had a clinical history consistent with a diagnosis of focal epilepsy, as determined by an eligibility panel of epilepsy specialists. Participants were required to have two or more spontaneous seizures with clinically observable features in the past 12 months, and 4 or fewer months of anticonvulsant treatment. Those with major medical comorbidities, intellectual disability, or significant psychiatric disease were excluded, as were those with progressive neurological lesions on imaging or known neurodegenerative disease. Participants completed daily electronic diaries tracking seizures, medication adherence, and mood. Mood and cognition were assessed periodically via standardized instruments, and brain MRIs and EEGs were obtained for all participants. Blood was collected and banked annually, allowing for study of DNA, RNA and protein. HEP was approved by the IRBs at all participating sites, and all participants or their parent/legal guardian gave written informed consent. Minors also gave written assents. HEP was funded by the Epilepsy Study Consortium.

**USA: Penn/CHOP, Philadelphia (USAUPN)**
The USAUPN cohort was recruited at the Children's Hospital of Philadelphia (CHOP) and Hospital of the University of Pennsylvania (UPenn), including pediatric and adult patients with epilepsies through dedicated IRB protocols at CHOP and UPenn. Patients were recruited in an inpatient and outpatient setting and samples were also contributed through the Penn Biobank.

***Details of individual control cohorts (not including samples obtained from dbGaP)***

**Italian controls (PI: Spalletta)**
Right-handed healthy individuals were recruited and had whole blood drawn by local advertisements at Santa Lucia Foundation in Rome, Italy. All of the individuals were born and educated in Italy and had Italian-Caucasian ancestry, to reduce the possibility of artifactual association caused by ethnic stratification. Exclusion criteria were: (i) major medical illnesses and/or known or suspected history of alcoholism or drug dependency and abuse; (ii) mental disorders (i.e. schizophrenia, mood, anxiety, personality and/or any other significant mental disorders) according to the DSM-IV-TR criteria assessed by the Structured Clinical Interviews for DSM-IV-TR [SCID-I and SCID-II] and/or neurological disorders diagnosed by an accurate clinical neurological examination; (iii) presence of vascular brain lesions, brain tumour and/or marked cortical and/or subcortical atrophy on magnetic resonance imaging (MRI) scan; and (iv) suspicion of cognitive impairment or dementia based on Mini Mental State Examination (MMSE) scores ≤24 (a cut-off point for dementia screening in the Italian population)[15] and confirmed by a clinical neuropsychological evaluation using the Mental Deterioration Battery[16] and the NINCDS-ADRDA criteria for dementia[17]. The presence of anxiety symptoms was assessed using the Hamilton Rating Scale for Anxiety (HAM-A)[18]. Written, informed consent was obtained from all subjects participating in the study, which was approved by the local ethics committee at the Santa Lucia Foundation of Rome (protocol number CE/11.9)[19].

**German controls**
Subjects have been recruited at the Department of Psychiatry and Psychotherapy, University of Würzburg, Germany, with the exception of the TK samples (n=63, they are anonymous blood donors). All subjects have been screened for the absence of mental disorders (by MINI) as well

as severe medical and neurological (including epilepsy) disorders (by self-report). Ethnicity is Caucasian by self-report in all cases, and DNA source is blood. Studies were approved by the IRB, University Hospital Würzburg; all participants gave written informed consent.

**UK/IRL controls 1 (PI: McQuillin)**
The UCL control sample consisted of 480 genomic DNA samples that were extracted from EBV transformed peripheral blood lymphocytes from unscreened healthy British blood donors (https://www.phe-culturecollections.org.uk/products/dna/hrcdna/hrcdna). The remaining DNA samples were extracted from whole blood samples from healthy volunteers of UK or Irish ancestry who were interviewed with the initial clinical screening questions of the SADS-L and selected on the basis of not having a past or present personal history of any RDC-defined mental disorder. Heavy drinking and a family history of schizophrenia, alcohol dependence or bipolar disorder, were also used as exclusion criteria for controls. UK National Health Service multi-centre and local research ethics approvals were obtained and all subjects gave signed informed consent[20].

**UK/IRL controls 2 (PIs: McIntosh, Blackwood, and Johnstone)**
Participants were recruited from clinical service around Edinburgh and Scotland and screened using the SADS-L. DNA samples were extracted from whole blood for genotyping and sequencing. Research was conducted after research ethics and NHS management approvals[21].

**FINRISK controls**
The controls from FINRISK that contributed to the Epi25 WES study were part of the FINRISK inflammatory bowel disease (IBD) cohort. The population-based FINRISK study have been followed up for IBD and other disease end-points using annual record linkage with the Finnish National Hospital Discharge Register, the National Causes-of-Death Register and the National Drug Reimbursement Register. Controls were chosen to have a high polygenic risk score for IBD without an IBD diagnosis. A detailed description of the FINRISK cohort can be found at Borodulin *et al*[22].

**Genomic Psychiatry Cohort (GPC) controls**
The controls from GPC that contributed to the Epi25 WES study were a subset of the overall control participants of European ancestry with no personal or family history of schizophrenia or bipolar disorder. All the samples were exome-sequenced at the Broad Institute. A detailed description of the GPC cohort can be found at Pato *et al*[23].

### *Supplemental acknowledgments*

Additional funding sources and acknowledgment of individual patient cohorts are as follows: The **AUSAUS** cohort was supported by NHMRC Program Grant (ID: 1091593). Effort at **AUSRMB** was supported by NHMRC Project Grant (#1059858) and NHMRC Program Grant (#1091593). H.S. from the **BELATW** cohort is a PhD fellow of the Fund for Scientific Research Flanders (FWO, 1125416N); and S.W. receives funding support from the BOF-University of Antwerp (FFB180053) and FWO (Senior Clinical Investigator Fellowship 1861419N). Funding that supported activities at the **BELULB** site included Fonds National de la Recherche Scientifique, Fonds Erasme, Université Libre de Bruxelles, and European Commission grant 279062. Work conducted at **CANUTN** was supported by funding from Ontario Brain Institute – EpLink. The **CHEUBB** cohort was supported by directorate for Public Health and Social prevention of the Canton of Bern: Medical Innovation “next-generation-sequencing epilepsy” 2015. Work at the **CYPCYP** site was funded by TELETHON/33173158. Effort at **CZEMTH** was supported by Ministry of Health of Czech Republic AZV 15–33041 and DRO 00064203. At **DEUPUM**, all patients were phenotyped in detail by epilepsy specialists (KMK, FR, SK, PSR) within the EpimiRNA project (European Union's 'Seventh Framework' Programme (FP7) under Grant Agreement no. 602130). KMK, FR and PSR were supported by the Detlev-Wrobel-Fonds for Epilepsy Research Frankfurt. At **DEUUKB**, sample recruitment of the University Bonn cohort was funded by the European Community (FP7 project EpiPGX, grant 279062 to WSK). The work at **USAUPN** and **DEUUKL** was supported by intramural funds of the Children's Hospital of Philadelphia, the German Research Foundation (HE5415/3-1 to I.H.) within the EuroEPINOMICS framework of the European Science Foundation, the German Research Foundation (DFG, HE5415/5-1, HE5415/6-1, HE5415/7-1 to I.H.), and the Genomics Research and Innovation Network (GRIN, grinnetwork.org), a collaboration amongst the Children's Hospital of Philadelphia, Boston Children's Hospital, Cincinnati Children's Hospital Medical Center. I.H. also received support through the International League Against Epilepsy. This work was also partially supported by NIH grant NS108874. Further funding for the **DEUUTB** cohort was obtained from the German Research Foundation and the Fond Nationale de la Recherche in Luxembourg in the frame of the Research Unit FOR-2715 (grants Le1030/16-1, Kr5093/2-1, We4896/4-1, He5415/7-1), by the German Society for Epileptology, and by the foundation no-epilep. The **FINKPH** cohort was supported by funding from the Saastamoinen Foundation, Vaajasalo Foundation and Kuopio University Hospital State Research. Effort at **FINUVH** was supported by Folkhälsan Research Foundation, Helsinki Finland and Foundation for Pediatric Research, Helsinki, Finland. Funding for the **GBRSWU** cohort included Health and Care Research Wales, Wales Gene Park, The BRAIN Unit. The **GBRUCL** cohort thanks the Epilepsy Society, Muir Maxwell Trust and Wellcome Trust Strategic Award (WT104033AIA) for funding support. The work conducted at **GBRUCL** was undertaken at University College London Hospitals, which received a proportion of funding from the NIHR Biomedical Research Centres funding scheme. Assembly of the **GBRUNL** cohort was supported by generous funding from The Wellcome Trust, the Imperial College NIHR Biomedical Research Centre, the Department of Health (UK), the Medical Research Council (UK), and the National Institute of Health Research (UK). The **HKGHKK** cohort was supported by Research Grants Council of the Hong Kong Special Administrative Region, China (project number CUHK4466/06M). The **HRVUZG** cohort acknowledges Sarah Weckhuysen and Hannah Stamberger for help with sample shipping. The work at **IRLRCI** was supported by research grants from Science Foundation Ireland (SFI) (16/RC/3948 and 13/CDA/2223) and co-funded under the European Regional Development Fund and by FutureNeuro industry partners. Effort at **ITAICB** was supported by European project, Grant agreement 602531 (DESIRE) (Silvana Franceschetti only was involved) The **ITAIGI** cohort thanks the Genetic Commission of Italian League Against Epilepsy (LICE) for its contribution to this work. The **ITAUBG** cohort thanks Dr Baldassari S; Dr Caporali L and Prof Carelli V and all the staff of the Laboratory of Neuro-genetics in Bologna; and Dr Margherita Santucci and the Child Neurology Unit, Bellaria Hospital in Bologna, as well as all physicians and nurses of the Epilepsy Centre from the Bellaria Hospital in Bologna. Work at **ITAUMR** was supported by the European Commission Seventh Framework Programme under the project DESIRE (grant agreement no. 602531) (to R.G.). Research at the **JPNRKI** cohort was (partly) supported by AMED under Grant Number JP18dm0107092 and JP18ek0109388 (K.Y.). Effort at **NZLUTO** was supported by Health Research Council of New Zealand: 15/070 – Gene Discover in Epilepsy: the building block of precision medicine; Cure Kids New Zealand: 3576 – Genetics of Epilepsy. The **TURBZU** cohort was supported by Bogazici University Research Fund, Project No: 17B01P3. **USACHP** and **USACRW** were supported by NIH funding (R01NS493060, 2001-2007 RJ Buono PI and R01NS06415401, 2009-2012 RJ Buono and H Hakonarson Co- PI). U01HG006830 (PI Hakonarson) and The Children's Hospital of Philadelphia Endowed Chair in Genomic Research (PI Hakonarson). Work at **USAEGP** was supported by National Institute of Neurological Diseases and Stroke (NINDS) grant U01 NS053998, as well as planning grants from the Finding a Cure for Epilepsy and Seizures (FACES) Foundation and the Richard Thalheimer Philanthropic Fund. At **USAHEP**, H.E.P is sponsored by The Epilepsy Study Consortium (ESCI).

ESCI is a non-profit organization dedicated to accelerating the development of new therapies in epilepsy to improve patient care. The funding provided to ESCI to support HEP comes from industry, philanthropy and foundations (UCB Pharma, Eisai, Pfizer, Lundbeck, Sunovion, The Andrews Foundation, The Vogelstein Foundation, Finding A Cure for Epilepsy and Seizures (FACES), Friends of Faces and others). C.A.E. from the **USAUPN** cohort was supported in part by a Ruth L. Kirschstein National Research Service Award (NRSA) Institutional Research Training Grant, T32 NS091008-01.

Additional funding sources and acknowledgment of individual control cohorts are as follows: Collection of the **Italian controls** was supported by RF-2013-02359074; NET-2011-02346784 (Italian Ministry of Health) and was sequenced as part of the CCDG funding. Sequencing of the **German controls** was funded by the Dalio Foundation DFG grant TRR-CRC 58, Z02. The **UK/IRL controls** were collected and sequenced with support from the Neuroscience Research Charitable Trust, the Central London NHS (National Health Service) Blood Transfusion Service and the National Institute of Health Research (NIHR) funded Mental Health Research Network. The **FINRISK controls** were part of the FINRISK studies supported by THL (formerly KTL: National Public Health Institute) through budgetary funds from the government, with additional funding from institutions such as the Academy of Finland, the European Union, ministries and national and international foundations and societies to support specific research purposes. The **GPC controls** were sequenced with funding from NIH grants MH085548 and MH085542 and the Stanley Center for Psychiatric Research, Broad Institute of MIT and Harvard. Public controls used for the analysis were obtained from dbGaP at http://www.ncbi.nlm.nih.gov/gap. Controls from the MIGen studies were obtained through dbGaP accession numbers phs000806.v1.p1, phs001000.v1.p1, and phs000814.v1.p1. We thank the Broad Institute for generating high-quality sequence data for the MIGen studies supported by NHGRI funds (grant # U54 HG003067) with Eric Lander as PI. The Sweden-Schizophrenia (SCZ) controls were obtained through dbGaP accession number phs000473.v2.p2. These samples were provided by the Swedish Cohort Collection supported by the NIMH grant R01MH077139, the Sylvan C. Herman Foundation, the Stanley Medical Research Institute and The Swedish Research Council (grants 2009-4959 and 2011-4659). Support for the Swedish-SCZ exome sequencing was provided by the NIMH Grand Opportunity grant RCMH089905, the Sylvan C. Herman Foundation, a grant from the Stanley Medical Research Institute and multiple gifts to the Stanley Center for Psychiatric Research at the Broad Institute of MIT and Harvard.

### Supplemental Figures

(A) Raw data

(B) Pre-filtered

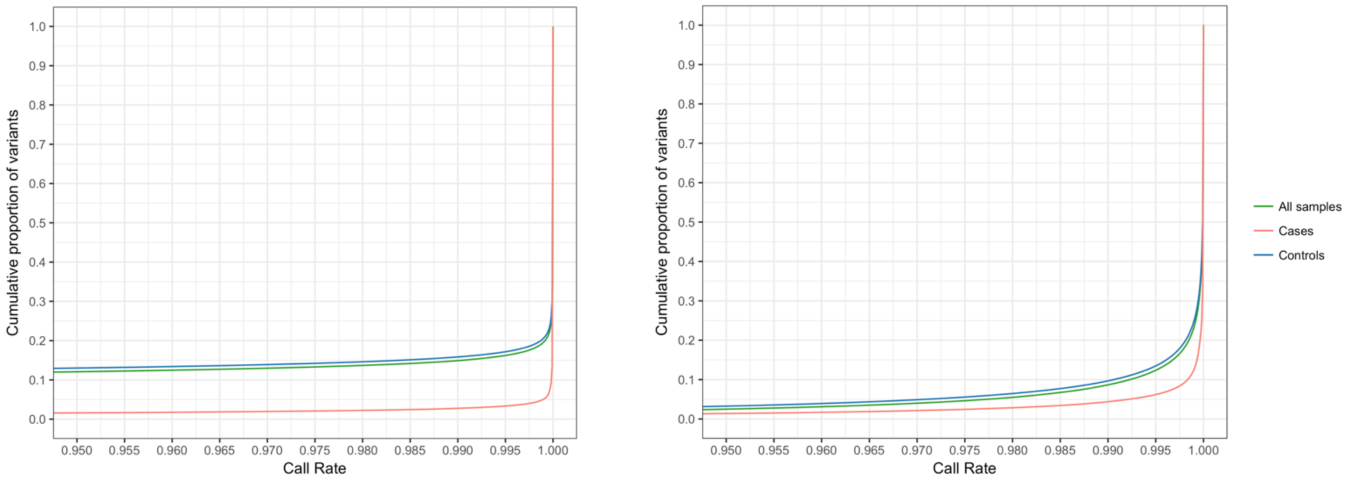


**Figure S1.** Variant pre-filtering: restricting to well-covered exome intervals across platforms reduced variant calling bias due to capture difference

As the first step of the QC process, we removed sites that fell outside regions with at least 10x coverage in 80% of the samples across sequencing batches and platforms (Illumina and Agilent), regardless of targeted capture intervals. This pre-filtering effectively minimized the call rate difference between cases and controls.

(A)

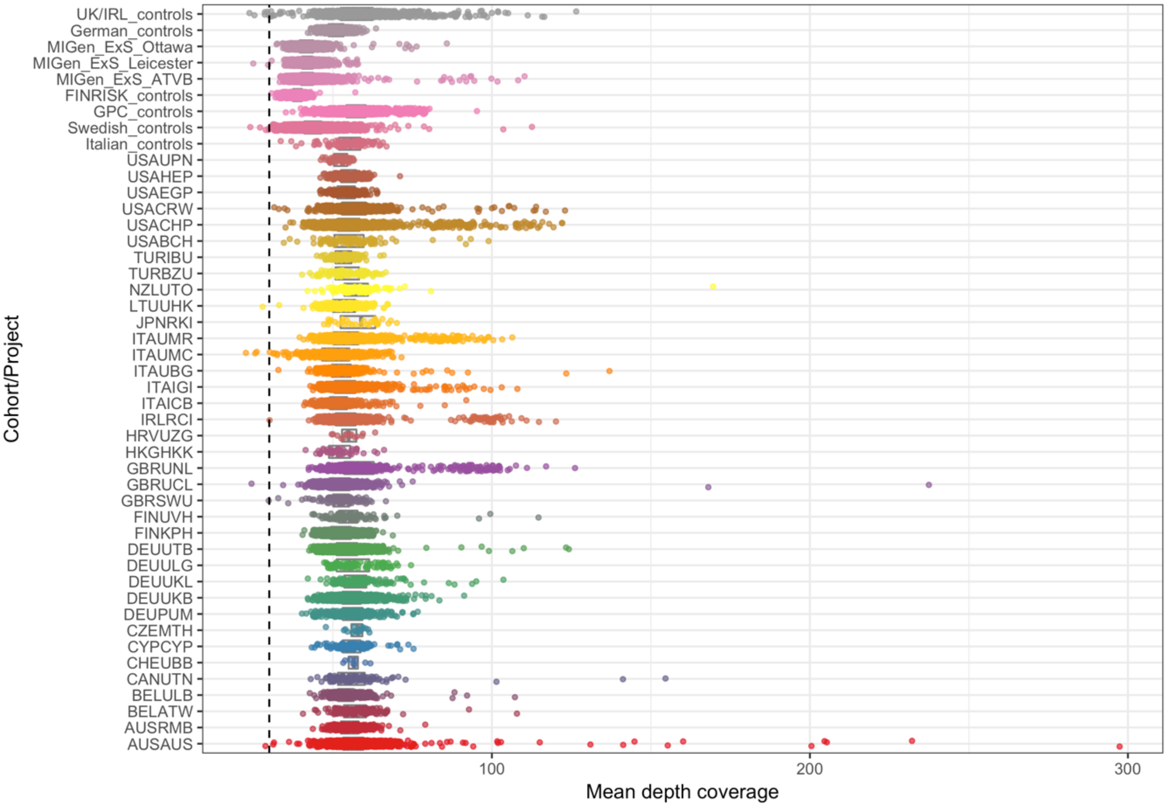


(B)

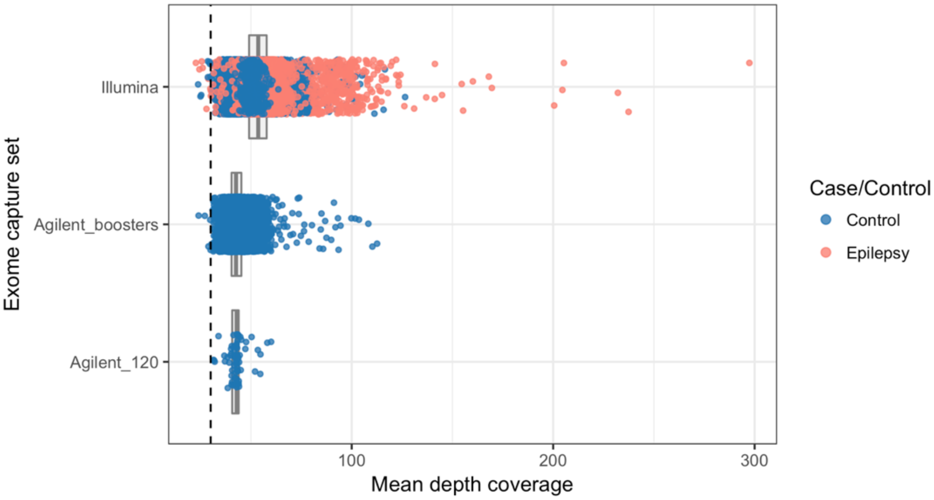


**Figure S2.** Average sample sequencing depth for pre-filtered data across cohorts (A) and by capture platform (B)

Cohorts with a six-letter name were case cohorts, and the rest were controls cohorts. Epilepsy cases overall had better average coverage than control samples due to capture platform difference (Illumina vs. Agilent exome enrichment kit). Dashed lines indicated the QC threshold, below which samples were removed (mean depth < 30).

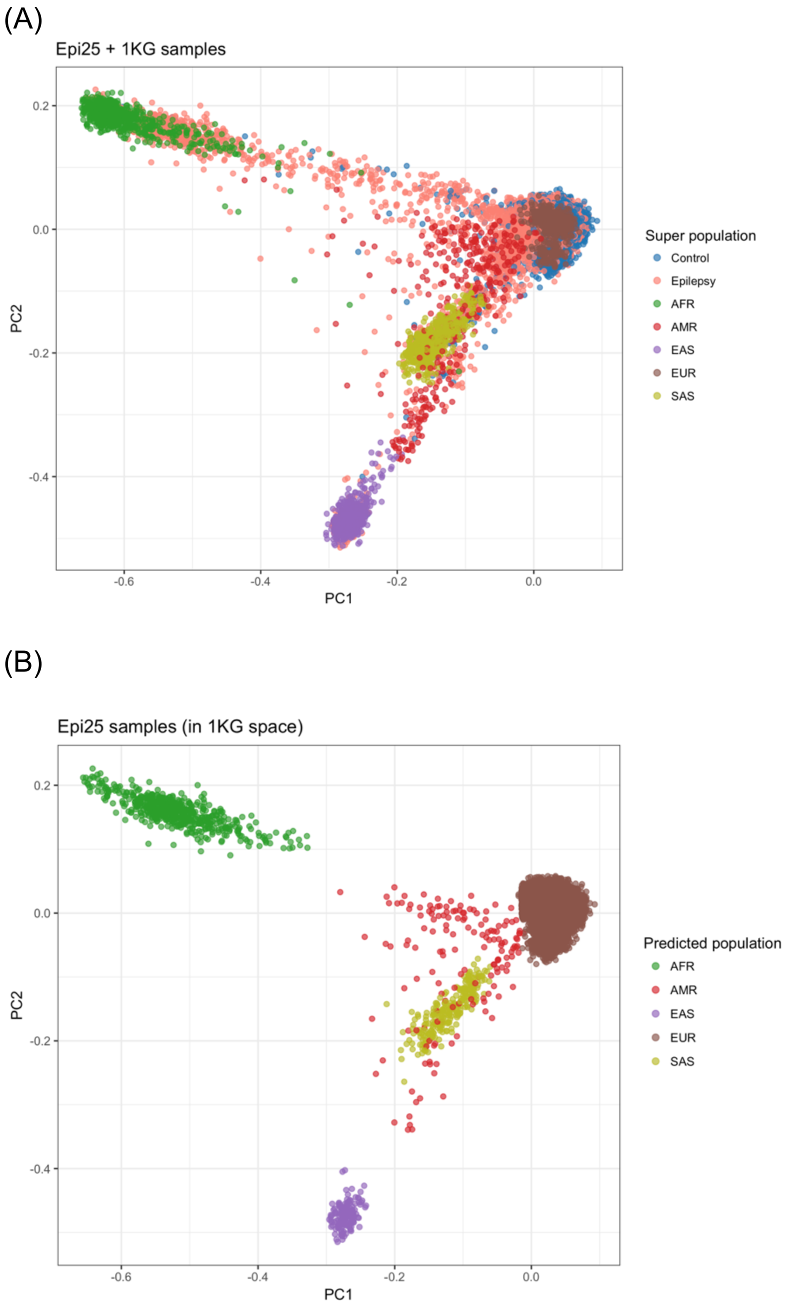


**Figure S3.** Population structure of the initial cases and controls based on principal component analysis (PCA)

(A) PCA was run on the study samples along with 1000 Genomes (1KG) phase 3 super populations to infer genetic ancestry. Epilepsy cases consisted of individuals from each of the super populations, while control samples were mostly of European descent.

(B) Genetic ancesty of the epi25 cases and controls were predicted by the Random Forest classifier based on top 6 PCs, using 1KG samples as the training data. Individuals were assgined to a particular 1KG-ancestry with a predicted probability >0.9, as depicted in the figure. We removed samples with unknown or unclassified ancestry and focused on European samples—the largest population under study with abundant cases and controls—moving forward.

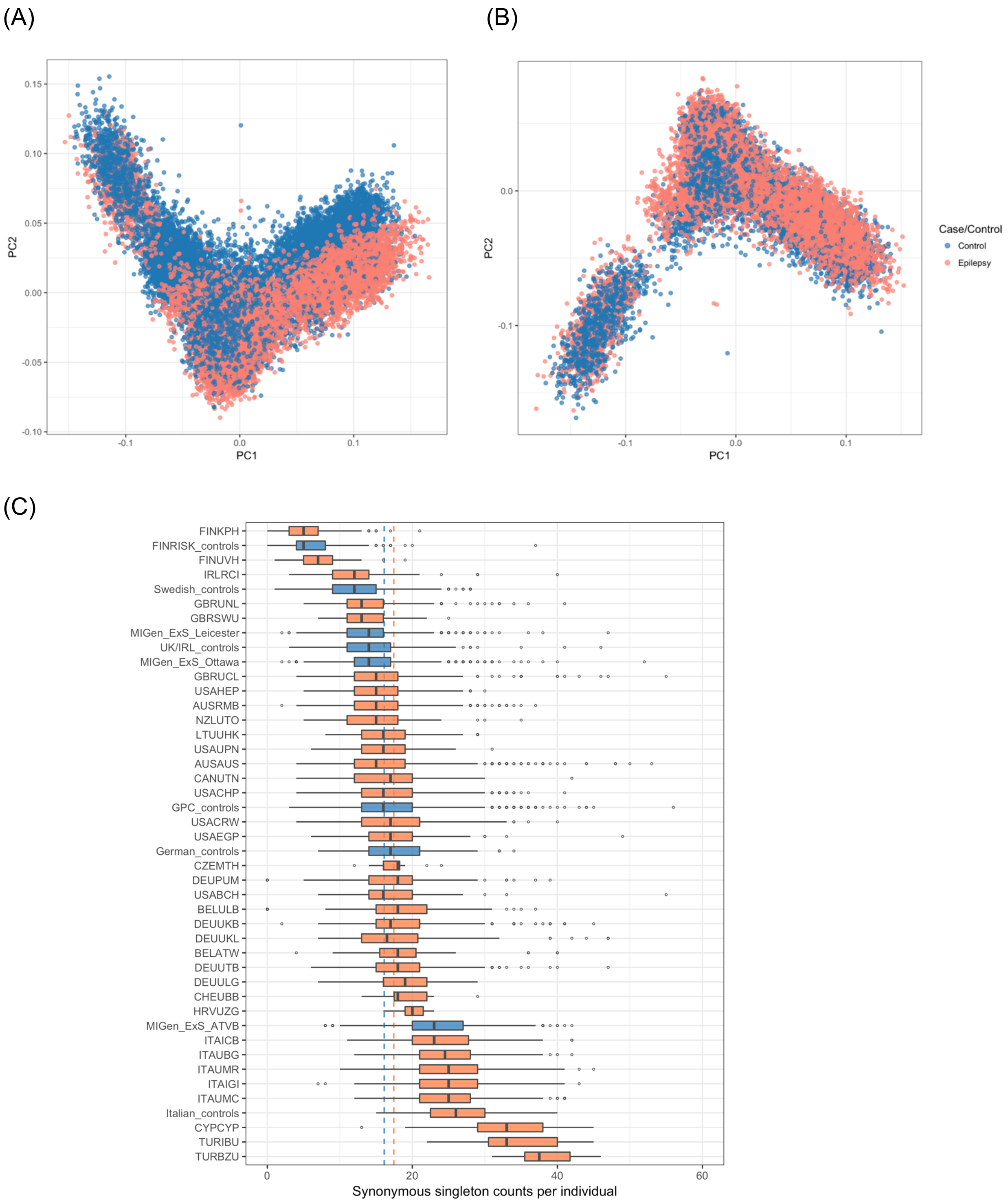

(A)
(B)
(C)
PC1
PC2
Case/Control
Control
Epilepsy
FINKPH
FINRISK_controls
FINUVH
IRLRCI
Swedish_controls
GBRUNL
GBRSWU
MIGen_ExS_Leicester
UK/IRL_controls
MIGen_ExS_Ottawa
GBRUCL
USAHEP
AUSRMB
NZLUTO
LTUUHK
USAUPN
AUSAUS
CANUTN
USACHP
GPC_controls
USACRW
USAEGP
German_controls
CZEMTH
DEUPUM
USABCH
BELULB
DEUUKB
DEUUKL
BELATW
DEUUTB
DEUULG
CHEUBB
HRVUZG
MIGen_ExS_ATVB
ITAICB
ITAUBG
ITAUMR
ITAIGI
ITAUMC
Italian_controls
CYPCYP
TURIBU
TURBZU
Synonymous singleton counts per individual

(D)

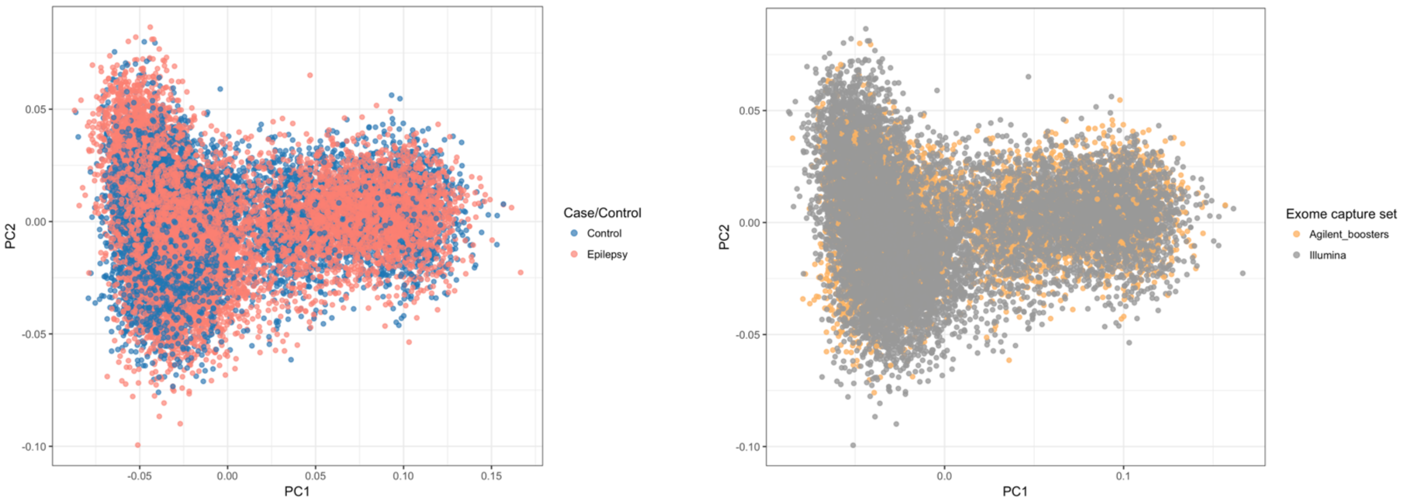


**Figure S4.** PCA on European samples

(A) Initial PCA on individuals classified as having a European descent. Ancestry was not well-matched between cases and controls.
(B) PCA after removing controls not pair-matched with cases based on top 2 PCs. Discarded controls were mostly Swedish and Finnish due to their unique genetic architecture.
(C) Synonymous singleton count distribution of ancestry-matched samples. This number tracks with intra-continental ancestry, showing a larger count in Southern Europeans relative to Northern Europeans. Case cohorts (orange) and control cohorts (blue) overall had a similar rate of singleton synonymous variants, indicated by the group averages (dashed lines). A few cohorts (Finland, Cyprus, Turkey) showed deviation from the mean and were removed from further analysis.
(D) Post-QC PC plots (PC1 vs. PC2), by case-control status and exome capture platform, showing a genetically homogeneous population for which analyses would be performed

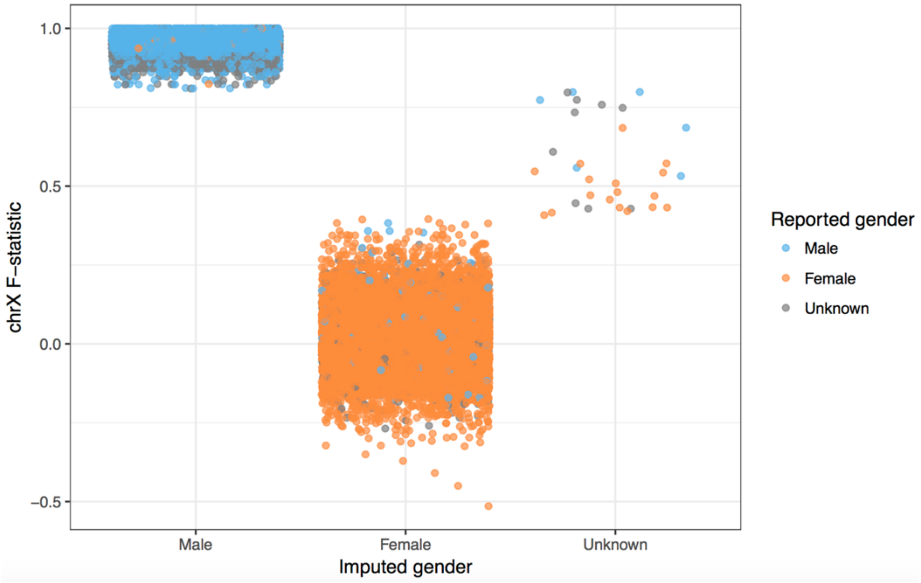


**Figure S5.** Sex check: concordance between genetically-imputed sex and self-reported sex

Sex was determined using the X-chromosome homozygosity rate, or F-statistics (≤0.4: female; ≥0.8: male; in between: unknown). Samples with either undefined genetic sex or discordant between imputed and reported gender were discarded.

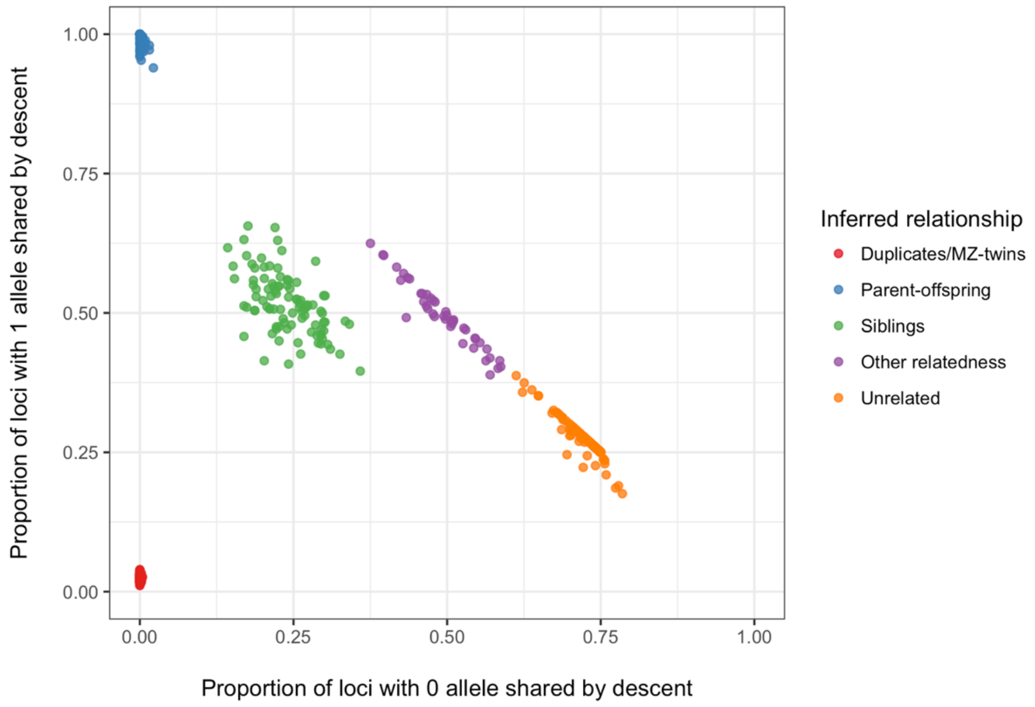


**Figure S6.** Identify-by-descent (IBD) estimate for ancestry-matched (EUR) samples

This plot shows only pairs of individuals with a relatedness > 0.125 for clarity. One individual in each of the related pairs with IBD > 0.2 was removed. Samples appearing in multiple related pairs were removed first, and cases were preserved over controls.

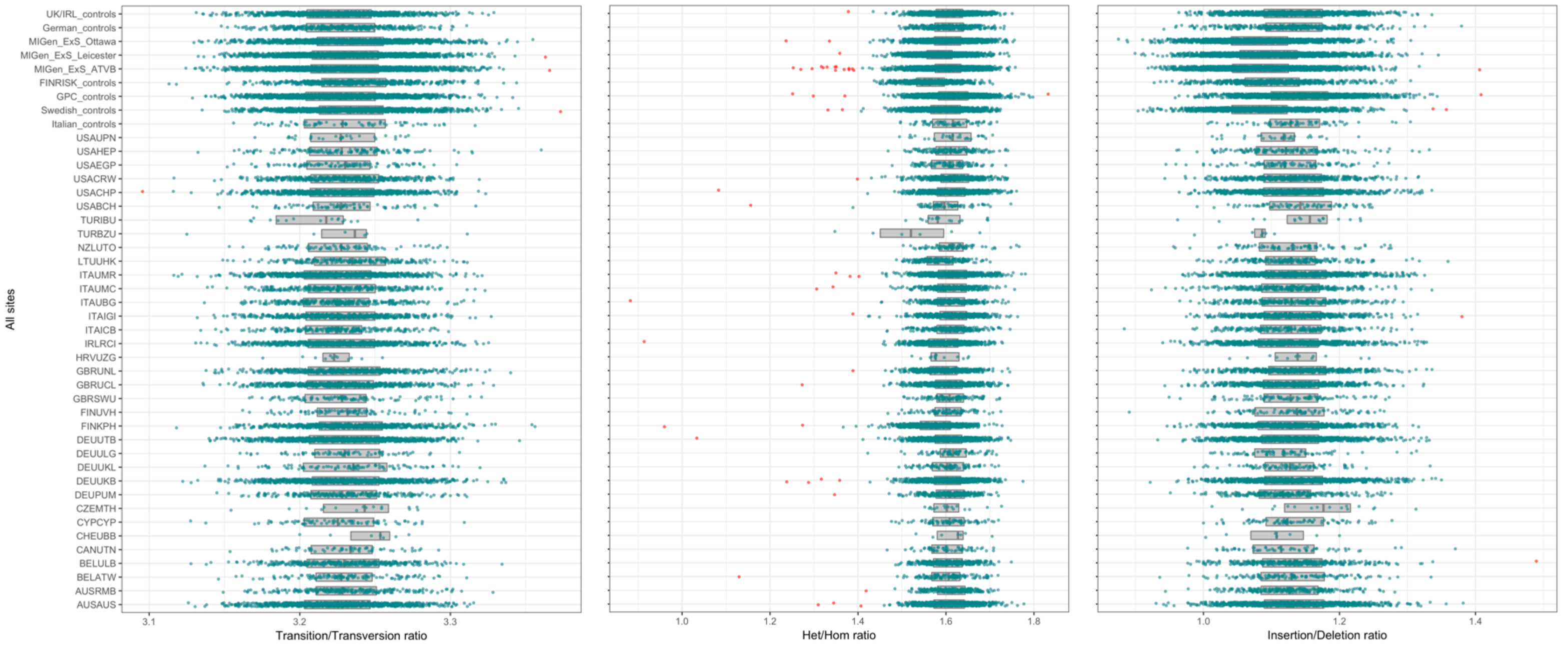


**Figure S7.** Sample outliers of QC metrics: transition to transversion ratio (Ti/Tv), heterozygous to homozygous ratio (Het/Hom), and insertion to deletion ratio.

Samples in each cohort that had a value outside the range of mean±4SD were removed (red dots).

(A)

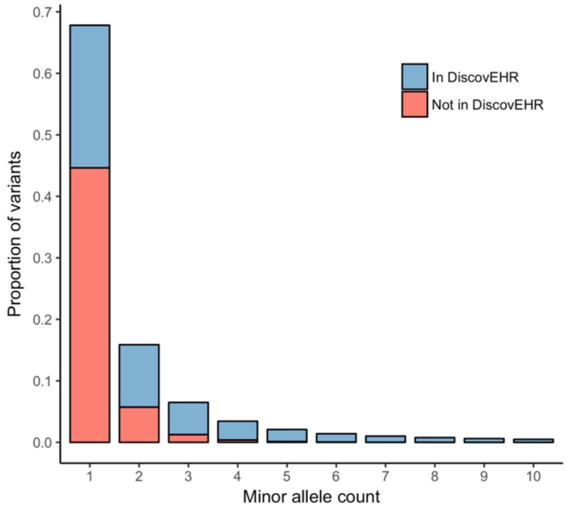


(B)

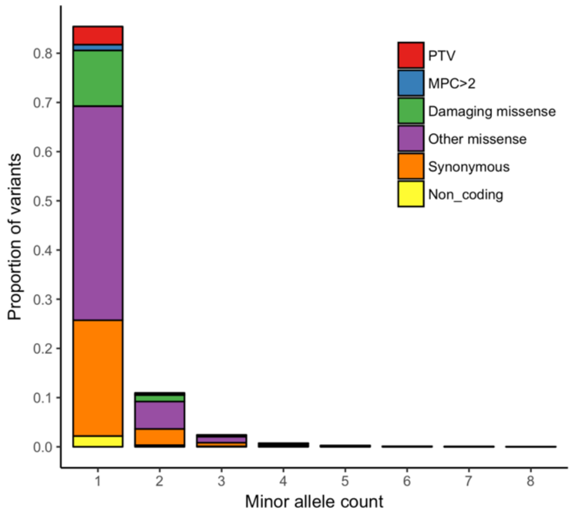


**Figure S8.** Post-QC allele count distribution

(A) showed the proportion of variants absent in DiscovEHR (48%) for which a majority were singletons
(B) showed the proportion of *non-DiscovEHR* sites by functional annotation. For the purpose of visualization, plotted here were non-overlapping proportions from the six functional classes of variants. Specifically, 4.1% were PTVs, 1.3% were missense variants with MPC≥2, 13% were non-MPC damaging missense, 50.1% were non-MPC benign missense variants (predicted by Polyphen-2 and SIFT), 27.9% were synonymous variants, and 2.8% were non-coding variants. In the analyses, however, the full damaging and benign missense annotations (predicted by PolyPhen-2 and SIFT) contained 59% and 41% of the MPC≥2 variants, respectively.

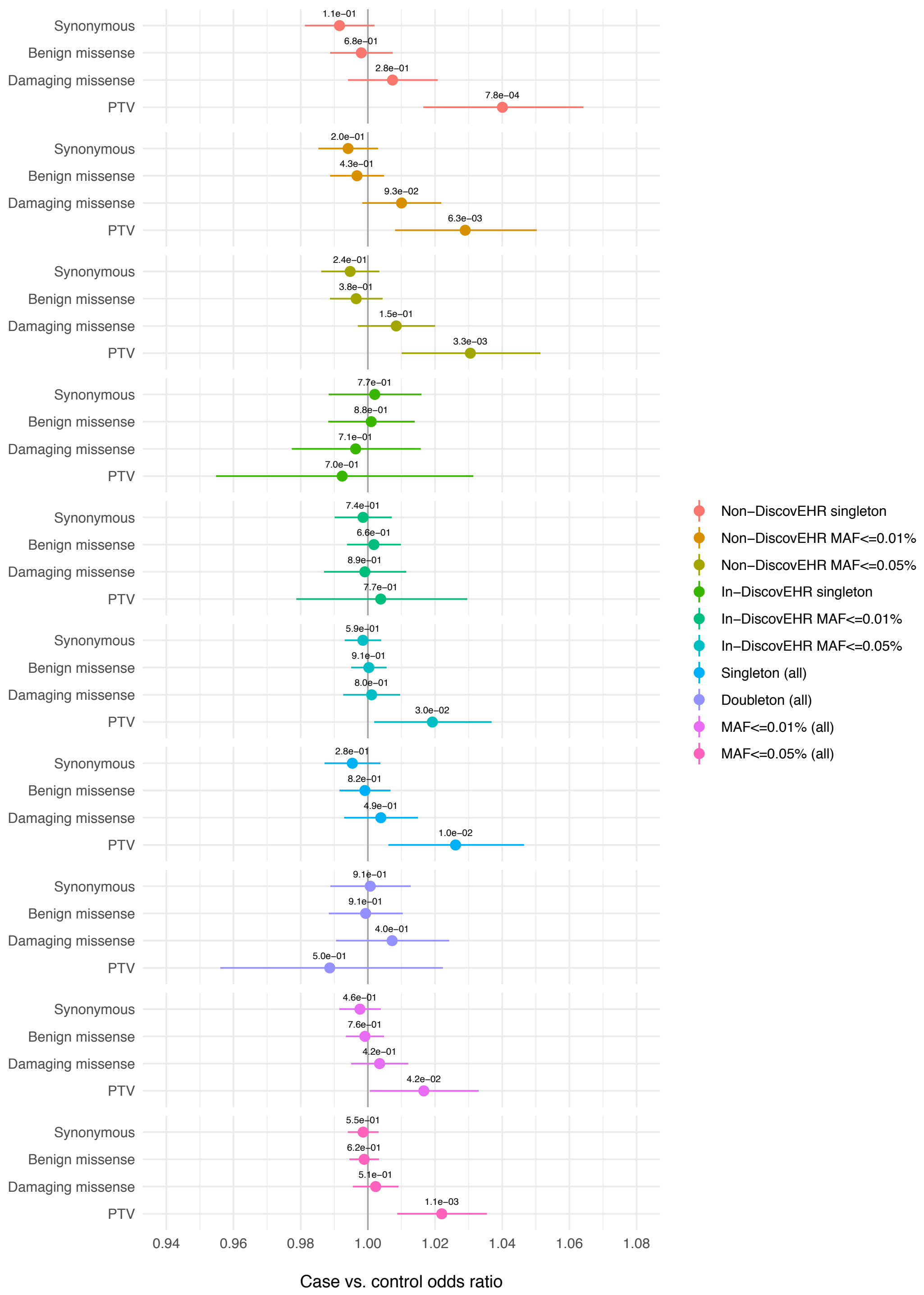


**Figure S9.** Exome-wide burden of rare variants in epilepsy versus controls across allele frequency categories

The burden of deleterious variants increased with a decrease in allele frequency. Risk of epilepsy was predominantly enriched among the singleton variants absent from the DiscovEHR cohort (a population allele frequency database).

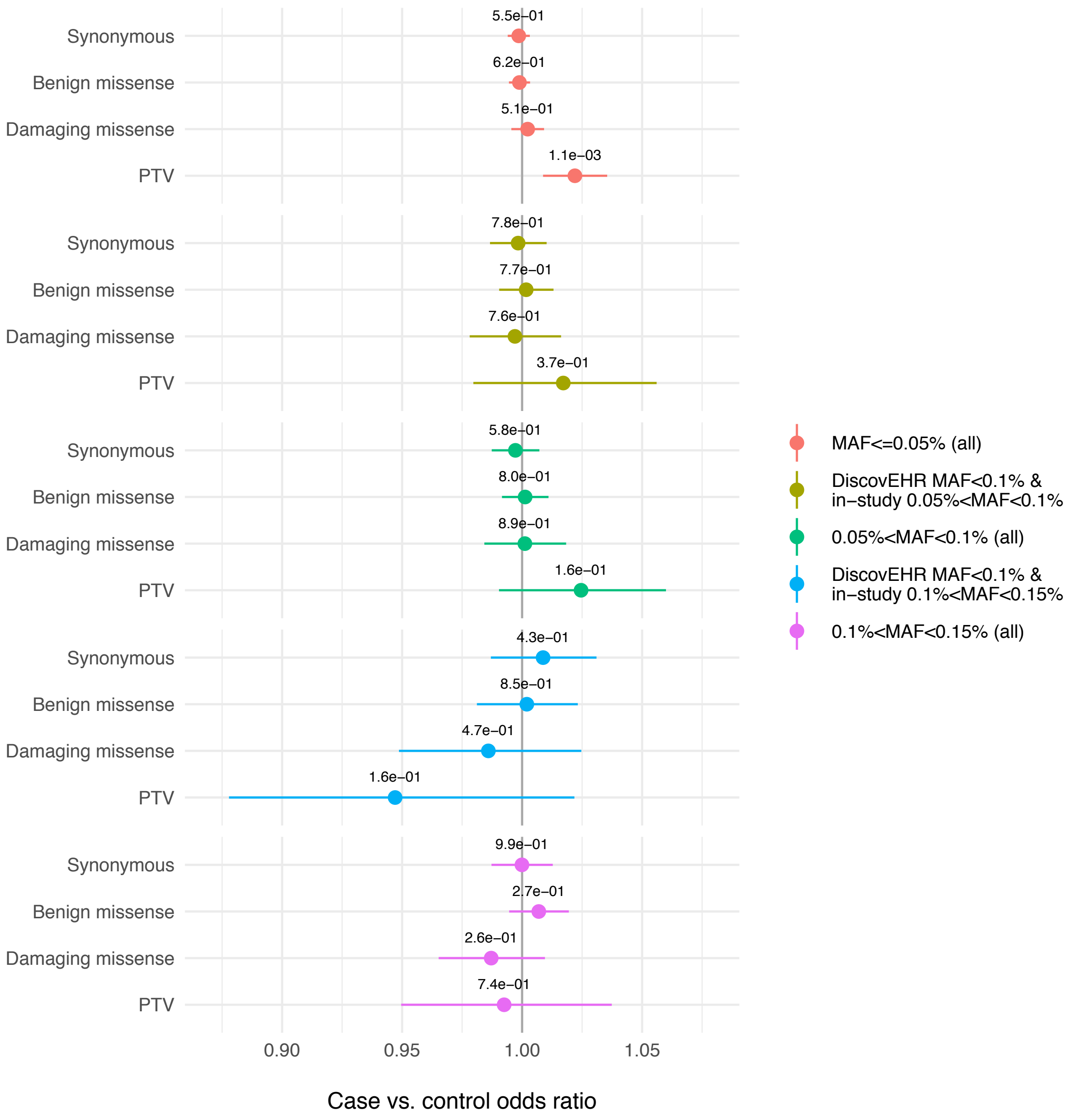


**Figure S10.** Exome-wide burden of rare and low-frequency variants in epilepsy versus controls

The excess burden of deleterious variants diminished in epilepsy with an increased in allele frequency, and are contained within ultra-rare variants (**Figure S9**).

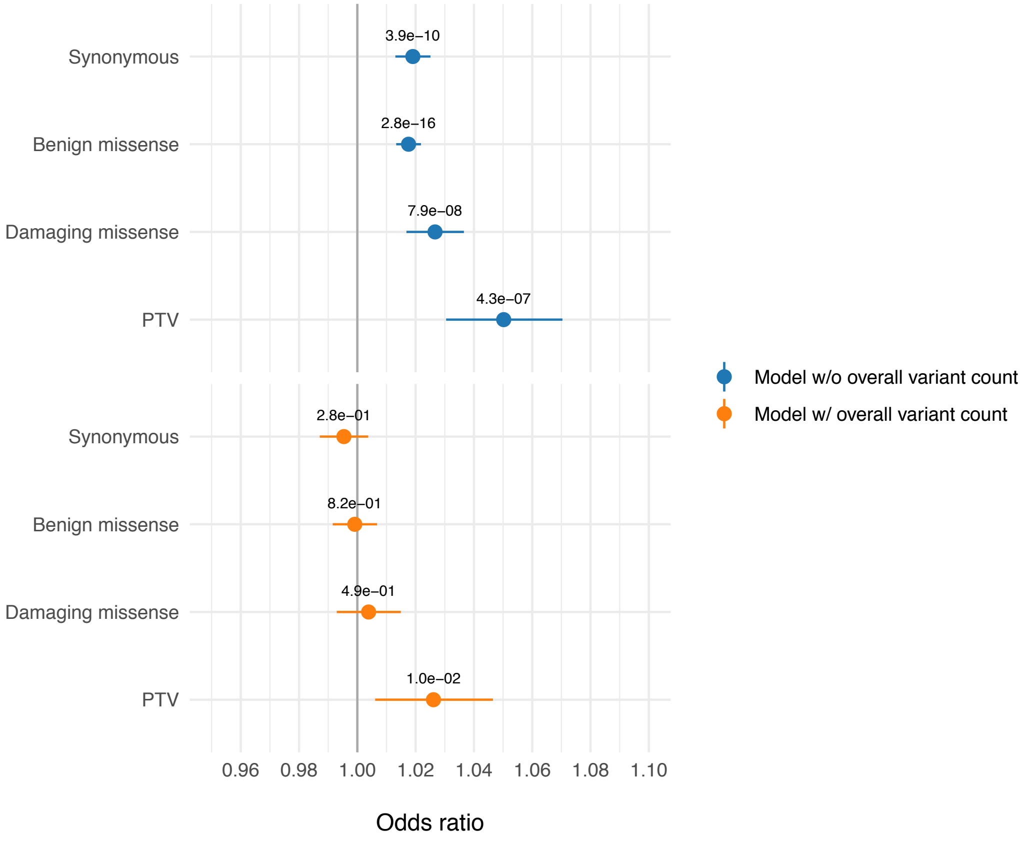


**Figure S11.** Correcting for exome-wide burden in gene-set burden test: controlling for overall variant count calibrated burden test p-values

This example was a burden analysis that assessed the exome-wide singleton enrichment in all epilepsy cases versus controls. Logistic regression model adjusting for total singleton count regardless of annotation (orange) successfully reduced the inflated burden across all annotation categories due to residual population stratification not captured by PCA.

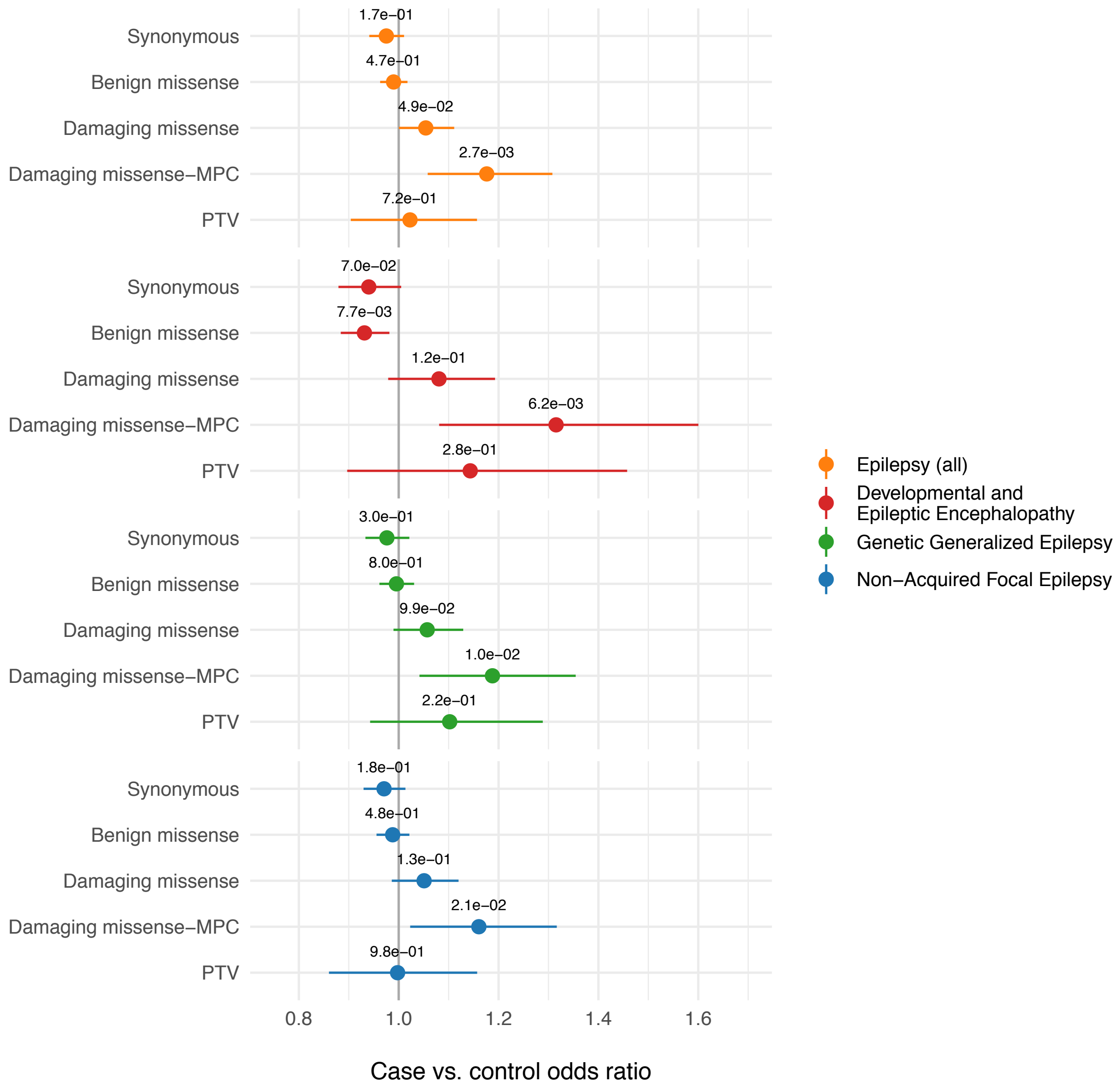


**Figure S12.** Burden of ultra-rare singletons in LoF-intolerant genes with 0.9 < pLI < 0.995), using only non-ExAC controls (N=4,042)

Burden signals of PTVs disappeared after restricting to genes within the range of 0.9 < pLI < 0.995, suggesting that most enrichment of PTVs in haploinsufficient genes associated with epilepsy was driven by genes with a pLI > 0.995.

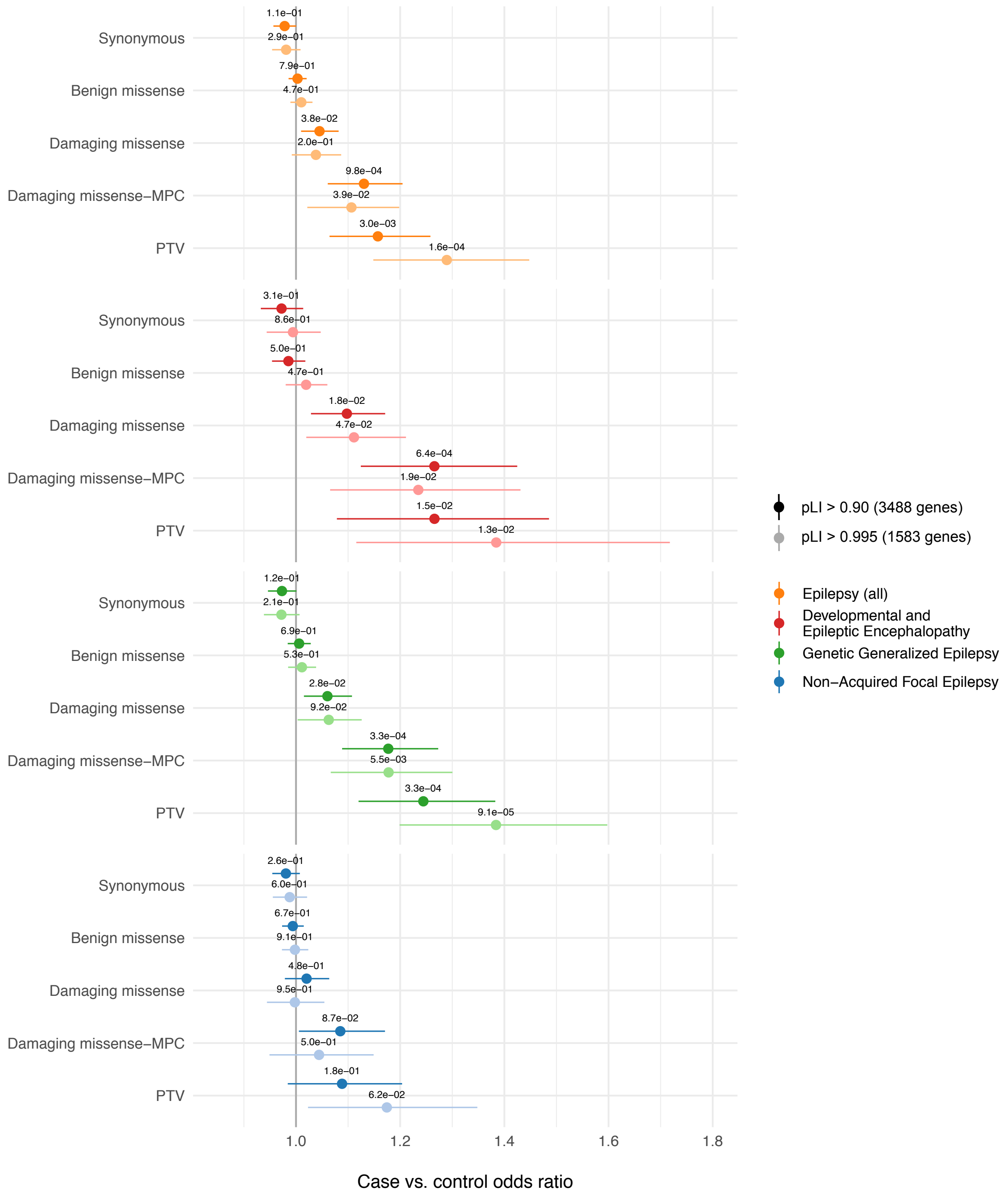


**Figure S13.** Burden of ultra-rare singletons in LoF-intolerant genes (pLI > 0.9 and > 0.995), using only non-ExAC controls (N=4,042)

Significance of association was displayed in FDR-adjusted p-values. A strongly significant enrichment of PTVs and damaging missense variants (MPC≥2) were seen in patients with DEE and GGE, but not NAFE (FDR < 0.05). PTV burden was even stronger when restricting to genes with pLI>0.995. Only non-ExAC controls were used as pLI scores were derived from ExAC samples.

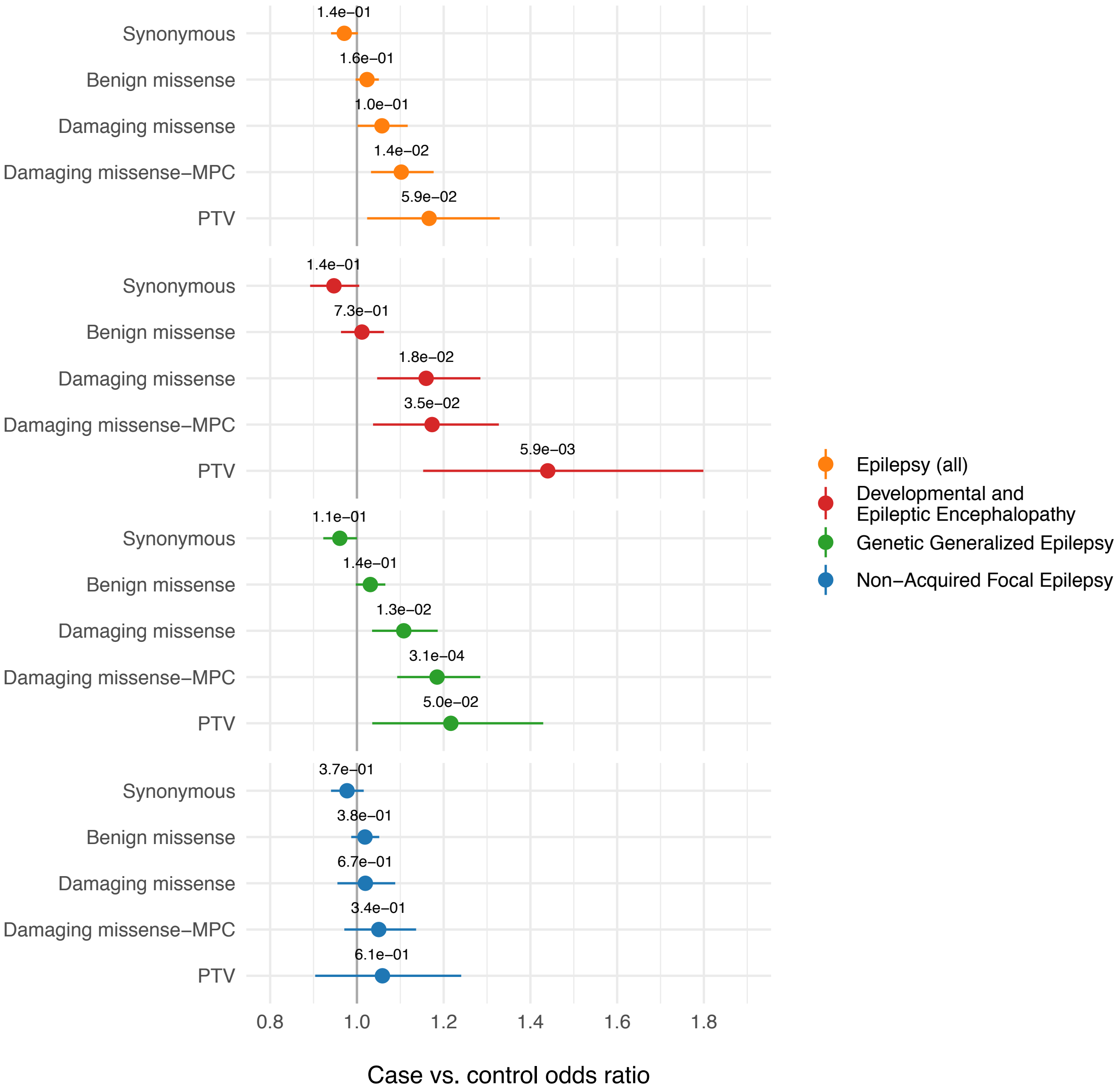


**Figure S14.** Burden of ultra-rare singletons in missense constrained genes (mi-Z > 3.09), using only non-ExAC controls (N=4,042)

Significance of association was displayed in FDR-adjusted p-values. PTV and damaging missense (MPC≥2) burden were significant for all epilepsy phenotypes at FDR < 0.05 except NAFE.

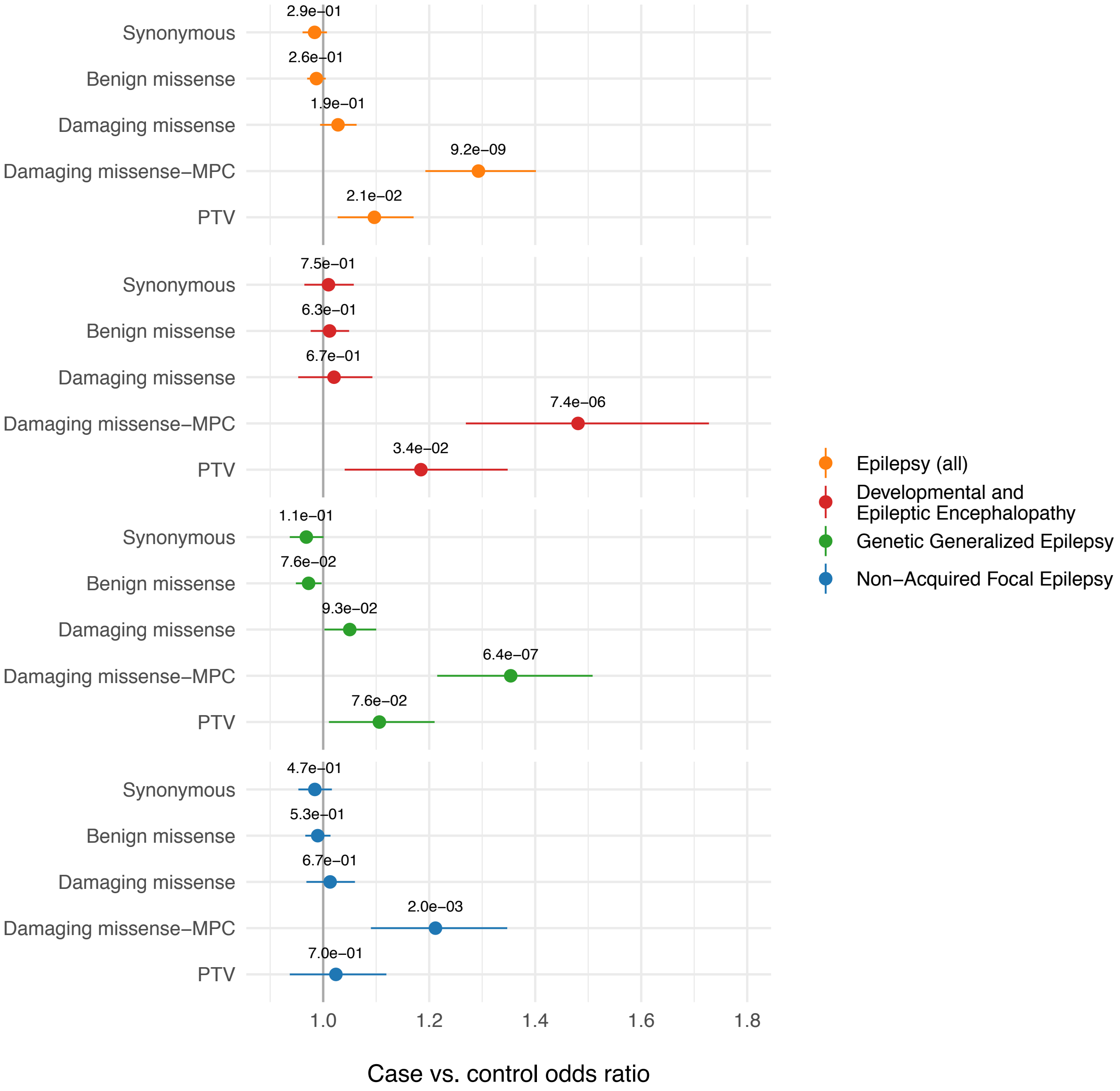


**Figure S15.** Burden of ultra-rare singletons in 2,649 brain-enriched genes

Significance of association was displayed in FDR-adjusted *p*-values. Damaging missense (MPC≥2) variants in brain-enriched genes were significantly enriched in all epilepsy types and had a much higher burden relative to PTVs. PTV burden was stronger in patients with DEE (FDR < 0.05) than with common epilepsy types.

(A)

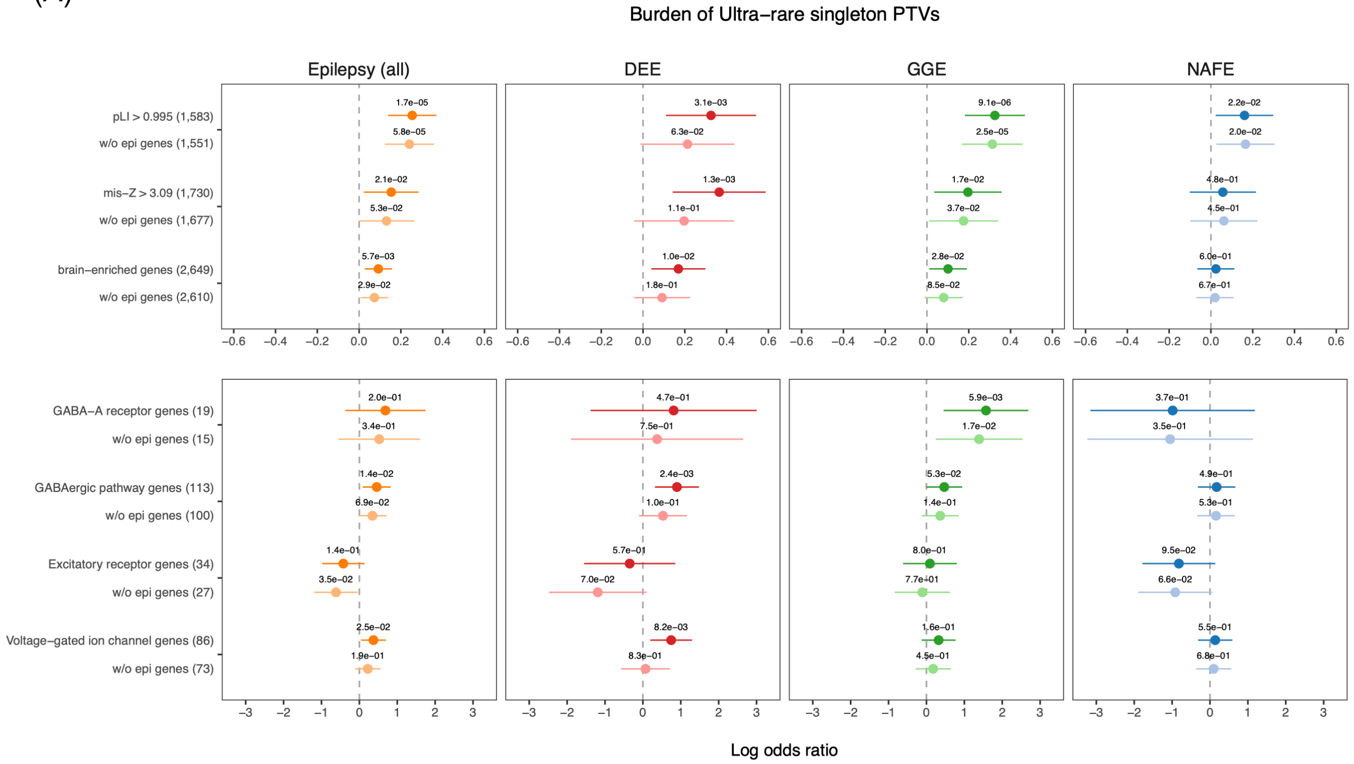


(B)

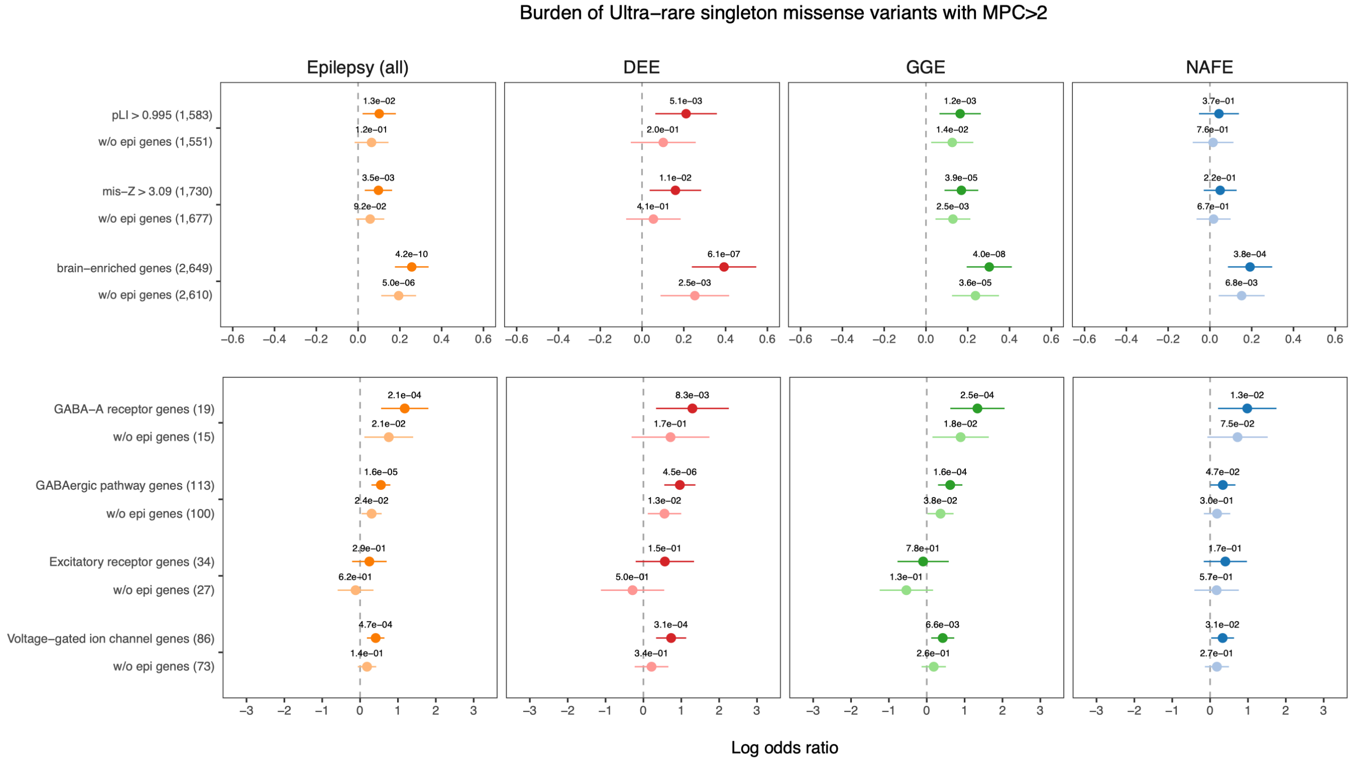


**Figure S16.** Burden of ultra-rare singleton PTVs (A) and missese-MPC≥2 variants (B) before and after correcting for events in genes previously associated with epilepsy

Genes previously associated with epilepsy included 74 non-overlapping genes from the 43 dominant epilepsy genes, 50 dominant DEE genes, and 33 NDD-EPI genes (**Table S6**). For each gene set, the darker (upper) color showed burden signals before adjusting for counts in the 74 genes and the lighter (lower) color showed the residual signals after correcting for associations driven by the 74 genes. P-values were shown in original values.

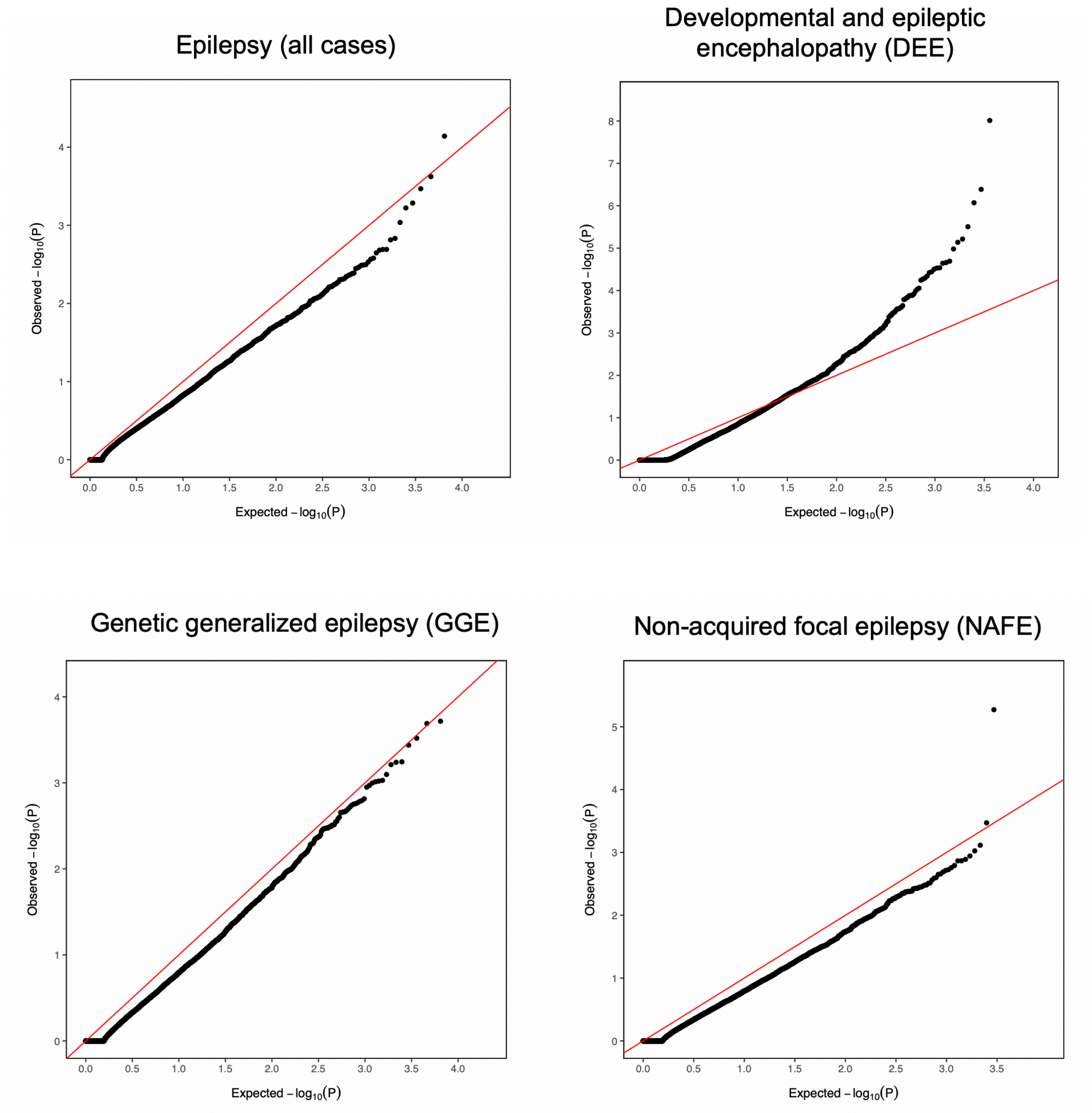


**Figure S17.** Burden of all low-frequency damaging missense variants assessed using SKAT

Shown were the SKAT gene burden results based on damaging missense variants (predicted by PolyPhen-2 and SIFT) with MAF < 0.01. Exome-wide significant gene associations were observed only among DEE patients compared to controls, where the top genes included previously associated genes (e.g. *STXBP1*, P=9.3e-09; *KCNA2*, P=1.0e-05). Burden of missense variants with MPC≥2 showed a very similar pattern but was in general more underpowered due to low counts (data not shown).

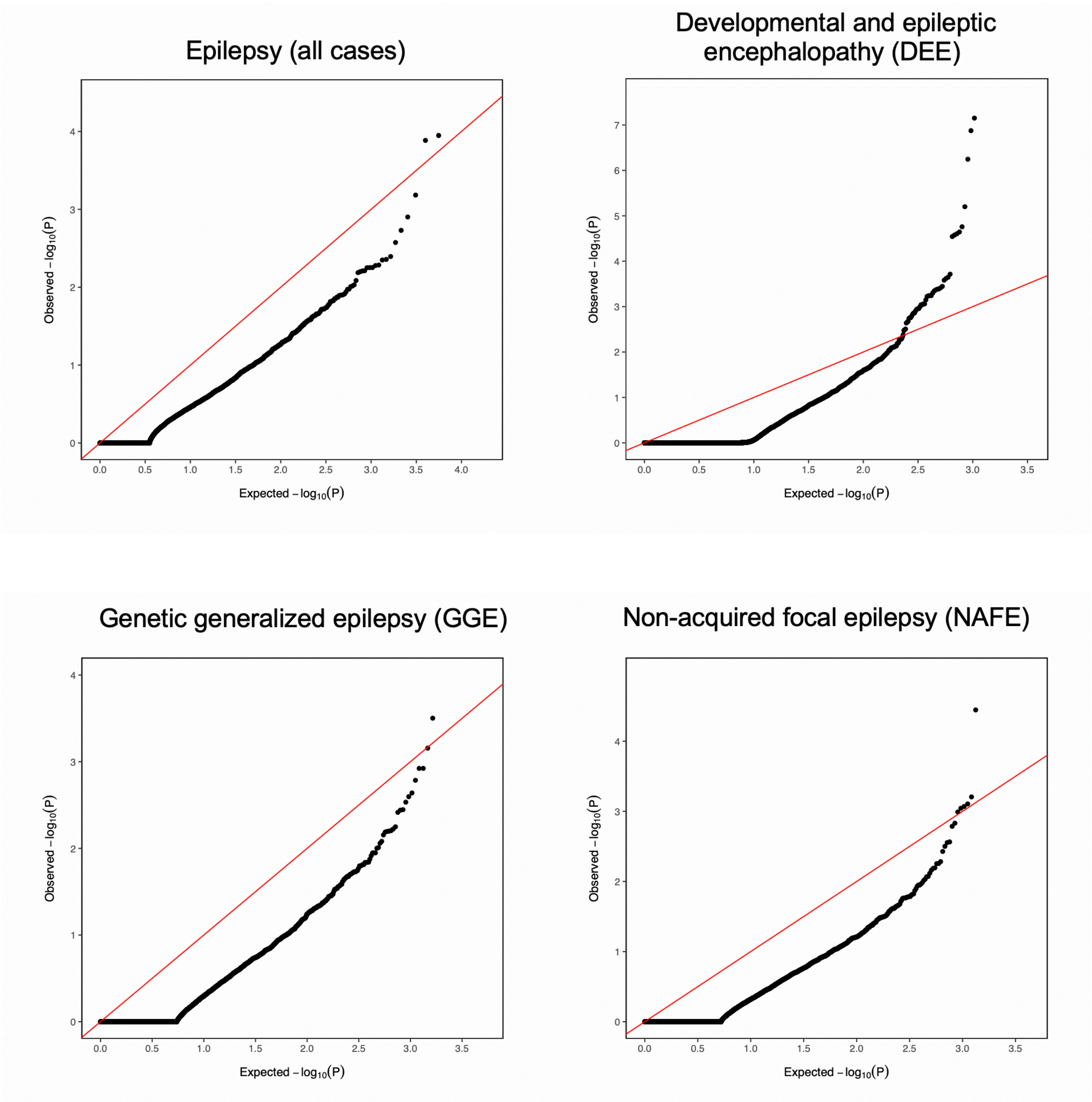


**Figure S18.** Burden of all low-frequency protein truncating variants assessed using SKAT

Shown were the SKAT gene burden results based on protein truncating variants with MAF < 0.01. Exome-wide significant gene associations were observed only among DEE patients compared to controls, with the top genes including some previously associated genes (e.g. *KIAA2022, P* = 7.1e-08; *SCN1A*, *P* = 3.9e-04). Associations with other epilepsy phenotypes were largely underpowered due to low counts.

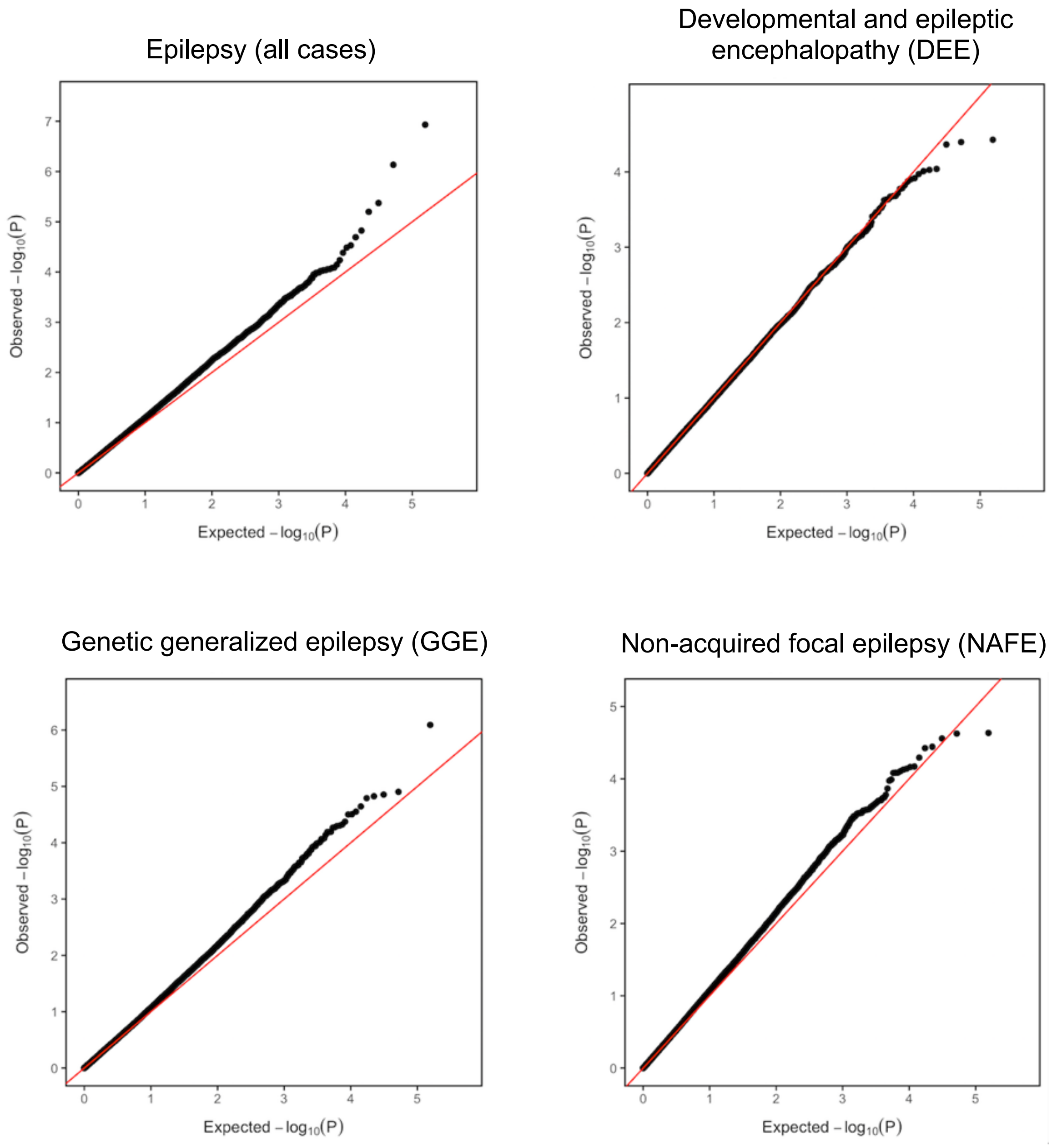


**Figure S19.** Association with common and low frequency variants (for variants with MAF > 0.001)

After multiple-testing correction, no coding variants were significantly associated with overall epilepsy (all cases) or within any epilepsy type.

(A) ***TRIM3*** (2 DEE, 2 GGE, 7 NAFE, 3 Lesional)

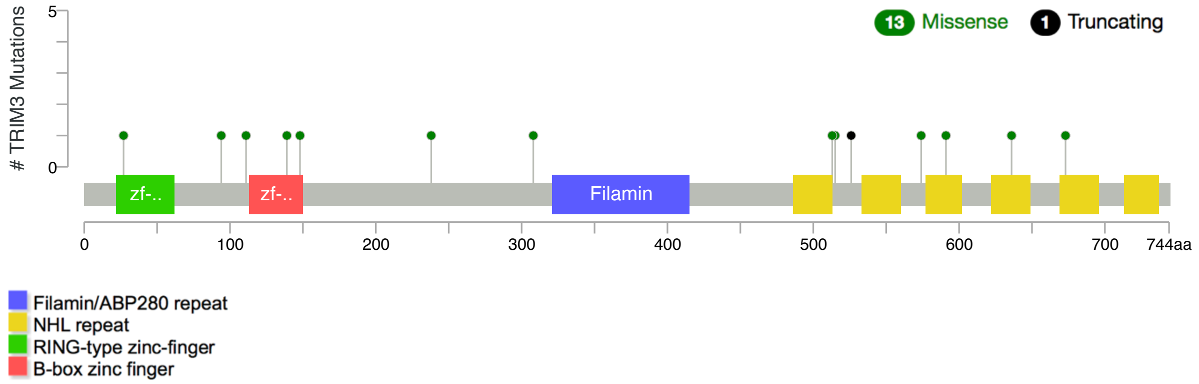


(B) ***GABRG2*** (7 GGE, 7 NAFE, 1 Lesional, 1 Control)

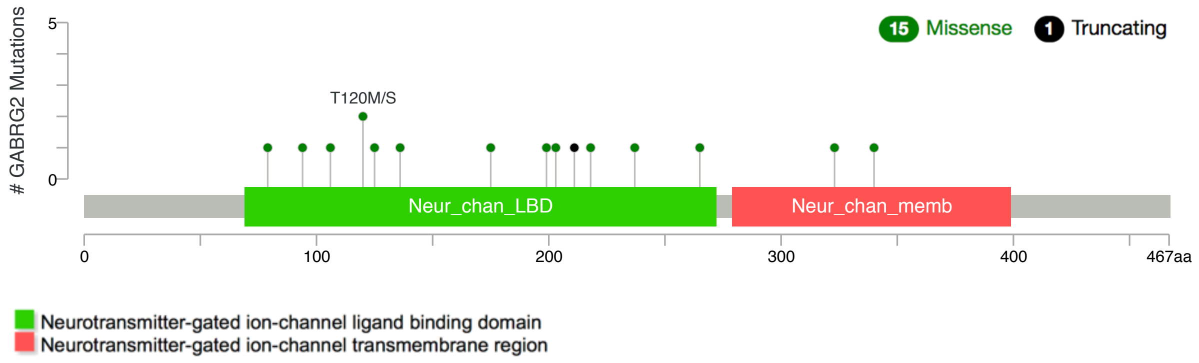


(C) ***CACNA1G*** (2 DEE, 10 GGE, 4 NAFE, 3 Lesional, 3 Control)

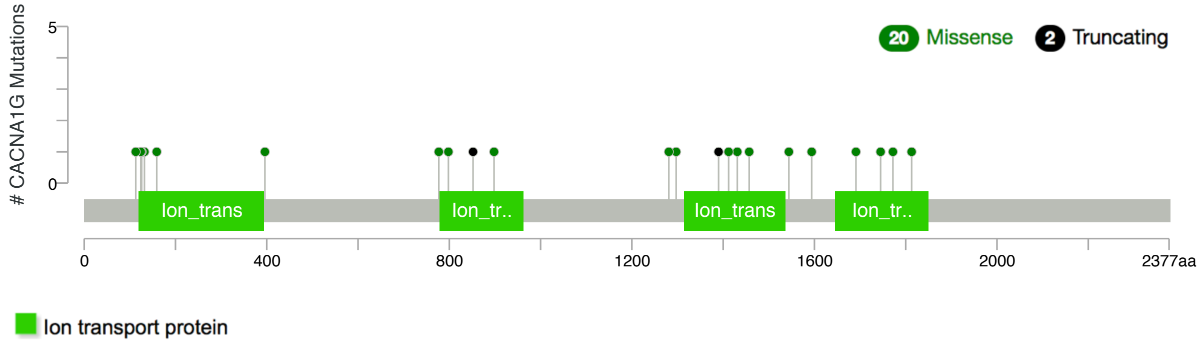


**Figure S20.** Mutational diagrams of selected top genes in all-epilepsy gene burden analysis

These genes were shown to harbor a higher burden of missense (MPC≥2) than protein-truncating URVs in epilepsy patients.

**Table S1.** Epi25 epilepsy patients for whole-exome sequencing (WES)

DEE: developmental and epileptic encephalopathy; GGE: genetic generalized epilepsy; NAFE: non-acquired focal epilepsy; LFE: lesional focal epilepsy; FS: febrile seizure; UE: unclassified epilepsy (including those under review or labeled excluded)

| Site Name | Site Code | Epilepsy | | | | | | TOTAL |
|---|---|---|---|---|---|---|---|---|
| | | DEE | GGE | NAFE | LFE | FS | UE | |
| **Australia: Melbourne** | **AUSAUS** | 139 | 412 | 366 | 236 | 57 | 6 | 1216 |
| **Australia: Royal Melbourne** | **AUSRMB** | 0 | 88 | 128 | 42 | 0 | 12 | 270 |
| **Belgium: Antwerp** | **BELATW** | 87 | 36 | 20 | 0 | 0 | 0 | 143 |
| **Belgium: Brussels** | **BELULB** | 6 | 71 | 190 | 120 | 0 | 0 | 387 |
| **Canada: Andrade** | **CANUTN** | 37 | 43 | 8 | 3 | 0 | 5 | 96 |
| **Switzerland: Bern** | **CHEUBB** | 0 | 0 | 6 | 2 | 0 | 0 | 8 |
| **Cyprus** | **CYPCYP** | 7 | 53 | 52 | 4 | 0 | 2 | 118 |
| **Czech Republic: Prague** | **CZEMTH** | 16 | 0 | 0 | 0 | 0 | 0 | 16 |
| **Germany: Frankfurt/Marburg** | **DEUPUM** | 4 | 58 | 130 | 60 | 0 | 7 | 259 |
| **Germany: Bonn** | **DEUUKB** | 0 | 206 | 557 | 61 | 193 | 36 | 1053 |
| **Germany: Kiel** | **DEUUKL** | 47 | 81 | 20 | 0 | 0 | 1 | 149 |
| **Germany: Leipzig** | **DEUULG** | 0 | 0 | 89 | 0 | 0 | 0 | 89 |
| **Germany: Tuebingen** | **DEUUTB** | 63 | 346 | 256 | 351 | 33 | 6 | 1055 |
| **Finland: Kuopio** | **FINKPH** | 20 | 55 | 636 | 21 | 0 | 2 | 734 |
| **Finland: Helsinki** | **FINUVH** | 26 | 51 | 17 | 1 | 0 | 0 | 95 |
| **Wales: Swansea** | **GBRSWU** | 0 | 45 | 56 | 0 | 0 | 6 | 107 |
| **UK: UCL** | **GBRUCL** | 5 | 304 | 314 | 2 | 0 | 87 | 712 |
| **UK: Imperial/Liverpool** | **GBRUNL** | 0 | 190 | 343 | 0 | 0 | 0 | 533 |
| **Hong Kong** | **HKGHKK** | 0 | 21 | 23 | 0 | 0 | 0 | 44 |
| **Croatia** | **HRVUZG** | 11 | 2 | 0 | 0 | 0 | 1 | 14 |
| **Ireland: Dublin** | **IRLRCI** | 14 | 145 | 464 | 155 | 15 | 18 | 811 |
| **Italy: Milan** | **ITAICB** | 50 | 97 | 14 | 29 | 0 | 6 | 196 |
| **Italy: Genova** | **ITAIGI** | 100 | 269 | 13 | 26 | 0 | 2 | 410 |
| **Italy: Bologna** | **ITAUBG** | 82 | 60 | 117 | 34 | 0 | 6 | 299 |
| **Italy: Catanzaro** | **ITAUMC** | 5 | 72 | 264 | 28 | 0 | 4 | 373 |
| **Italy: Florence** | **ITAUMR** | 402 | 186 | 128 | 155 | 0 | 56 | 927 |
| **Japan: RIKEN Institute** | **JPNRKI** | 0 | 31 | 0 | 0 | 0 | 0 | 31 |
| **Lithuania** | **LTUUHK** | 33 | 96 | 70 | 14 | 0 | 3 | 216 |
| **New Zealand: Otago** | **NZLUTO** | 27 | 42 | 42 | 6 | 0 | 0 | 117 |
| **Turkey: Bogazici** | **TURBZU** | 89 | 0 | 0 | 0 | 0 | 0 | 89 |
| **Turkey: Istanbul** | **TURIBU** | 5 | 50 | 52 | 6 | 0 | 1 | 114 |
| **USA: BCH** | **USABCH** | 60 | 12 | 9 | 7 | 1 | 5 | 94 |
| **USA: Philadelphia/CHOP** | **USACHP** | 0 | 988 | 479 | 0 | 0 | 0 | 1467 |
| **USA: Philadelphia/Rowan** | **USACRW** | 0 | 324 | 236 | 0 | 0 | 0 | 560 |
| **USA: EPGP** | **USAEGP** | 126 | 2 | 1 | 1 | 0 | 0 | 130 |
| **USA: NYU HEP** | **USAHEP** | 0 | 0 | 214 | 0 | 0 | 0 | 214 |
| **USA: Penn/CHOP** | **USAUPN** | 15 | 17 | 17 | 1 | 0 | 0 | 50 |
| **TOTAL** | | **1476** | **4453** | **5331** | **1365** | **299** | **272** | **13196** |

**Table S2.** Control collection for Epi25 WES analysis

| Control cohorts | General control | MI/CAD/CHD[1] | Exome capture platform | Source | dbGaP Accession or PMID | In ExAC |
|---|---|---|---|---|---|---|
| MIGen ATVB | 1802 | 1875 | Agilent | dbGaP | phs000814.v1.p1 | Yes |
| MIGen Leicester | 1100 | 0 | Illumina | dbGaP | phs001000.v1.p1 | No |
| MIGen Ottawa Heart Study | 987 | 993 | Agilent | dbGaP | phs000806.v1.p1 | Yes |
| Genomic Psychiatry Cohort (GPC) controls | 1946 | 0 | Illumina | Broad local | 23650244[23] | No |
| German controls | 414 | 0 | Illumina | Broad local | N/A | No |
| FINRISK controls | 681 | 0 | Illumina | Broad local | 29165699[22] | No |
| Swedish SCZ controls | 6242 | 0 | Agilent | dbGaP | phs000473.v2.p2 | Yes |
| UK/IRL controls | 1223 | 0 | Illumina | Broad local | N/A | No |
| Italian controls | 106 | 0 | Illumina | Broad local | N/A | No |
| Epi25 Italian controls (ITAICB & ITAIGI) | 300 | 0 | Illumina | Epi25 | N/A | No |
| **Total** | **17669** | | | | | |

[1]Myocardial infarction, coronary artery disease, or coronary heart disease

**Table S3.** Sample QC and number of sample dropout at each QC step

| Sample QC metric | Number of cases (%) | Number of controls (%) |
|---|---|---|
| **Initial numbers** | **13,196** (100%) | **17,669** (100%) |
| ***Initial sample QC***<br>Call rate < 0.98 / %Chimeras > 0.014<br>Avg. sequence depth < 30<br>Avg. genotype quality < 85<br>Freemix contamination > 0.04 | 84 (0.63%) | 372 (2.11%) |
| ***PCA*** | | |
| Non-European | 2034 (15.4%) | 338 (1.91%) |
| Case-Control ancestry matching (using top PCs) | 0 | 6838 (38.7%) |
| ***Relatedness filtering*** | | |
| IBD > 0.2 | 218 (1.65%) | 149 (0.84%) |
| ***Sex check*** | | |
| Ambiguous imputed sex | 6 (0.05%) | 5 (0.03%) |
| Imputed sex not equal to reported sex | 148 (1.12%) | 25 (0.14%) |
| ***Outlier removal*** | | |
| Per-cohort outliers (>4 SD) of Ti/Tv, Het/Hom, or indel ratio | 34 (0.19%) | 34 (0.19%) |
| ***Control for residual population stratification*** (using #synonymous singletons) | | |
| Swedish/Finnish: lower counts | 1232 (9.33%) | 1472 (8.33%) |
| Cypriot/Turkish: higher counts | 99 (0.75%) | 0 |
| TBD-phenotype cases / Excluded by review | 171 (1.30%) | 0 |
| **Final numbers** | **9,170** (69.5%) | **8,436** (47.7%) |

**Table S4.** Variant QC and number of variant dropout at each QC step

| Variant QC metric | Number of variants removed | %Removed |
|---|---|---|
| ***Pre-filtering:*** | | |
| Failing VQSR | 614,459 | 8.90% |
| In low complexity region | 69,003 | 1.00% |
| Outside well-covered regions | 2,996,409 | 43.40% |
| ***Initial QC:*** | | |
| Call rate < 0.975<br>Case- & Ctrl-call rate < 0.975 | 220,614 | 3.20% |
| Case/Ctrl call rate diff > 0.005 | 224,033 | 3.20% |
| pHWE < 1e-06 | 662 | 0.01% |
| AC = 0 | 732,640 | 10.60% |
| ***Final QC:***<br>Call rate < 0.98<br>Case/Ctrl call rate diff > 0.005<br>pHWE < 1e-06<br>AC = 0 | 195,586 | 2.80% |
| Associated with capture platform (p<0.05) | 5,906 | 0.09% |

**Table S5.** Grouping of functional consequences of the called sites

| Consequence | VEP Terms/annotation |
|---|---|
| Protein-truncating variant (PTV) | "transcript_ablation", "splice_acceptor_variant", "splice_donor_variant", "stop_gained", or "frameshift_variant" |
| Missense | "stop_lost", "start_lost", "transcript_amplification", "inframe_insertion", "inframe_deletion", "missense_variant", "protein_altering_variant", or "splice_region_variant" |
| Benign/Other missense | Not classified by PolyPhen-2 and SIFT as damaging or deleterious |
| Damaging missense | PolyPhen-2 "probably_damaging" & SIFT "deleterious" |
| Damaging missense-MPC | MPC score (**M**issense badness, **P**olyPhen-2, and **C**onstraint) ≥ 2 |
| Synonymous | "incomplete_terminal_codon_variant", "stop_retained_variant", "synonymous_variant" |

**Table S6.** Gene sets collected for burden analysis

| Gene set | Number of genes | Reference | | Genes (display only smaller gene sets) |
|---|---|---|---|---|
| **Constrained genes** | | | | |
| LoF-intolerant (pLI > 0.9) | 3,488 | Samocha 2014[24] | ExAC non-psych | |
| LoF-intolerant (pLI > 0.995) | 1,583 | Samocha 2014[24] | ExAC non-psych | |
| Missense constrained (misZ > 3.09) | 1,730 | Samocha 2014[24] | ExAC non-psych | |
| **Tissue-specific** | | | | |
| Brain enriched (GTEx) | 2,649 | Ganna 2016[25] | | |
| **Candidate epilepsy genes** | | | | |
| Known epilepsy genes | 43 | Epi4K 2017[26] | Table S5 | *SPTAN1, SCN8A, CHD2, SCN2A, GRIN2B, GRIN2A, SYNGAP1, KCNMA1, PRICKLE2, SCN1A, CDKL5, KCNT1, GRIN1, PCDH19, CHRNA4, KCNQ2, DEPDC5, KCNQ3, HCN1, CHRNA2, LGI1, SLC6A1, GNAO1, SLC2A1, GABRG2, KCNC1, DNM1, CHRNB2, KCNA2, HNRNPU, EEF1A2, STXBP1, MEF2C, GABRB3, GABRA1, SCN1B, SLC35A2, STX1B, KCNB1, PRRT2, SCN9A, SIK1, ALG13* |
| Known DEE genes | 50 | Heyne 2018[27] | Table S3 | *ALG13 ,ARHGEF9, ARX, CACNA1A, CASK, CDKL5, CHD2, DNM1, EEF1A2, FOXG1, GABRA1, GABRB3, GNAO1, GNB1, GPHN, GRIN1, GRIN2A, GRIN2B, HCN1, IQSEC2, KCNA2, KCNB1, KCNQ2, KCNT1, MBD5, MECP2, MEF2C, PCDH19, PIGA, PURA, SCN1A, SCN2A, SCN8A, SIK1, SLC2A1, SLC35A2, SLC6A1, SLC6A8, SLC9A6, SPTAN1, STXBP1, SYN1, SYNGAP1, TSC1, TSC2, UBE3A, WDR45, ZEB2, SLC1A2, GRIN2D* |
| Genes in neurodevelopmental disorders (NDDs) with epilepsy | 33 | Heyne 2018[27] | Table S6 | *KCNQ2, SCN2A, SCN1A, STXBP1, CHD2, CDKL5, DNM1, DYRK1A, MEF2C, SYNGAP1, GABRB3, EEF1A2, SLC6A1, SCN8A, PURA, WDR45, GNAO1, HNRNPU, SMC1A, FOXG1, ARID1B, GRIN2A, GRIN2B, ALG13, ASXL3, KCNH1, GABRB2, NEXMIF, MECP2, SNAP25, COL4A3BP, SLC35A2, ARHGEF9* |
| **Neurotransmission** | | | | |
| GABA-A receptor genes | 19 | May 2018[28] | Table S5 | *GABRA1, GABRA2, GABRA3, GABRA4, GABRA5, GABRA6, GABRB1, GABRB2, GABRB3, GABRD, GABRE, GABRG1, GABRG2, GABRG3, GABRP, GABRQ, GABRR1, GABRR2, GABRR3* |

| | | | | |
|---|---|---|---|---|
| GABAergic pathway genes | 113 | May 2018[28] | Table S5 | *ABAT, ADCY1, ADCY2, ADCY3, ADCY4, ADCY5, ADCY6, ADCY7, ADCY8, ADCY9, ANK2, ANK3, ARHGEF9, DISC1, DLC1, DLC2, DNAI1, FGF13, GABARAP, GABARAPL1, GABARAPL2, GABBR1, GABBR2, GABRA1, GABRA2, GABRA3, GABRA4, GABRA5, GABRA6, GABRB1, GABRB2, GABRB3, GABRD, GABRE, GABRG1, GABRG2, GABRG3, GABRP, GABRQ, GABRR1, GABRR2, GABRR3, GAD1, GAD2, GLS, GLS2, GLUL, GNAI1, GNAI2, GNAI3, GNAO1, GNB1, GNB2, GNB3, GNB4, GNB5, GNG10, GNG11, GNG12, GNG13, GNG2, GNG3, GNG4, GNG5, GNG7, GNG8, GNGT1, GNGT2, GPHN, HAP1, KCNB2, KCNC1, KCNC2, KCNC3, KCNJ6, KIF5A, KIF5B, KIF5C, MAGI, MKLN1, MYO5A, NLGN2, NRXN1, NSF, PFN1, PLCL1, PRKACA, PRKACB, PRKACG, PRKCA, PRKCB, PRKCG, RAFT1, RDX, SCN1A, SCN1B, SCN2B, SCN3A, SCN8A, SEMA4D, SLC12A2, SLC12A5, SLC32A1, SLC38A1, SLC38A2, SLC38A3, SLC38A5, SLC6A1,SLC6A11, SLC6A13, SRC, TRAK1, TRAK2* |
| Excitatory receptor genes (Glutamate ionotropic receptors & cholinergic receptors) | 34 | May 2018[28] | Table S5 | *CHRNA1, CHRNA10, CHRNA2, CHRNA3, CHRNA4, CHRNA5, CHRNA6, CHRNA7, CHRNA9, CHRNB1, CHRNB2, CHRNB3, CHRNB4, CHRND, CHRNE, CHRNG, GRIA1, GRIA2, GRIA3, GRIA4, GRIK1, GRIK2, GRIK3, GRIK4, GRIK5, GRIN1, GRIN2A, GRIN2B, GRIN2C, GRIN2D, GRIN3A, GRIN3B, GRID1, GRID2* |
| Voltage-gated cation channel genes | 86 | May 2018[28] | Table S5 | *SCN10A, SCN11A, SCN1A, SCN1B, SCN2A2, SCN2B, SCN3A, SCN3B, SCN4A, SCN4B, SCN5A, SCN7A, SCN8A, SCN9A, CACNA1A, CACNA1B, CACNA1C, CACNA1D, CACNA1E, CACNA1F, CACNA1G, CACNA1H, CACNA1I, CACNA1S, CACNA2D1, CACNA2D2, CACNA2D3, CACNA2D4, CACNB1, CACNB2, CACNB3, CACNB4, KCNA1, KCNA10, KCNA2, KCNA3, KCNA4, KCNA5, KCNA6,KCNA7, KCNAB1, KCNAB2, KCNAB3, KCNB1, KCNB2, KCNC1, KCNC2, KCNC3, KCNC4, KCND1, KCND2, KCND3, KCNE1, KCNE1L, KCNE2, KCNE3, KCNE4, KCNF1, KCNG1, KCNG2, KCNG3, KCNG4, KCNH1, KCNH2, KCNH3, KCNH4, KCNH5, KCNH6, KCNH7, KCNH8, KCNQ1, KCNQ2, KCNQ3, KCNQ5, KCNQ4, KCNRG, KCNS1, KCNS2, KCNS3, KCNT1, KCNV1, KCNV2, HCN1, HCN2, HCN3, HCN4* |

**Table S7.** Prior gene panel screening of the 1,021 DEE-affected individuals
More than 50 different gene panels were used across our patient cohorts. This table summarizes the overall screening results, but heterogeneity exists in the numbers and the types of genes tested, how the testing was performed, etc.

| Gene panel screening | Results | | |
|---|---|---|---|
| | Abnormal or unknown | Normal | Total |
| Yes | 89 | 257 | 346 |
| No | - | - | 247 |
| Not entered/answered | - | - | 428 |
